# Discovery and Optimization of Indoline-2-Carboxamide-Based FBXO31 Recruiters for Targeted Protein Degradation

**DOI:** 10.64898/2026.11.23.768409

**Authors:** Xiaokang Jin, Chenlu Zhang, Yunfeng Li, Chen Zhou, Daria Spasskaya, Daniel Banas, Bryan W. Dorsey, Kyle Mangano, Bradley Brasher, Michele Pagano, Bing Hao, Xiaoyu Zhang

## Abstract

Expanding the repertoire of chemically recruitable E3 ligases is critical for advancing targeted protein degradation (TPD). FBXO31 has recently emerged as a promising E3 ligase for TPD, but existing peptide-derived recruiters exhibit limited degradation potency and efficacy at endogenous FBXO31 expression levels. Here, we report the discovery and optimization of indoline-2-carboxamide-based FBXO31 recruiters through structure-activity relationship studies. Optimization of the recruiter scaffold and linker yielded JZ26, a highly potent degrader that induced near-complete FKBP12 degradation at endogenous FBXO31 expression levels (DC_50_ = 2.5 nM, D_max_ = 91%). JZ26 also induced potent FKBP12 degradation in primary mouse fibroblasts expressing FBXO31. Furthermore, the optimized recruiter was incorporated into bifunctional degraders targeting BRD9 and SMARCA2, demonstrating its applicability to multiple protein targets.

## Introduction

Targeted protein degradation (TPD) has emerged as a powerful strategy for modulating protein function by harnessing the cellular ubiquitin-proteasome system^1–4^. Bifunctional degraders, such as proteolysis-targeting chimeras (PROTACs), induce proximity between a protein of interest and an E3 ubiquitin ligase, leading to target ubiquitination and subsequent proteasomal degradation^5–8^. Although the human genome encodes more than 600 E3 ligases, only a small fraction have been successfully exploited for TPD, with cereblon (CRBN) and von Hippel-Lindau (VHL) remaining the most widely used^9^. Expanding the repertoire of chemically recruitable E3 ligases is therefore important for overcoming the limitations of existing degradation platforms and enabling new therapeutic applications^10–12^.

FBXO31 is an F-box protein that functions as a substrate-recognition component of the SKP1-CUL1-F-box (SCF) E3 ubiquitin ligase complex. It recognizes proteins bearing C-terminal amides and targets them for ubiquitination and proteasomal degradation^13^. In our previous study, we identified a peptide-derived chemical motif containing a C-terminal primary amide that enables FBXO31 recruitment and demonstrated its applicability to TPD^14^. Despite this initial discovery, the peptide-derived nature of the initial FBXO31 recruiters presents potential challenges for further development, including susceptibility to metabolic degradation and limited cellular permeability.

Moreover, the initial FBXO31-recruiting degraders exhibited limited degradation potency and efficacy in cells expressing endogenous FBXO31. These limitations underscore the need to develop more potent and structurally simplified FBXO31 recruiters to facilitate their broader application in TPD. Here, we report the discovery and optimization of indoline-2-carboxamide-based FBXO31 recruiters through systematic structure-activity relationship (SAR) studies. The optimized recruiters enable potent protein degradation at endogenous FBXO31 expression levels and can be extended to additional protein targets, including BRD9 and SMARCA2.

## Results

### Structure-activity relationship studies identify optimized non-natural peptide-derived FBXO31 recruiters

In our previous study, we identified an Ala-Phe-CONH_2_-derived chemical motif capable of recruiting the E3 ligase FBXO31 to induce targeted protein degradation^14^ (**Figure 1A**). To further improve degradation potency and efficacy, we conducted systematic SAR studies by modifying individual structural elements of the original dipeptide scaffold. For this optimization campaign, we conjugated the FBXO31-recruiting motifs to the FKBP12-binding ligand SLF to generate bifunctional degraders and evaluated their activity using the same cellular system as in our previous study, in which HEK293T cells expressed endogenously HiBiT-tagged FKBP12 and overexpressed FBXO31^14^.

**Figure 1.**
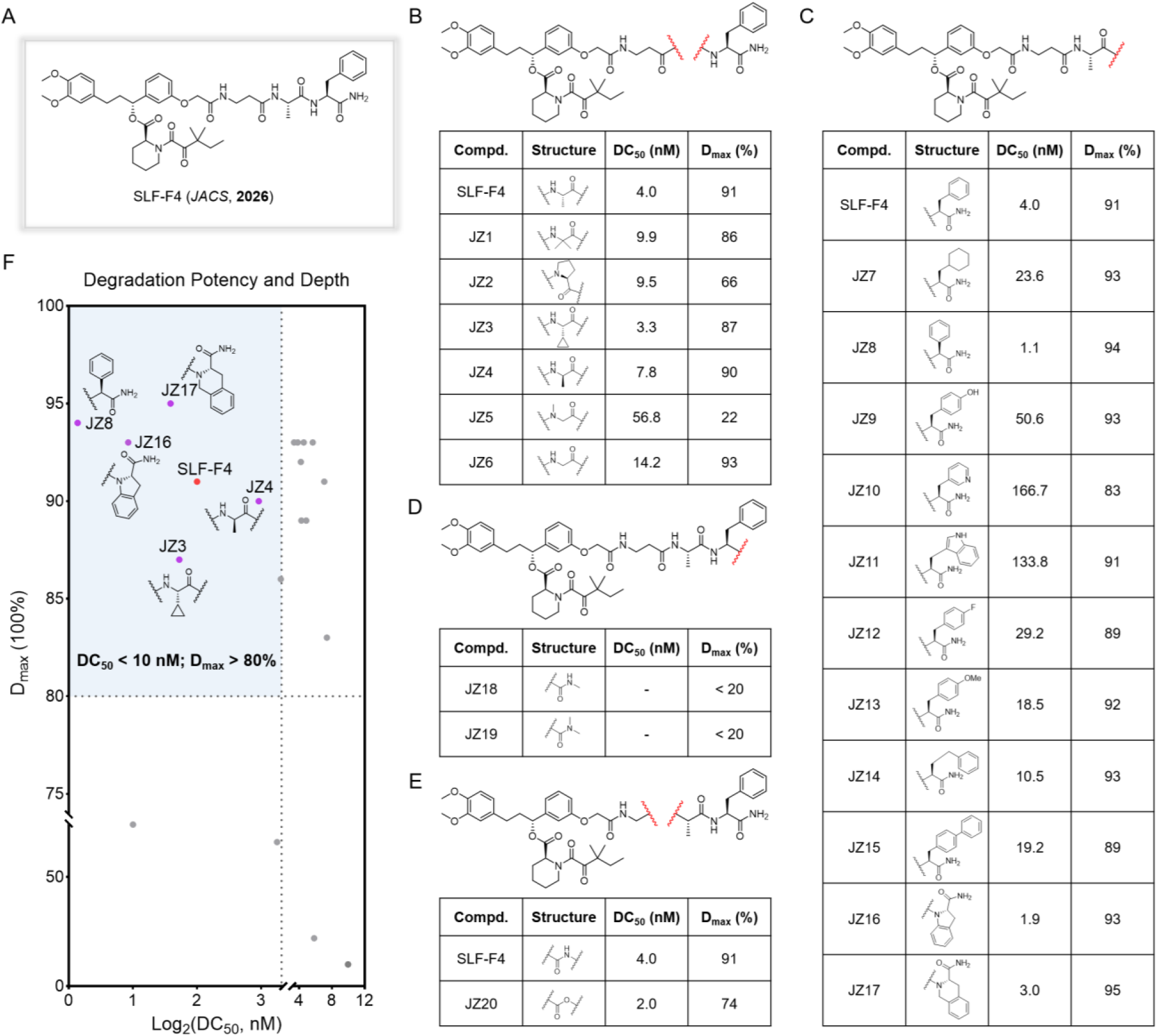
Structure-activity relationship studies of FBXO31 recruiters. **A**. Chemical structure of the original FBXO31-recruiting degrader SLF-F4. **B**. SAR studies examining modifications to the alanine residue. The table summarizes the DC_50_ and D_max_ values following 24 h of treatment. **C**. SAR studies examining modifications to the phenylalanine residue. The table summarizes the DC_50_ and D_max_ values following 24 h of treatment. **D**. Evaluation of N-methylation of the C-terminal primary amide. The table summarizes the DC_50_ and D_max_ values following 24 h of treatment. **E**. Evaluation of amide-to-ester replacement within the linker. The table summarizes the DC_50_ and D_max_ values following 24 h of treatment. **F**. Summary of degradation potency (DC_50_) and maximal degradation efficacy (D_max_) for the evaluated analogs.

We first investigated the contribution of the alanine residue by modifying its side chain while retaining the phenylalanine residue and C-terminal primary amide (**Figure 1B** and **S1**). Replacement of alanine with other amino acid residues, including those bearing larger hydrophobic or polar side chains, generally maintained degradation activity comparable to or slightly lower than that of the parent compound. An exception was JZ5, which contains an N-methylated glycine residue and exhibited substantially reduced degradation activity, with a D_max_ (maximal degradation efficacy) of only 22%.These results suggest that modifications at this position may offer limited opportunities for improving degradation potency and efficacy.

We next explored modifications to the C-terminal phenylalanine residue (**Figure 1C** and **S1**). Replacement of the phenyl group with a cyclohexyl group (JZ7) reduced degradation potency, whereas removal of the methylene group between the phenyl ring and the α-carbon (JZ8) improved potency relative to SLF-F4. Introduction of a para-hydroxyl group onto the phenyl ring (JZ9) or replacement of the phenyl group with a pyridyl (JZ10) or indolyl (JZ11) group substantially reduced degradation potency, with DC_50_ values ranging from 50.6 to 166.7 nM. Similarly, introduction of a para-fluoro (JZ12) or para-methoxy (JZ13) substituent reduced degradation potency, with DC_50_ values of 29.2 and 18.5 nM, respectively. Introduction of an additional methylene group between the phenyl ring and the α-carbon (JZ14) or incorporation of a biphenyl group (JZ15) also reduced potency, with DC_50_ values of 10.5 and 19.2 nM, respectively.

Importantly, incorporation of an indoline-2-carboxamide motif (JZ16) improved degradation potency approximately twofold relative to SLF-F4 (DC_50_ = 1.9 versus 4.0 nM), while maintaining high maximal degradation efficacy (D_max_ = 93% versus 91%). The related tetrahydroisoquinoline-3-carboxamide analog (JZ17), containing a six-membered nitrogen-containing ring, also exhibited potent degradation activity (D_max_ = 3.0 nM, D_max_ = 95%). Taken together, these findings identify the modified phenylalanine-derived scaffold (JZ8) and the conformationally constrained indoline-2-carboxamide (JZ16) and tetrahydroisoquinoline-3-carboxamide (JZ17) motifs as promising alternatives to the original FBXO31 recruiter for further optimization.

We also investigated the C-terminal primary amide and the peptide backbone. N-monomethylation (JZ18) or N,N-dimethylation (JZ19) of the C-terminal primary amide abolished degradation activity, suggesting an essential role for this functional group in FBXO31-mediated degradation (**Figure 1D** and **S1**). In addition, replacement of a single amide bond in the linker with an ester (JZ20) maintained comparable degradation potency (DC_50_ = 2.0 versus 4.0 nM) but reduced degradation depth (D_max_ = 74% versus 91%) (**Figure 1E** and **S1**), suggesting that this amide linkage is important for the overall degradation efficacy. To summarize these SAR results, we compared the degradation potency (DC_50_) and maximal degradation efficiency (D_max_) of all evaluated analogs in a single plot (**Figure 1F**).

### Optimization of indoline- and tetrahydroisoquinoline-based FBXO31 recruiters

Among the evaluated analogs, JZ16 and the structurally related JZ17 both exhibited potent degradation activity. We therefore selected these two conformationally constrained, non-natural peptide-derived scaffolds for further optimization. We first evaluated JZ16 in an in vitro ubiquitination assay and observed the formation of ubiquitinated FKBP12 species (**Figure S2**), providing additional evidence that JZ16 supports FBXO31-mediated substrate ubiquitination. Notably, inclusion of the priming E3 ligase ARIH1 and its cognate E2, UBE2L3, increased FKBP12 ubiquitylation, consistent with observations reported for CRL1-FBXO22^15^. We next assessed the contribution of the alanine residue within the indoline-2-carboxamide-based JZ16 scaffold (**Figure 2A** and **2B**). Replacement of alanine with glycine (JZ21) substantially improved degradation potency, reducing the DC_50_ by more than 20-fold while maintaining high degradation efficacy (D_max_ = 96%). Incorporation of an isopropyl side chain (JZ22) also improved potency, although to a lesser extent, with a twofold reduction in DC_50_. In contrast, introduction of other bulkier hydrophobic side chains (JZ23 and JZ24) markedly reduced degradation potency, with DC_50_ values of 27.9 and 36.0 nM, respectively. Incorporation of proline (JZ25) maintained degradation activity comparable to that of JZ16 (DC_50_ = 1.2 nM, D_max_ = 95%). These results suggest that smaller amino acid residues are generally favored at this position, with glycine showing the greatest improvement in degradation potency.

**Figure 2.**
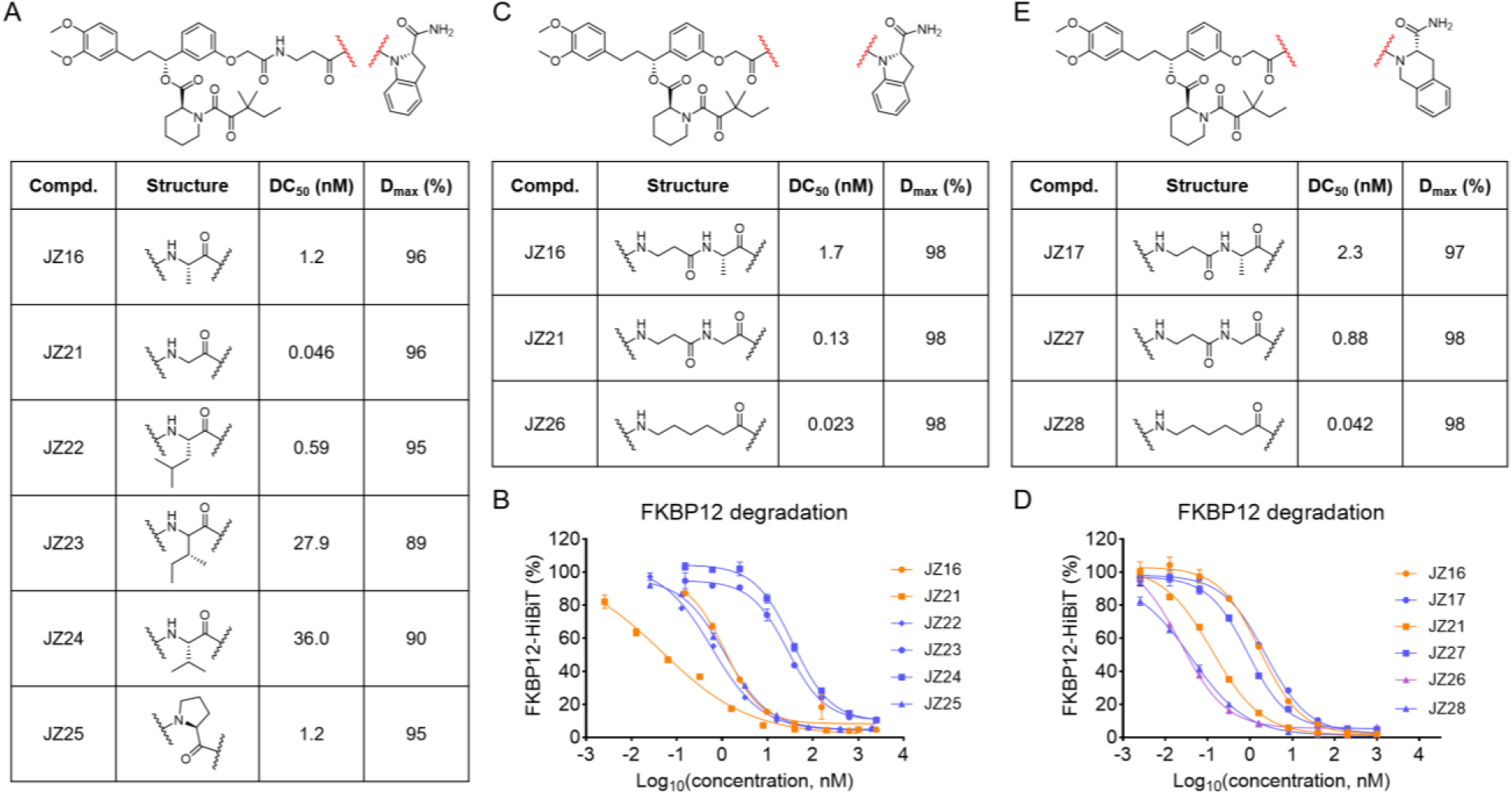
Optimization of indoline- and tetrahydroisoquinoline-based FBXO31 recruiters. **A**. Chemical structures of JZ16 analogs with modifications to the alanine residue and their corresponding DC_50_ and D_max_ values following 24 h of treatment. **B**. Concentration-dependent degradation of FKBP12-HiBiT in HEK293T cells treated with JZ16 and its analogs for 24 h. Data are presented as mean ± SEM (n = 3). **C**. Chemical structures of JZ16, JZ21, and JZ26 and their corresponding DC_50_ and D_max_ values following 24 h of treatment. **D**. Concentration-dependent degradation of FKBP12-HiBiT in HEK293T cells treated with JZ16 and its analogs for 24 h. Data are presented as mean ± SEM (n = 3). **E**. Chemical structures of JZ17, JZ27, and JZ28 and their corresponding DC_50_ and D_max_ values following 24 h of treatment.

Given that reducing the size of the amino acid side chain improved degradation potency (JZ21 versus JZ16), we wonder whether further simplification by replacing the amide-containing linker with an all-carbon chain of comparable length might further enhance degradation activity. To test this, we synthesized JZ26, in which the amide-containing linker was replaced with an alkyl chain. JZ26 exhibited substantially improved degradation potency, with the DC_50_ of 0.023 nM and the D_max_ of 98% (**Figure 2C** and **2D**). We next examined whether this linker optimization strategy could be extended to the tetrahydroisoquinoline-3-carboxamide-based JZ17 scaffold. Consistent with our findings for the JZ16 scaffold, replacement of the alanine residue with glycine (JZ27) improved degradation potency relative to JZ17 (DC_50_ = 2.3 versus 0.88 nM), while maintaining high degradation efficacy (D_max_ = 98%) (**Figure 2D** and **2E**). Further replacement of the amide-containing linker with an all-carbon linker (JZ28) substantially improved degradation potency, with the DC_50_ of 0.042 nM and the D_max_ of 98% (**Figure 2D** and **2E**).

### Characterization and mechanistic validation of optimized FBXO31-recruiting degraders

Having identified JZ26 and JZ28 as highly potent FBXO31-recruiting degraders, we next sought to further characterize their degradation activity. Given that FBXO31 is a relatively low-abundance E3 ligase in many cell types, our initial SAR studies were conducted in an FBXO31-overexpressing system. To determine whether the optimized degraders could also function at endogenous FBXO31 expression levels, we evaluated JZ16, JZ17, JZ26, and JZ28 in HEK293T parental cells expressing endogenous FBXO31 (**Figure 3A**). The original degrader SLF-F4 exhibited substantially reduced degradation efficacy in this system, with the D_max_ of only 45%. JZ16 and JZ17 also exhibited limited degradation depth, with D_max_ values of 61% and 64%, respectively (**Figure 3A** and **S3**). In contrast, the optimized degraders JZ26 and JZ28 maintained potent and near-complete degradation activity. JZ26 achieved the DC_50_ of 2.5 nM and the D_max_ of 91%, whereas JZ28 exhibited the DC_50_ of 2.8 nM and the D_max_ of 90% (**Figure 3A** and **S3**).

**Figure 3.**
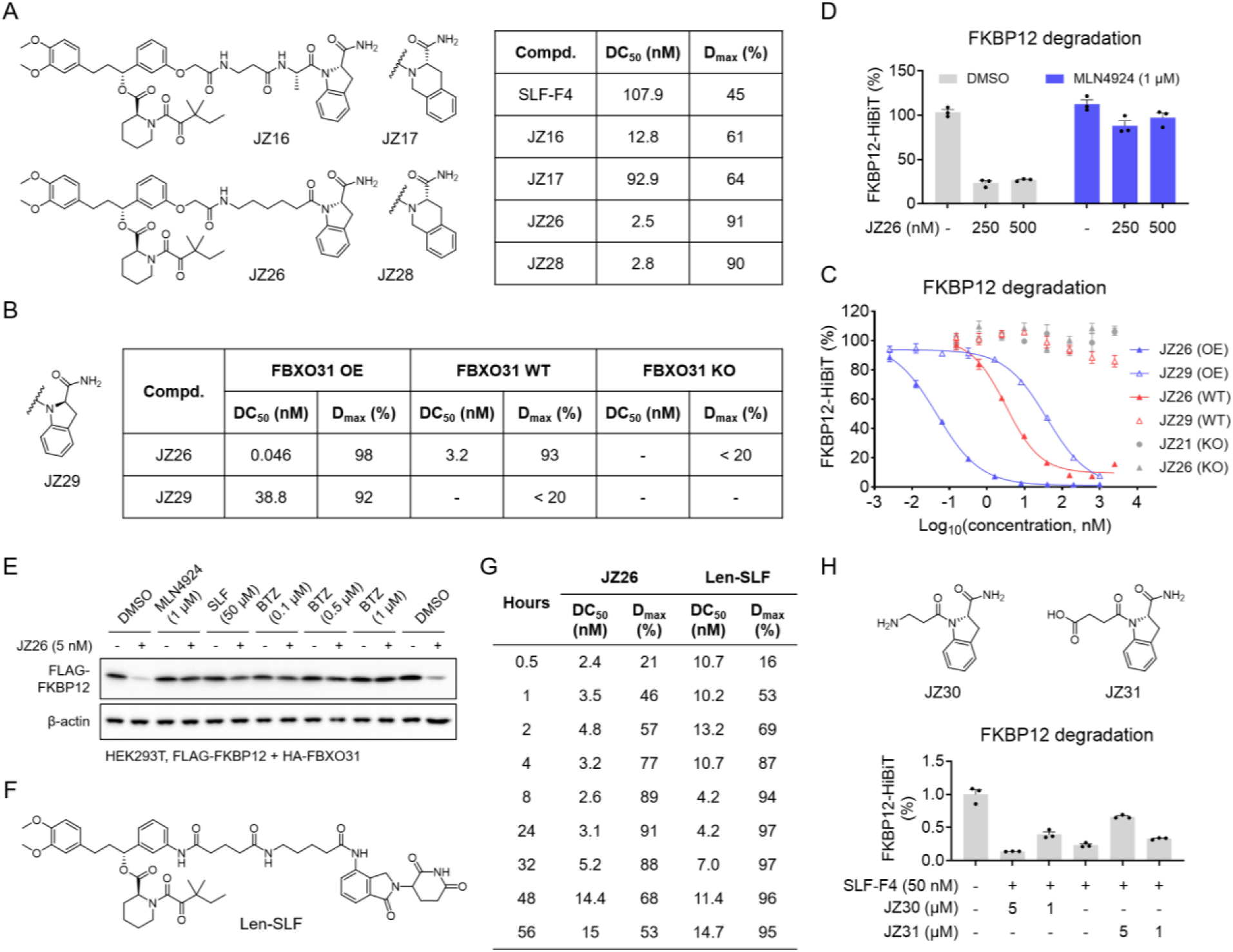
Characterization and mechanistic validation of optimized FBXO31-recruiting degraders. **A**. DC_50_ and D_max_ values for FKBP12 degradation induced by SLF-F4, JZ16, JZ17, JZ26, and JZ28 in HEK293T cells expressing endogenous FBXO31. **B**. Chemical structure of JZ29 and DC_50_ and D_max_ values for FKBP12 degradation induced by JZ26 and JZ29 following 24 h treatment in FBXO31-overexpressing, parental, and FBXO31-knockout HEK293T cells. **C**. Concentration-dependent FKBP12 degradation induced by JZ26 and JZ29 in FBXO31-overexpressing, parental, and FBXO31-knockout HEK293T cells following 24 h treatment. Data are presented as mean ± SEM (n = 3). **D**. Effect of MLN4924 on JZ26-induced FKBP12 degradation in HEK293T cells expressing endogenous FBXO31 following 1 h pretreatment with MLN4924 and subsequent 6 h co-treatment with JZ26. Data are presented as mean ± SEM (n = 3). **E**. Western blot analysis of JZ26-induced FLAG-FKBP12 degradation in HEK293T cells expressing HA-FBXO31, with or without 1 h pretreatment with bortezomib (BTZ), MLN4924, or SLF, followed by 6 h co-treatment with JZ26. **F.** Chemical structure of the CRBN-recruiting degrader Len-SLF. **G**. DC_50_ and D_max_ values for FKBP12 degradation induced by JZ26 or Len-SLF in HEK293T cells expressing endogenous FBXO31 at different treatment time points. **H**. Chemical structures of JZ30 and JZ31 and their inhibitory effects on SLF-F4-induced FKBP12 degradation following 1 h pretreatment with JZ30 or JZ31 and subsequent 8 h co-treatment with SLF-F4. Data are presented as mean ± SEM (n = 3).

We next examined the stereochemistry within the indoline-2-carboxamide motif by synthesizing JZ29 (*R*), a diastereomer of JZ26 (*S*) with the opposite configuration at the indoline stereocenter (**Figure 3B**). Although JZ29 retained degradation activity in FBXO31-overexpressing cells, its potency was > 840-fold lower than that of JZ26 (DC_50_ = 38.8 versus 0.046 nM) (**Figure 3B** and **3C**). Importantly, JZ29 failed to induce FKBP12 degradation in parental cells expressing endogenous FBXO31, whereas JZ26 maintained potent degradation activity (DC_50_ = 3.2 nM, D_max_ = 93%) (**Figure 3B** and **3C**). Furthermore, JZ26-induced degradation was abolished in FBXO31-knockout cells, confirming its dependence on FBXO31. These results demonstrate the essentiality of the stereochemical configuration of the indoline-2-carboxamide motif for efficient FBXO31-mediated degradation at endogenous expression levels.

To further investigate the mechanism of degradation induced by JZ26, the most potent degrader, we examined the effect of MLN4924, an inhibitor of cullin neddylation that suppresses Cullin-RING ligase (CRL) activity^16^. Treatment with MLN4924 blocked JZ26-induced FKBP12 degradation in cells expressing endogenous FBXO31 (**Figure 3D**), consistent with the involvement of the CRL machinery in FBXO31-mediated protein degradation. We further validated JZ26 activity using an orthogonal degradation system in HEK293T cells expressing FLAG-tagged FKBP12 and HA-tagged FBXO31. Western blot analysis confirmed that JZ26 induced robust degradation of FLAG-FKBP12, which was blocked by co-treatment with the proteasome inhibitor bortezomib (BTZ)^17^, MLN4924, or the FKBP12-binding ligand SLF (**Figure 3E**). We next compared the degradation kinetics of JZ26 with Len-SLF, an established cereblon (CRBN)-recruiting FKBP12 degrader^18^ (**Figure 3F**). Time-course experiments revealed that JZ26 induced FKBP12 degradation rapidly, achieving the D_max_ of 57% within 2 h and 77% after 4 h, with maximal degradation reaching 91% at 24 h (**Figure 3G** and **S3**). During the first 32 h, JZ26 exhibited greater degradation potency than Len-SLF, as indicated by its lower DC_50_ values (**Figure 3G** and **S3**). At later time points, Len-SLF maintained high degradation efficacy through 56 h, whereas JZ26 exhibited a gradual reduction in degradation activity after 32 h. These results indicate that JZ26 induces rapid and efficient FKBP12 degradation, although its activity is less sustained than that of the CRBN-based degrader under the conditions tested.

Finally, to assess whether the optimized indoline-2-carboxamide motif engages the same FBXO31-binding site as the original peptide-derived recruiter, we synthesized two indoline-2-carboxamide derivatives bearing either a terminal primary amine (JZ30) or a carboxylic acid (JZ31) (**Figure 3H**). Both compounds inhibited FKBP12 degradation induced by the original FBXO31-recruiting degrader SLF-F4 (**Figure 3H**), suggesting that these indoline-derived compounds likely engage the same FBXO31-binding pocket as the original peptide-based recruiter.

### Structural modeling and proteomic characterization of JZ26-induced ternary complex formation

To further understand the molecular basis of JZ26-mediated FBXO31 recruitment, we employed Boltz-2 to model the ternary complex formed between FKBP12, JZ26, and FBXO31. This approach was previously shown to generate high-confidence structural predictions for the FBXO31-bifunctional degrader-FKBP12 ternary complex^14^. The predicted structure positioned JZ26 at the interface between FKBP12 and FBXO31, supporting the formation of a productive ternary complex (**Figure 4A** and **S4**). Four FBXO31 residues (Y309, D334, I337, and T343), previously identified as important for recruiter recognition, were located within the JZ26-binding pocket (**Figure 4A**). To validate their contributions to JZ26-mediated degradation, we re-expressed the corresponding FBXO31 mutants (Y309A, D334N, I337D, and T343V) in FBXO31-knockout cells. Whereas re-expression of wild-type FBXO31 restored JZ26-induced FKBP12 degradation, none of these mutants rescued degradation activity (**Figure 4B**). These results indicate that the optimized recruiter retains a recognition mechanism similar to that of the original peptide-derived scaffold.

**Figure 4.**
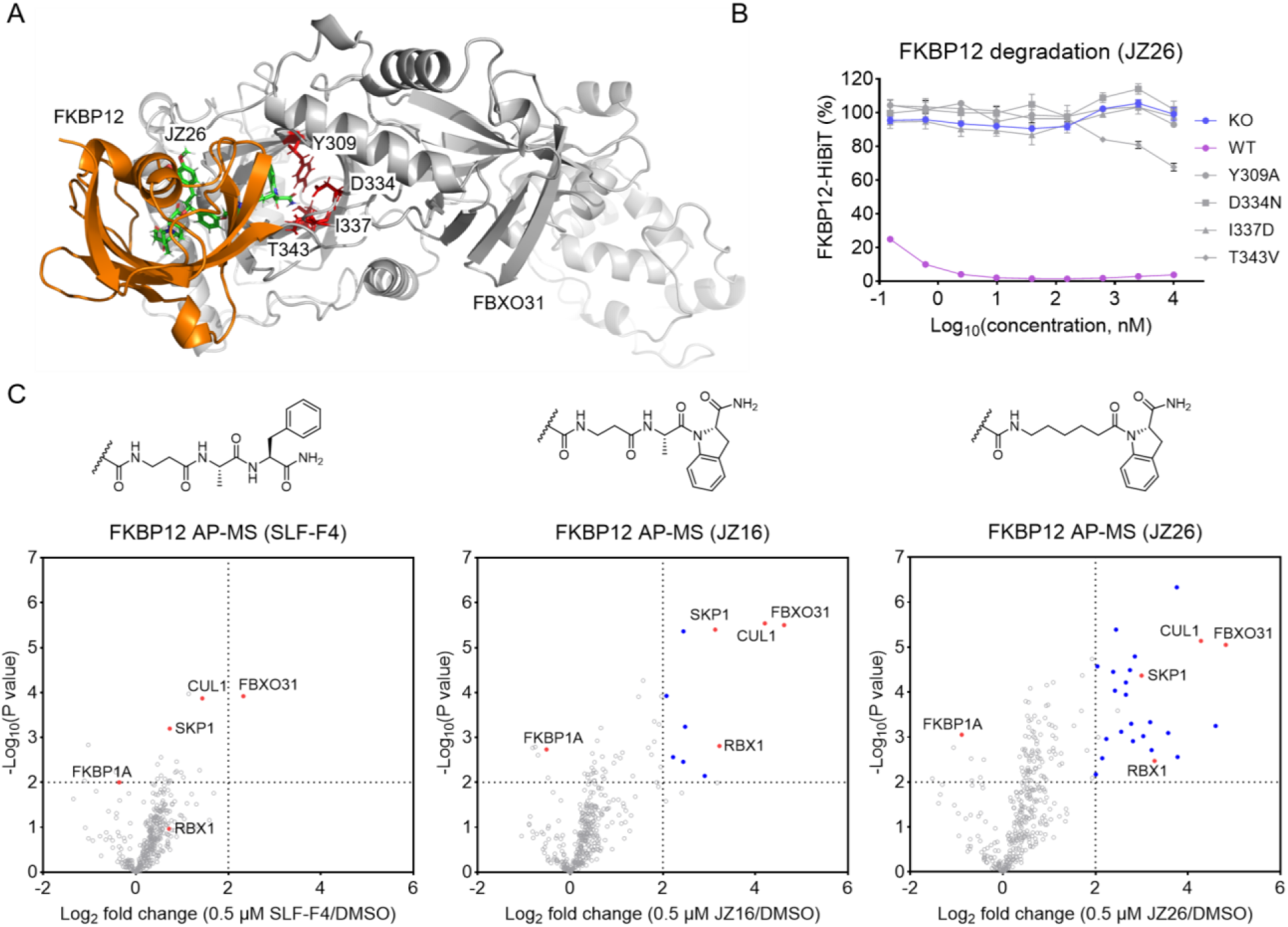
Structural modeling and proteomic characterization of JZ26-induced ternary complex formation. **A**. Boltz-2-predicted ternary complex structure of FKBP12, JZ26, and FBXO31, showing the JZ26-binding pocket and key FBXO31 residues (Y309, D334, I337, and T343). **B**. Concentration-dependent FKBP12 degradation induced by JZ26 in FBXO31-knockout HEK293T cells re-expressing wild-type FBXO31 or its mutants (Y309A, D334N, I337D, and T343V) following 24 h treatment. Data are presented as mean ± SEM (n = 3). **C**. Volcano plots of AP-MS analysis using FKBP12 as the bait in cells treated with SLF-F4, JZ16, or JZ26. Cells were pretreated with 100 nM BTZ for 1 h, followed by treatment with DMSO or 0.5 µM SLF-F4, JZ16, or JZ26 for 3 h. *P* values were calculated using a two-sided t-test and adjusted for multiple comparisons by the Benjamini-Hochberg method (n = 4 for DMSO; n = 3 for SLF-F4, JZ16, or JZ26). FBXO31 and components of the SCF ubiquitin ligase complex (SKP1, CUL1, and RBX1) are indicated. Proteasomal subunits are highlighted as blue dots.

We next employed affinity purification coupled with mass spectrometry (AP-MS) to assess JZ26-induced protein complex formation, using FKBP12 as the bait. We compared the original degrader SLF-F4, the initial indoline-2-carboxamide-based degrader JZ16, and the optimized degrader JZ26 under identical experimental conditions. All three compounds promoted enrichment of FBXO31 and components of the SCF ubiquitin ligase complex, including SKP1, CUL1, and RBX1 (**Figure 4C** and **Table S1**). In particular, JZ16 induced greater enrichment of the CRL complex than SLF-F4, while JZ26 induced even greater enrichment than JZ16. This trend is consistent with the relative degradation activities of the three compounds. Interestingly,

JZ26 treatment also led to the enrichment of multiple proteasomal subunits (blue dots in **Figure 4C**), which was not observed with other degraders or E3 ligases in our previous studies. This suggests that JZ26 promotes the association of FKBP12 not only with the FBXO31-containing ubiquitin ligase complex but also with components of the proteasomal machinery.

### Mouse FBXO31 supports potent FKBP12 degradation by optimized FBXO31-recruiting degraders

To investigate whether the FBXO31-recruiting degraders could also function with mouse FBXO31, we expressed mouse FBXO31 in HEK293T FBXO31-knockout cells and evaluated FKBP12-HiBiT degradation induced by SLF-F4 and JZ26. Both compounds induced potent FKBP12 degradation in cells expressing mouse FBXO31, whereas JZ26 failed to induce degradation in FBXO31-knockout cells (**Figure 5A**). As observed with human FBXO31, JZ26 exhibited substantially greater degradation potency than SLF-F4, with the DC_50_ of 0.033 nM compared with 2.9 nM, while both compounds achieved near-complete degradation (D_max_ = 99% and 98%, respectively) (**Figure 5B**). We further performed structural modeling of human and mouse FBXO31 in complex with the indoline-derived analog JZ31 (**Figure 5C**). Superposition of the predicted structures revealed a high degree of structural similarity within the recruiter-binding pocket, including the conserved positioning of four key residues previously implicated in recruiter recognition (Y309/Y295, D334/D320, I337/I323, and T343/T329 in human/mouse FBXO31, respectively) (**Figure 5C**). The predicted binding configurations of JZ31 were also highly similar in the two protein complexes, suggesting a conserved mode of recruiter recognition.

**Figure 5.**
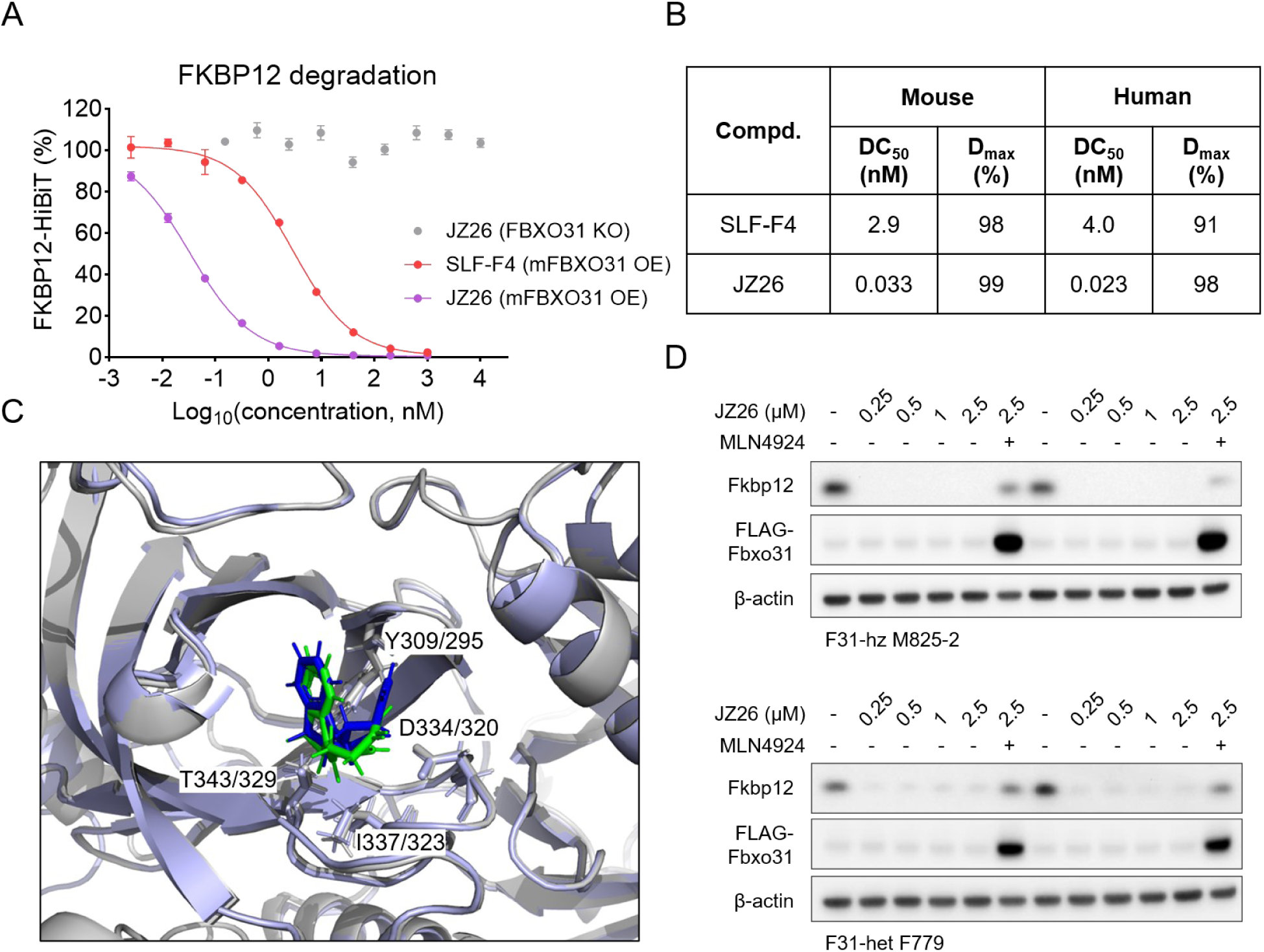
Mouse FBXO31 supports FKBP12 degradation by FBXO31-recruiting degraders. **A**. Concentration-dependent FKBP12 degradation induced by SLF-F4 and JZ26 in FBXO31-knockout HEK293T cells with or without re-expression of mouse FBXO31 following 24 h treatment. Data are presented as mean ± SEM (n = 3). **B**. DC_50_ and D_max_ values for FKBP12 degradation induced by SLF-F4 and JZ26 in HEK293T cells expressing human or mouse FBXO31. **C**. Superposition of Boltz-2-predicted structures of human and mouse FBXO31 in complex with JZ31. **D**. Western blot analysis of JZ26-induced endogenous mouse Fkbp12 degradation in primary mouse fibroblasts from two transgenic mouse lines overexpressing FLAG-tagged Fbxo31 (F31-hz M825-2 and F31-het F779), with or without co-treatment with MLN4924 for 24 h.

To further validate the activity of JZ26 in mouse cells, we evaluated the degradation of endogenous Fkbp12 in primary mouse fibroblasts isolated from two transgenic mouse lines overexpressing FLAG-tagged Fbxo31 (F31-hz M825-2 and F31-het F779). JZ26 induced concentration-dependent degradation of endogenous Fkbp12 in both cell lines (**Figure 5D**). Further evaluation using lower compound concentrations revealed potent degradation activity, with estimated DC_50_ values of 0.5-1 nM in F31-hz M825-2 cells and about 1 nM or slightly higher in F31-het F779 cells (**Figure S5**). Co-treatment with MLN4924 blocked JZ26-induced degradation (**Figure 5D**), supporting a cullin-RING ligase-dependent mechanism in mouse cells.

### Development of FBXO31-recruiting degraders targeting BRD9 and SMARCA2

We next investigated whether the indoline-2-carboxamide scaffold could be extended to the degradation of additional protein targets. We first developed a series of FBXO31-recruiting degraders targeting bromodomain-containing protein 9 (BRD9) by conjugating the optimized recruiter to a BRD9-binding ligand^19,20^ through alkyl linkers of varying lengths (JZ32, JZ33, and JZ34) (**Figure 6A**). All three degraders induced robust BRD9 degradation in FBXO31-overexpressing HEK293T cells, with the JZ34 analog exhibiting the greatest degradation potency. JZ34 induced substantial BRD9 degradation at subnanomolar concentrations, with the DC_50_ of < 0.05 nM and the D_max_ > 95% (**Figure 6A**).

**Figure 6.**
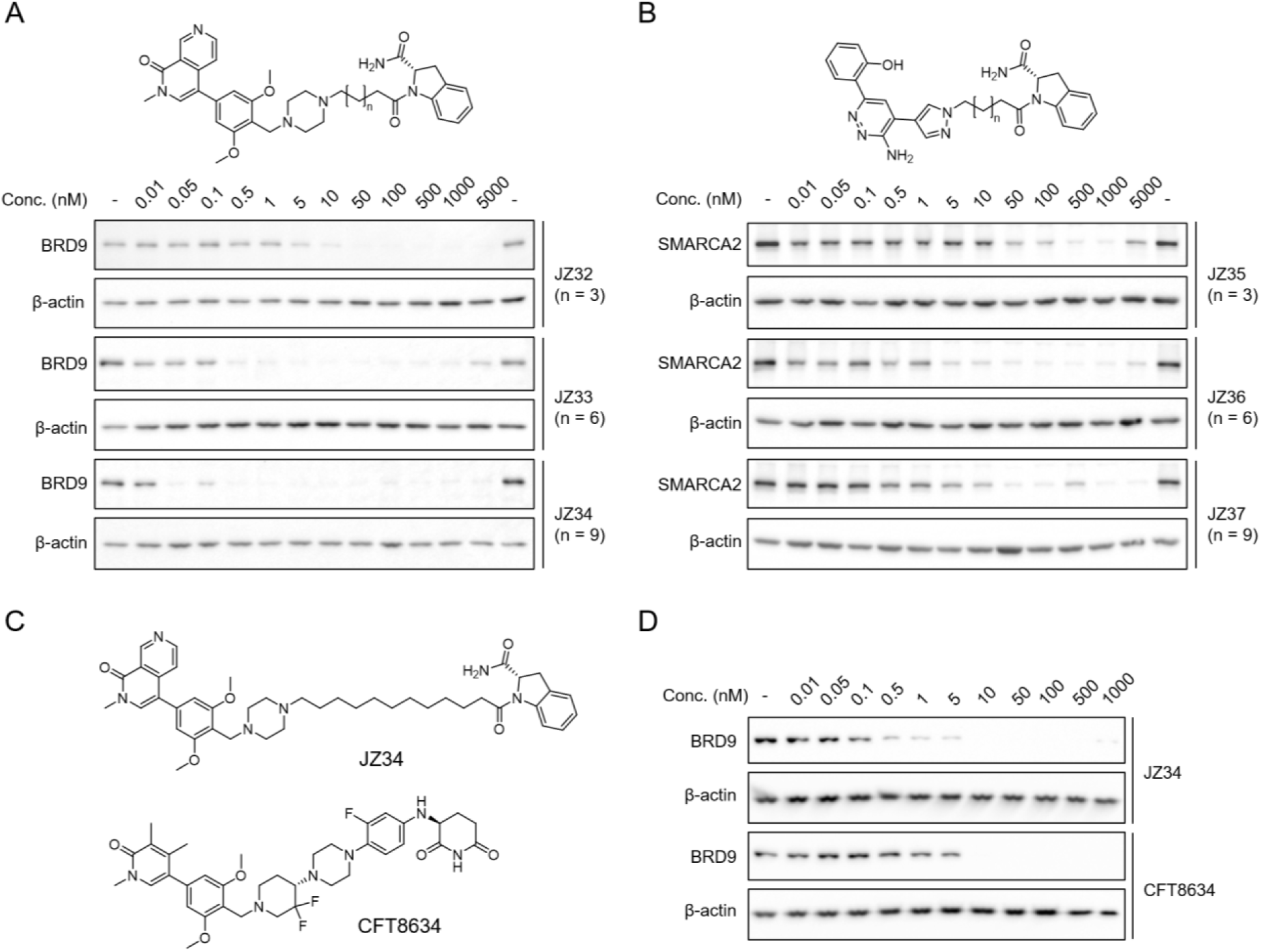
Development of FBXO31-recruiting degraders targeting BRD9 and SMARCA2. **A**. Chemical structures of FBXO31-recruiting BRD9 degraders with varying alkyl linker lengths (JZ32, JZ33, and JZ34) and their degradation activities in FBXO31-overexpressing HEK293T cells following 24 h of treatment. **B**. Chemical structures of FBXO31-recruiting SMARCA2 degraders with varying alkyl linker lengths (JZ35, JZ36, and JZ37) and their degradation activities in FBXO31-overexpressing HEK293T cells following 24 h of treatment. **C**. Chemical structures of JZ34 and the CRBN-recruiting BRD9 degrader CFT8634. **D**. Western blot analysis of concentration-dependent BRD9 degradation induced by JZ34 and CFT8634 in HEK293T cells expressing endogenous FBXO31 following 24 h of treatment.

We then applied a similar strategy to develop three FBXO31-recruiting bifunctional degraders targeting SMARCA2^21^, an ATP-dependent chromatin-remodeling protein, with C5, C8, or C11 alkyl linkers (JZ35, JZ36 and JZ37, **Figure 6B**). All three compounds induced concentration-dependent SMARCA2 degradation, with JZ37 exhibiting the greatest degradation potency (**Figure 6B**). Although SMARCA2 degradation was generally less efficient than BRD9 degradation, these findings demonstrate that the optimized FBXO31 recruiter can be incorporated into bifunctional degraders targeting structurally and functionally distinct proteins.

We next evaluated the lead BRD9-C11 degrader in HEK293T cells expressing endogenous FBXO31. We confirmed that JZ34 induced concentration-dependent BRD9 degradation, and this degradation was completely abolished in FBXO31-knockout cells (**Figure S6**). We further compared its degradation activity with that of CFT8634, a previously developed CRBN-recruiting BRD9 degrader that has undergone clinical investigation^22^ (**Figure 6C**). Remarkably, JZ34 induced robust BRD9 degradation at low nanomolar concentrations and exhibited greater degradation potency than CFT8634 under the same experimental conditions (**Figure 6D**). These results demonstrate the broader applicability of the optimized FBXO31 recruiter and its potential as an alternative E3 ligase recruitment strategy for targeted protein degradation.

## Discussion

In this study, we report the discovery and optimization of indoline-2-carboxamide-based FBXO31 recruiters that enable potent targeted protein degradation. Our SAR studies also identified tetrahydroisoquinoline-3-carboxamide, a related six-membered ring scaffold, as another promising motif for FBXO31 recruitment. Compared with the original peptide-derived FBXO31 recruiters, these conformationally constrained scaffolds offer opportunities for developing non-peptidic recruiters with reduced susceptibility to proteolytic cleavage and potentially improved metabolic stability.

Importantly, the optimized degrader JZ26 induced near-complete FKBP12 degradation at endogenous FBXO31 expression levels, which overcomes a major limitation of the original recruiter. Furthermore, the successful development of BRD9- and SMARCA2-targeting degraders demonstrates the broader applicability of this chemical scaffold to different protein targets. An additional advantage of the indoline-2-carboxamide scaffold is its relatively small molecular size. Although our current study focused on bifunctional degraders, the compact structure of this recruiter may facilitate the development of alternative TPD strategies, including FBXO31-recruiting molecular glues. Further structural and mechanistic studies will be important for determining whether this scaffold can support molecular glue-mediated protein degradation.

Beyond its chemical tractability, FBXO31 possesses biological properties that may offer opportunities distinct from those of established E3 ligases such as CRBN and VHL. FBXO31 has been reported to be stress-inducible^23^ and exhibits relatively high expression in the brain^24^. These characteristics raise the possibility of developing FBXO31-recruiting degraders for applications in specific tissues or cellular contexts. Another potential avenue for future investigation is the development of recruiters targeting disease-associated FBXO31 mutants. A recurrent de novo D334N mutation has been identified in patients with spastic cerebral palsy^25,26^. This mutation alters a key residue in the substrate-binding pocket of FBXO31. Our results demonstrate that the D334N mutation abolishes JZ26-induced degradation. Although our current recruiters are ineffective against this mutant, developing ligands that selectively recognize FBXO31 D334N could provide new opportunities to exploit disease-associated E3 ligase variants for TPD.

## Acknowledgements

We gratefully acknowledge support from the International Institute for Nanotechnology Seed Funding Program (X.Z.) and the Sherman Fairchild Bridge Fund (X.Z.)

