## Supplementary Information for "Discovery and Optimization of Indoline-2-Carboxamide-Based FBXO31 Recruiters for Targeted Protein Degradation"

3).

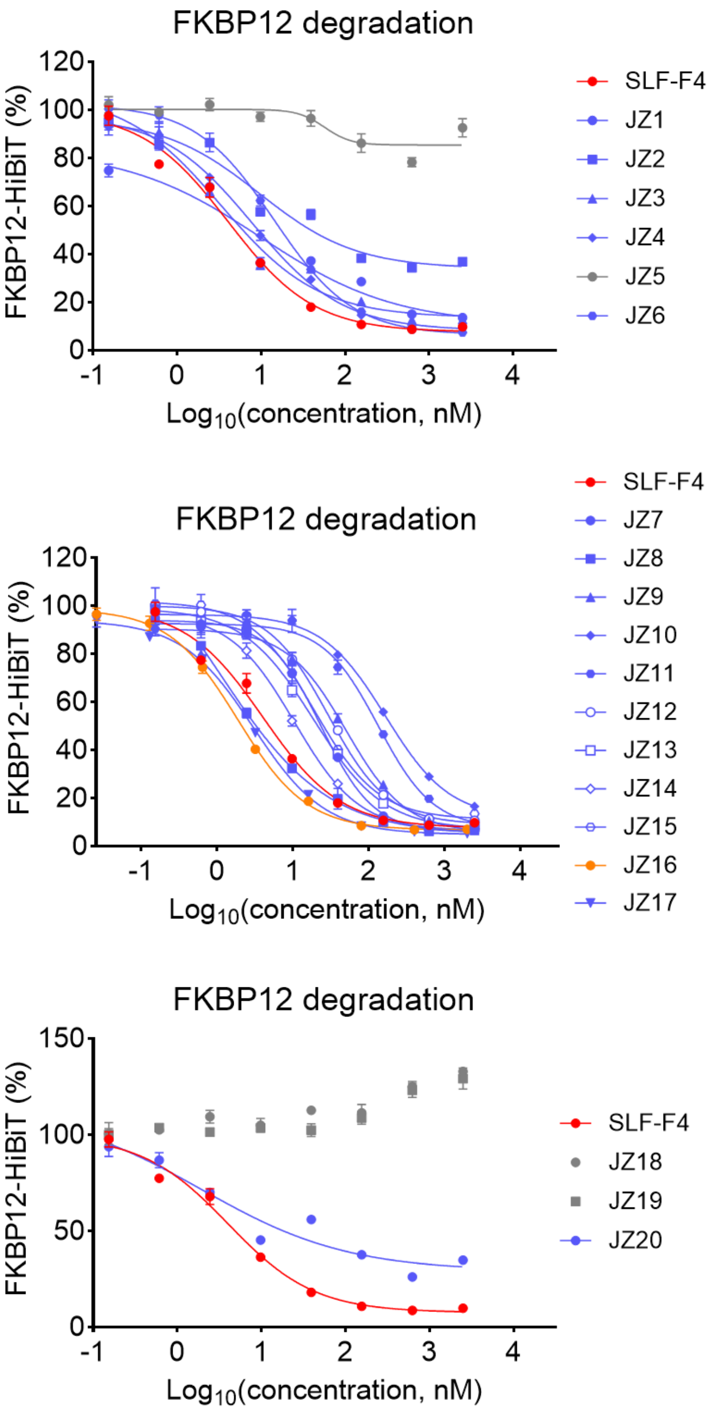


**Figure S1.** Concentration-dependent degradation of FKBP12-HiBiT in HEK293T cells treated with SLF-F4 and its analogs for 24 h. Data are presented as mean ± SEM (n = 3).

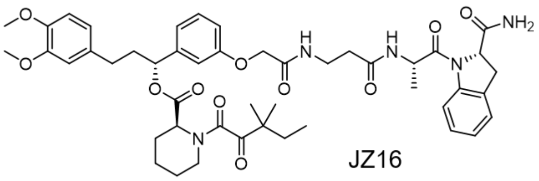

JZ16

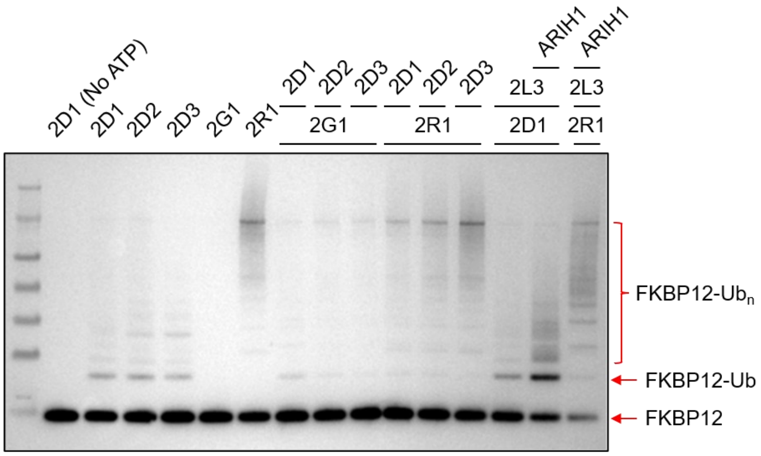


**Figure S2.** Western blot analysis of FKBP12 ubiquitination in a reconstituted in vitro ubiquitination assay using JZ16 and the indicated E2 ubiquitin-conjugating enzymes, including UBE2D1 (2D1), UBE2D2 (2D2), UBE2D3 (2D3), UBE2G1 (2G1), UBE2R1 (2R1), and UBE2L3 (2L3), either individually or in combination with ARIH1, an RBR-type E3 ubiquitin ligase. Reactions were performed with ATP unless otherwise indicated. Unmodified FKBP12, monoubiquitinated FKBP12 (FKBP12-Ub), and higher-molecular-weight ubiquitinated FKBP12 species (FKBP12-Ub$_n$) are indicated.

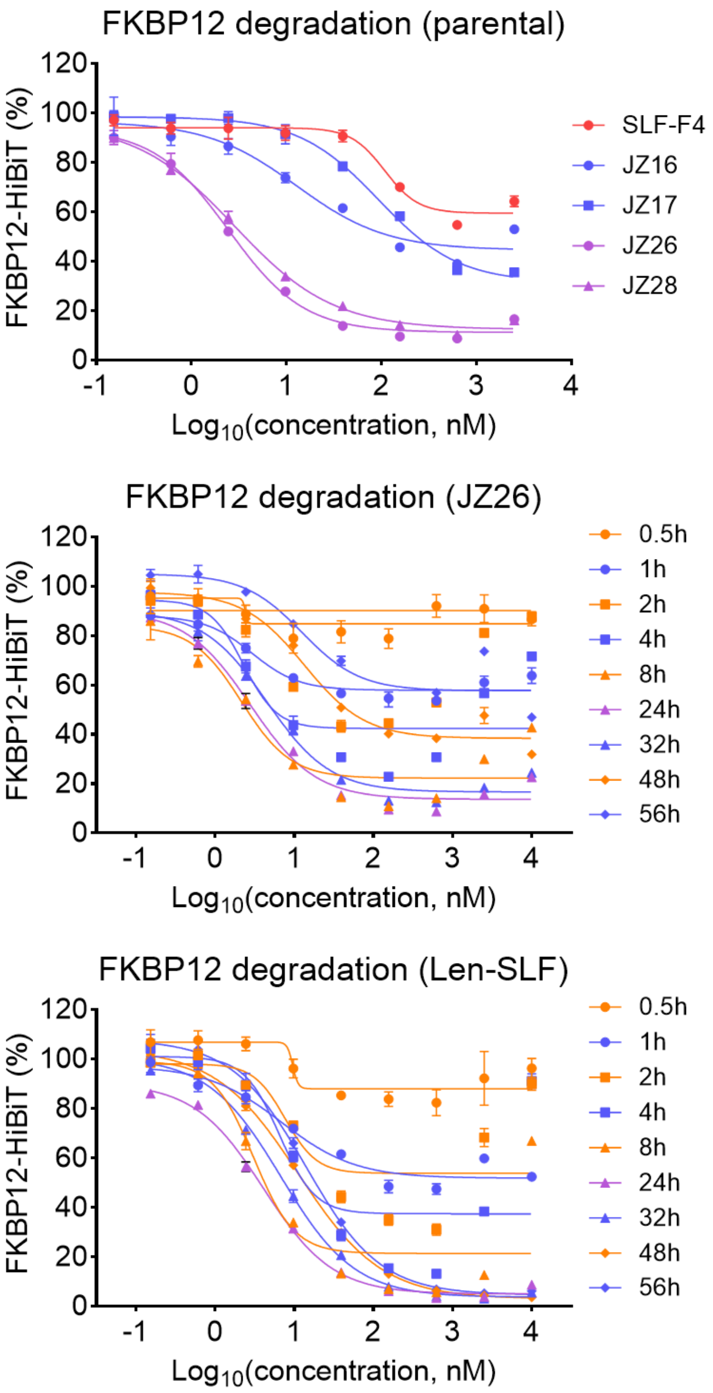


**Figure S3.** Concentration-dependent degradation of FKBP12-HiBiT in HEK293T cells treated with SLF-F4 and its analogs for 24 h, and time-course degradation study comparing JZ26 and Len-SLF. Data are presented as mean ± SEM (n = 3).

**Co-folding scores**

Focus on Protein

| Property | Value |
|---|---|
| Predicted TM-score (pTM) | 0.811 |
| ipTM | 0.820 |
| Confidence score | 0.817 |
| Average pLDDT | 0.817 |
| Affinity probability | 0.925 |
| Predicted $pIC_{50}$ | 7.849 |
| Predicted $IC_{50}$ | 14.2 nM |
| PoseBusters valid? | Yes |

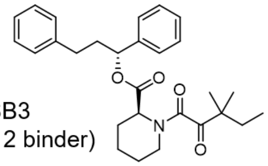

SB3
(FKBP12 binder)

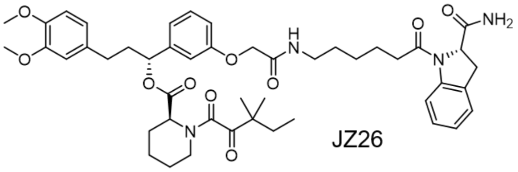

JZ26

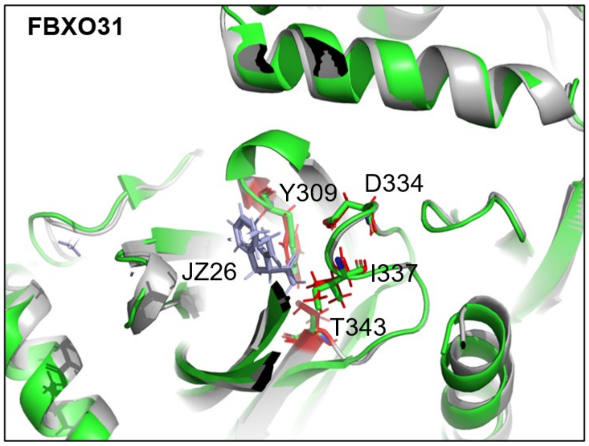


Gray: modeled FBXO31-JZ26 with key residues (red)
Green: FBXO31 with key residues (PDB: 5VZT)

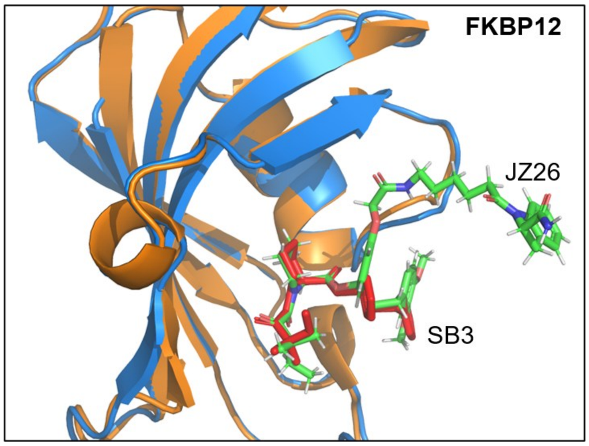


Orange: modeled FKBP12 bound to SLF-F4
Blue: FKBP12 bound to SB3 (red) (PDB: 1FKG)

**Figure S4.** Co-folding scores and alignment of the modeled FKBP12-JZ26-FBXO31 ternary complex with reported crystal structures of FKBP12-ligand complex (PDB: 1FKG) and FBXO31 (PDB: 5VZT).

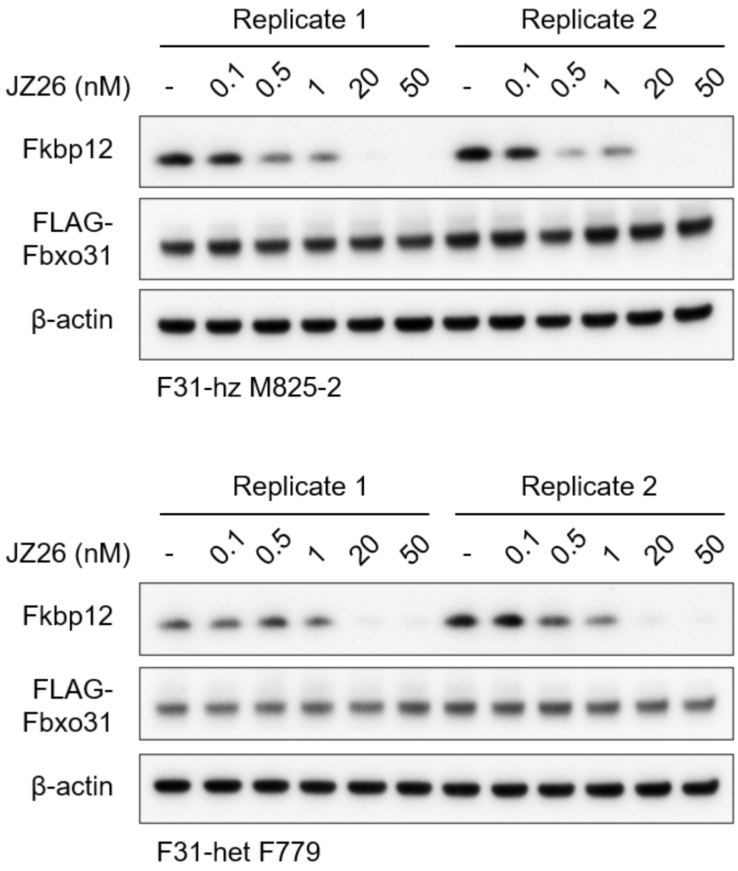


**Figure S5.** Western blot analysis of concentration-dependent endogenous mouse Fkbp12 degradation induced by JZ26 in primary mouse fibroblasts from two transgenic mouse lines overexpressing FLAG-tagged Fbxo31 (F31-hz M825-2 and F31-het F779) following 24 h of treatment. Two biological replicates are shown for each cell line.

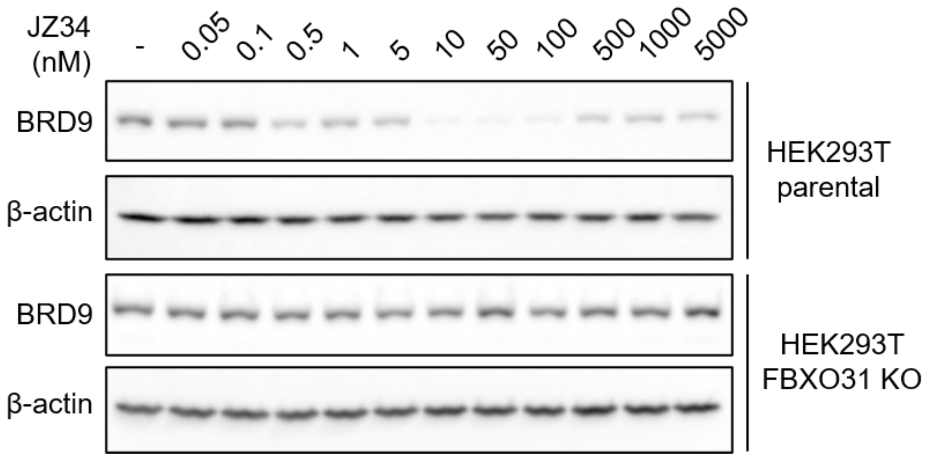


**Figure S6.** Western blot analysis of concentration-dependent BRD9 degradation induced by BRD9-C11 in parental and FBXO31-knockout HEK293T cells following 24 h of treatment.

### Supplementary Materials and Methods

#### Reagents

The anti-BRD9 (cat. #58906, dilution 1:1000), anti-SMCA2 (cat. #11966, dilution 1:1000), horseradish peroxidase (HRP)-linked rabbit IgG (cat. #7074, dilution 1:5000), and anti-β-Actin HRP (cat. #5125, dilution 1:5000) antibodies were purchased from Cell Signaling Technology. The anti-FLAG HRP antibody (clone M2, cat. #A8592, dilution 1:5000) was purchased from Sigma-Aldrich. Nano-Glo HiBiT Lytic Detection System (cat. #N3040) and sequencing grade trypsin/Lys-C mix (cat. #V5071) were purchased from Promega. Bortezomib (cat. #HY-10227), BI-7273 (cat. # HY-100351), and CFT8634 (cat. #HY-145925B) were purchased from MedChemExpress. MLN4924 (cat. #15217) and SLF (cat. #10007974) were purchased from Cayman Chemical. Pierce high pH reversed-phase peptide fractionation kit (cat. #84868), SuperSignal West Pico PLUS Chemiluminescent Substrate (cat. #34580), and Tandem Mass Tag (TMT) isobaric label reagent (cat. #90110) were purchased from Thermo Scientific. Recombinant FBXO31/SKP1 (cat. #TE3-091), neddylated CUL1/RBX1 (cat. #TE3-029), ARIH1 (cat. #TE3-136), and E2 enzymes were supplied by Trilogie Bioscience.

#### Cell Lines

HEK 239T cells were obtained from ATCC. Dulbecco's Modified Eagle Medium (DMEM, Corning, cat. #15013CV) with 10% (v/v) fetal bovine serum (FBS, Omega Scientific, cat. #FB-01) and L-glutamine (2 mM, Gibco, cat. #25030081) were used for HEK 293T cells. The HA-FBXO31 overexpressing, FBXO31 mutant expressing, FBXO31 knockout, and HiBiT-FKBP12 expressing cell lines were previously generated and characterized as described in our earlier study[1]. All cell lines were confirmed to be free of mycoplasma contamination using the Universal Mycoplasma Detection Kit (ATCC, Cat. No. 30-1012K).

#### HiBiT Assay

HiBiT-expressing HEK293T cells were seeded into 96-well white plates at 5,000 cells/well. After cell were seeded, the compounds were added in triplicates. Following compound incubation, 50 µL of Nano-Glo HiBiT Lytic Detection System reagent (Promega, cat. #N3040) was added to each well. The plates were place on shaker for 10 minutes at room temperature. Luminescence signals were then quantified using a CLARIOstar Plus microplate reader. For each experimental condition, HiBiT luminescence signals were normalized to those of the corresponding DMSO-treated control wells.

**Cell Lysis and Western Blot**

Cells were washed with ice-cold PBS then lysed RIPA buffer (Thermo Scientific, Cat. #AAJ63306AK) with Universal Nuclease (Thermo Scientific, Cat. #PI88702). DC assay (Bio-Rad, cat. #5000112) was used to normalize protein concentrations. Lysates were mixed with Laemmli sample buffer (Bio-Rad, Cat. #1610747) and 2-Mercaptoethanol (Bio-Rad, Cat. #1610710), then denatured at 95 °C for 10 minutes. The protein samples were separated with 4-20% Novex Tris-Glycine gels (Thermo Scientific, cat. #XP04205BOX) or 8% Novex Tris-Glycine gels (Thermo Scientific, cat. #XP00085BOX). Following electrophoretic separation, proteins were transferred onto 0.2 µm PVDF membranes (Bio-Rad, Cat. #1620177). The membranes were blocked with 5% nonfat milk prepared in TBST (20 mM Tris-HCl, pH 7.6, 150 mM NaCl, and 0.1% Tween-20) for 1 h at room temperature. Primary antibodies were diluted in blocking buffer and incubated at 4 °C overnight. After three TBST washes, membranes were incubated with secondary antibody for 1 hour at room temperature. Antibodies with HRP conjugated were diluted in blocking buffer and incubated at room temperature for 1 hour. Following three washes with TBST, chemiluminescent signals were developed using Pico detection reagents and captured with a ChemiDoc MP imaging system (Bio-Rad).

**Affinity Purification Mass Spectrometry (AP-MS)**

Cells were lysed in NP-40 lysis buffer (Thermo Scientific, Cat. #J60766.AP) with cOmplete protease inhibitor cocktail and disrupted by sonication. Cell lysates were centrifuged at 16,000 × g for 10 min at 4 °C to remove insoluble debris. Protein concentration was determined by DC assay. FLAG affinity agarose (Sigma-Aldrich, Cat. #A2220-1ML) (30 μL slurry per sample) was added to the lysates and incubated with rotation for 2 hours at 4 °C. The agarose was washed four times with immunoprecipitation wash buffer, followed by two additional washes with PBS. The proteins were then eluted by heating the agarose at 65 °C for 10 minutes in 8 M urea in PBS. Eluates were reduced with 12.5 mM DTT at 65 °C for 15 minutes and subsequently alkylated with 25 mM iodoacetamide at 37 °C for 30 minutes. Samples were diluted with PBS to lower the urea concentration to 2 M, followed by digestion with 2 μg of trypsin/LysC at 37 °C for 12 hours. TMT reagents (5 μL, in dry acetonitrile) were then added and reacted at room temperature for 1 hour. Reactions were quenched by addition of 6 μL of 5% hydroxylamine and 2.5 μL of formic acid. Labeled peptides were combined, desalted using a Sep-Pak C18 cartridge (Waters, cat. #WAT054955), dried by vacuum centrifugation, and subjected to LC-MS analysis using an Orbitrap Eclipse Tribrid mass spectrometer coupled to a Vanquish Neo UHPLC system.

Peptides were separated on an EASY-Spray C18 analytical column (75 μm × 150 mm, 2 μm particle size) at a flow rate of 0.25 μL/min using a multistep gradient. The mobile phases consisted of 0.1% formic acid in water (solvent A) and 80% acetonitrile containing 0.1% formic acid (solvent B). The gradient was maintained at 5% B for the first 15 min, followed by a linear increase to 45% B from 15 to 155 min and a subsequent increase to 100% B from 155 to 180 min. The nanoelectrospray voltage was set to 1.5 kV. Mass spectrometric analysis was performed using an MS3-based acquisition method with synchronous precursor selection (SPS). Full MS scans were collected in the Orbitrap at a resolution of 60,000 over an m/z range of 375-1600, with an RF lens setting of 60%, a standard automatic gain control (AGC) target, and an automatic maximum injection time. For MS2 acquisition, precursor ions were isolated using a quadrupole isolation window of 0.7 m/z and subjected to higher-energy collisional dissociation (HCD) at a normalized collision energy of 27%. MS2 spectra were acquired in the ion trap with a standard AGC target and a maximum injection time of 35 ms. Up to ten MS2 fragment ions were selected for SPS-MS3 analysis, followed by HCD fragmentation at a normalized collision energy of 55%. MS3 spectra were recorded in the Orbitrap at a resolution of 60,000, with an AGC target of 250% and a maximum injection time of 200 ms. Raw mass spectrometry data were acquired using Xcalibur software (version 4.5.445.18, Thermo Scientific).

Raw LC-MS/MS data were processed using Proteome Discoverer (version 2.5, Thermo Scientific). Peptide sequences were identified using the Sequest HT search engine against the UniProt human reference proteome database (UP000005640_9606_Human.fasta). Database searches were conducted with trypsin as the specified protease, allowing precursor and fragment mass tolerances of 10 ppm and 0.6 Da, respectively. Carbamidomethylation of cysteine residues and TMT modifications of peptide N-termini and lysine residues were assigned as static modifications, whereas methionine oxidation and protein N-terminal acetylation were considered dynamic modifications. Peptide-spectrum matches (PSMs) were assessed using Percolator, and peptide- and protein-level identifications were filtered at a false discovery rate (FDR) of 1%. TMT reporter ion intensities were quantified from SPS-MS3 spectra using a mass tolerance of 20 ppm. Protein abundances were determined using unique and razor peptides.

**Compound Treatment of Primary Mouse Fibroblasts**

Mouse primary fibroblasts were isolated from the ear skin of transgenic mice harboring 3xFLAG-FBXO31 overexpression cassette at the ROSA26 locus, following a previously published protocol[2]. Cells were maintained in RPMI 1640 medium supplemented with 10% FBS, 50 μM β-mercaptoethanol, and 1% penicillin-streptomycin. For compound treatment, cells were seeded in 12-well plates at a density of $2 \times 10^5$ cells/mL and treated with the indicated compounds diluted in fresh medium for 24 h. For MLN4924 treatment, cells were co-treated with the indicated compounds and 2 μM MLN4924 for 24 h. Following treatment, cells were washed with PBS and lysed in RIPA buffer. Cell lysates were then subjected to SDS-PAGE using 4-12% gradient gels, followed by Western blot analysis. The following primary antibodies were used: rabbit polyclonal anti-FKBP1A (Proteintech, 10273-1-AP; 1:2000), mouse monoclonal anti-FLAG M2 (Sigma-Aldrich,

F3165; 1:2000), and mouse monoclonal anti-β-actin (AC-15; Sigma-Aldrich, A5441; 1:2000). Amersham ECL HRP-conjugated secondary antibodies (NA931 or NA934) were used for detection with SuperSignal ECL detection reagents.

**In Vitro Ubiquitylation Assays**

Ubiquitylation assays were performed in a final volume of 25 µL. His-FBXO31/SKP1, neddylated CUL1/RBX1, ubiquitin, UBA1, biotinylated Avi-tagged FKBP12, JZ16, and reaction buffer were combined and pre-incubated for 30 min at room temperature. The final reaction buffer contained 50 mM HEPES (pH 7.5), 50 mM NaCl, and 1 mM TCEP. Reactions were initiated by adding Mg-ATP (pH 7.0) and the indicated E2 enzymes, with or without ARIH1. Final reaction concentrations were 250 nM His-FBXO31/SKP1, 250 nM neddylated CUL1/RBX1, 20 µM ubiquitin, 250 nM UBA1, 2 µM biotinylated FKBP12, 2 µM JZ16, 10 mM ATP, 10 mM $Mg^{2+}$ and 2 µM total E2. Individual E2 reactions contained UBE2D1, UBE2D2, UBE2D3, UBE2G1, or UBE2R1. E2 combinations included UBE2D1, UBE2D2, or UBE2D3 with either UBE2G1 or UBE2R1; UBE2D1 with UBE2L3, with or without ARIH1; and UBE2R1 with UBE2L3 and ARIH1. Where included, ARIH1 was present at 250 nM. A reaction containing UBE2D1 without ATP served as a control. Reactions were incubated for 60 min at 37 °C and quenched directly into SDS-PAGE loading buffer. Samples were separated by SDS-PAGE and transferred to a nitrocellulose membrane. Biotinylated FKBP12 was detected using streptavidin-HRP diluted 1:20000 in 5% BSA in PBST, followed by imaging with West Pico ECL.

**Statistical Analysis**

Quantitative data are displayed as scatter plots with the mean ± standard error of the mean (SEM). Statistical significance in proteomics experiments was assessed using two-sided Student's t-tests, and *P* values were adjusted for multiple comparisons using the Benjamini-Hochberg procedure. Statistical significance was defined as $P < 0.05$.

### Synthesis of JZ1-6

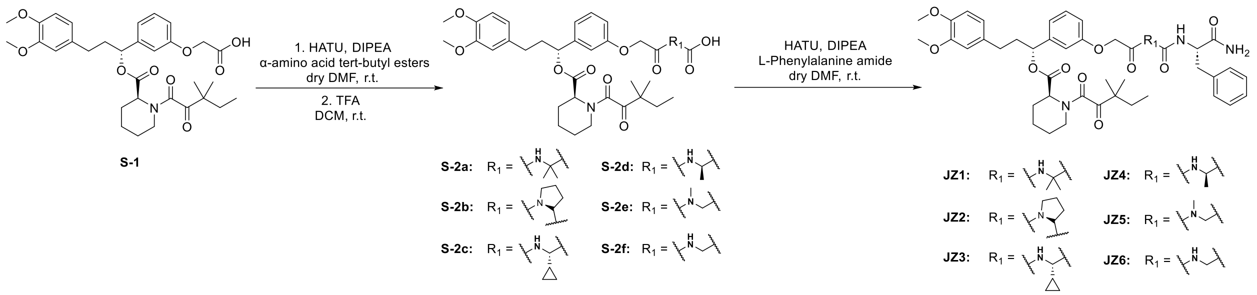

**JZ1:** *(R)-1-(3-(2-((3-((1-(((S)-1-amino-1-oxo-3-phenylpropan-2-yl)amino)-2-methyl-1-oxopropan-2-yl)amino)-3-oxopropyl)amino)-2-oxoethoxy)phenyl)-3-(3,4-dimethoxyphenyl)propyl (S)-1-(3,3-dimethyl-2-oxopentanoyl)piperidine-2-carboxylate*

Step 1: S-1 was synthesized according to the previously reported method[3]. S-1 (45 mg, 0.07 mmol, 1.0 eq), HATU (34 mg, 0.09 mmol, 1.3 eq), and tert-Butyl 2-amino-2-methylpropanoate hydrochloride (27 mg, 0.14 mmol, 2 eq) were dissolved in dry DMF (2.0 mL) at room temperature. DIPEA (59 µL, 0.34 mmol, 5.0 eq) was added to the reaction and stirred for 2 hours. The reaction was then concentrated and extracted with DCM (30 mL) and brine (30 mL). The organic layer was dried with anhydrous sodium sulfate. The concentrated resulting residue was dissolved in DCM (5.0 mL), and TFA (1.0 mL) was added and stirred for 2 hours. The reaction mixture was then concentrated and purified by flash chromatography to provide S-2a as a yellow oil (36.7 mg, 0.05 mmol, 72.5% over 2 steps).

$^1$H NMR (500 MHz, $CDCl_3$) δ 7.56 (s, 1H), 7.29 – 7.22 (m, 1H), 7.00 – 6.89 (m, 3H), 6.85 – 6.72 (m, 2H), 6.70 – 6.63 (m, 2H), 5.78 – 5.72 (m, 1H), 5.29 (t, *J* = 3.4 Hz, 1H), 4.49 (s, 2H), 3.90 – 3.79 (m, 6H), 3.62 (s, 2H), 3.47 (s, 1H), 3.38 – 3.32 (m, 1H), 3.25 – 3.12 (m, 1H), 2.66 – 2.49 (m, 2H), 2.44 (s, 2H), 2.41 – 2.34 (m, 1H), 2.30 – 2.17 (m, 1H), 2.09 – 2.00 (m, 1H), 1.83 – 1.58 (m, 5H), 1.53 – 1.29 (m, 8H), 1.22 – 1.10 (m, 6H), 0.83 (dt, *J* = 28.2, 7.4 Hz, 3H).

Step 2: S-2a (36.7 mg, 0.05 mmol, 1eq), HATU (24.5 mg, 0.06 mmol, 1.3 eq), and L-Phenylalanine amide (16.3 mg, 0.10 mmol, 2 eq) were dissolved in dry DMF (2.0 mL) at room temperature. DIPEA (42.8 µL, 0.43 mmol, 5.0 eq) was added to the reaction and stirred for 2 hours. The reaction was then concentrated and extracted with DCM (30 mL) and brine (30 mL). The organic layer was dried with anhydrous sodium sulfate. The reaction mixture was then concentrated and purified by flash chromatography to provide JZ1 as a colorless oil (30.4 mg, 0.03 mmol, 69.2% yield).

$^1$H NMR (500 MHz, $CDCl_3$) δ 7.32-7.22 (m, 4H), 7.19-7.15 (m, 3H), 7.02-6.94 (m, 2H), 6.91-6.89 (m, 1H), 6.84-6.81 (m, 2H), 6.78-6.76 (m, 1H), 6.72-6.66 (m, 3H), 5.77-5.72 (m, 1H), 5.57-5.51 (m, 1H), 5.27-5.25 (m, 1H), 4.67-4.62 (m, 1H), 4.50-4.43 (m, 2H), 3.84-3.82 (m, 6H), 3.59-3.48 (m, 2H), 3.37-3.17 (m, 3H), 3.13-3.06 (m, 1H), 2.63-2.50 (m, 2H), 2.41-2.33 (m, 3H), 2.27-2.19 (m, 1H), 2.08-2.01 (m, 1H), 1.82-1.61 (m, 5H), 1.51-1.31 (m, 5H), 1.20-1.13 (m, 9H), 0.88-0.79 (m, 3H). $^{13}$C NMR (126 MHz, $CDCl_3$) δ 208.13, 173.91, 172.26, 169.70, 169.07, 167.53, 157.56, 149.04, 147.53, 141.95, 137.47, 133.41, 130.21, 129.28, 128.61, 126.86, 120.28, 114.48, 113.12, 111.86, 111.48, 67.43, 57.14, 56.05, 55.98, 54.03, 51.51, 46.83, 44.29, 38.31, 36.76, 36.70, 35.46, 32.57, 31.44, 26.63, 26.05, 25.03, 24.16, 23.49, 23.38, 21.28, 8.87. HRMS (ESI$^+$) m/z calcd for $C_{48}H_{64}N_5O_{11}^+$ [M+H]$^+$: 886.4602, found 886.4554.

The synthetic route of JZ2, JZ3, JZ4, JZ5, JZ6 were similar to JZ1.

**JZ2:** *(R)-1-(3-(2-((3-((R)-2-(((S)-1-amino-1-oxo-3-phenylpropan-2-yl)carbamoyl)pyrrolidin-1-yl)-3-oxopropyl)amino)-2-oxoethoxy)phenyl)-3-(3,4-dimethoxyphenyl)propyl (S)-1-(3,3-dimethyl-2-oxopentanoyl)piperidine-2-carboxylate*

$^1$H NMR (500 MHz, $CDCl_3$) δ 7.37-7.34 (m, 1H), 7.30-7.23 (m, 4H), 7.21-7.16 (m, 3H), 7.08-7.06 (m, 1H), 6.99-6.90 (m, 2H), 6.84-6.82 (m, 1H), 6.78-6.76 (m, 1H), 6.70-6.60 (m, 3H), 5.78-5.74 (m, 1H), 5.53-5.42 (m, 1H), 5.27 (s, 1H), 4.71-4.66 (m, 1H), 4.52-4.42 (m, 2H), 4.38-4.35 (m, 1H), 3.85-3.83 (m, 6H), 3.62-3.59 (m, 2H), 3.42-3.31 (m, 2H), 3.27-3.23 (m, 1H), 3.20-3.14 (m, 1H), 3.09-3.04 (m, 1H), 2.63-2.35 (m, 5H), 2.28-

2.19 (m, 1H), 2.07-1.62 (m, 10H), 1.53-1.30 (m, 2H), 1.24-1.12 (m, 6H), 0.88-0.78 (m, 3H). $^{13}$C NMR (126 MHz, $CDCl_3$) δ 208.00, 173.64, 172.06, 171.27, 169.80, 168.67, 167.44, 157.48, 149.01, 147.49, 141.97, 137.42, 133.46, 130.12, 129.30, 128.51, 126.87, 120.27, 120.19, 114.05, 113.79, 111.85, 111.44, 67.36, 60.93, 56.03, 55.96, 53.56, 51.43, 47.65, 46.81, 44.27, 38.20, 36.91, 34.89, 34.82, 32.58, 31.37, 28.64, 26.53, 25.02, 24.72, 23.53, 23.52, 23.31, 21.28, 8.87. HRMS (ESI$^+$) m/z calcd for $C_{49}H_{63}N_5NaO_{11}^+$ [M+Na]$^+$: 920.4422, found 920.4380.

**JZ3:** *(R)-1-(3-(2-((3-(((S)-2-(((S)-1-amino-1-oxo-3-phenylpropan-2-yl)amino)-1-cyclopropyl-2-oxoethyl)amino)-3-oxopropyl)amino)-2-oxoethoxy)phenyl)-3-(3,4-dimethoxyphenyl)propyl (S)-1-(3,3-dimethyl-2-oxopentanoyl)piperidine-2-carboxylate*

$^1$H NMR (500 MHz, $CDCl_3$) δ 7.44 (t, *J* = 6.1 Hz, 1H), 7.38 – 7.33 (m, 1H), 7.32 – 7.28 (m, 2H), 7.28 – 7.22 (m, 3H), 7.17 (d, *J* = 8.0 Hz, 1H), 7.10 – 7.02 (m, 2H), 7.02 – 6.97 (m, 1H), 6.90 (dd, *J* = 8.3, 2.6 Hz, 1H), 6.88 – 6.82 (m, 1H), 6.78 – 6.67 (m, 3H), 5.86 – 5.78 (m, 1H), 5.72 (s, 1H), 5.37 (s, 2H), 4.77 (q, *J* = 7.2 Hz, 1H), 4.57 – 4.46 (m, 2H), 3.95 – 3.87 (m, 6H), 3.79 – 3.61 (m, 3H), 3.50 – 3.39 (m, 1H), 3.31 – 3.19 (m, 2H), 3.17 – 3.07 (m, 1H), 2.74 – 2.49 (m, 4H), 2.36 – 2.26 (m, 1H), 2.15 – 2.06 (m, 1H), 1.91 – 1.67 (m, 5H), 1.60 – 1.38 (m, 2H), 1.31 – 1.20 (m, 6H), 1.11 – 0.98 (m, 1H), 0.97 – 0.86 (m, 3H), 0.60 – 0.49 (m, 2H), 0.45 – 0.34 (m, 2H).$^{13}$C NMR (126 MHz, $CDCl_3$) δ 208.04, 173.39, 171.93, 171.02, 169.80, 168.76, 167.47, 157.54, 149.01, 147.49, 141.91, 136.86, 133.43, 130.12, 129.43 (2C), 128.61 (2C), 126.97, 120.27, 114.30, 113.50, 111.86, 111.46, 67.41, 58.25, 56.03, 55.95, 54.02, 53.55, 51.46, 46.82, 44.28, 38.23, 37.76, 35.84, 35.52, 32.56, 31.38, 28.05, 26.55, 25.02, 23.53, 23.29, 21.28, 13.59, 8.86, 3.62 (2C). HRMS (ESI$^+$) m/z calcd for $C_{49}H_{63}N_5NaO_{11}^+$ [M+Na]$^+$: 920.4422, found 920.4363.

**JZ4:** *(R)-1-(3-(2-((3-(((R)-1-(((S)-1-amino-1-oxo-3-phenylpropan-2-yl)amino)-1-oxopropan-2-yl)amino)-3-oxopropyl)amino)-2-oxoethoxy)phenyl)-3-(3,4-dimethoxyphenyl)propyl (S)-1-(3,3-dimethyl-2-oxopentanoyl)piperidine-2-carboxylate*

$^{1}H$ NMR (500 MHz, MeOD) δ 7.33 – 7.27 (m, 1H), 7.26 – 7.19 (m, 4H), 7.19 – 7.14 (m, 1H), 7.01 – 6.95 (m, 2H), 6.94 – 6.89 (m, 1H), 6.84 (d, *J* = 8.2 Hz, 1H), 6.79 – 6.74 (m, 1H), 6.72 – 6.67 (m, 1H), 5.77 – 5.71 (m, 1H), 5.47 – 5.43 (m, 1H), 5.25 – 5.20 (m, 1H), 4.64 – 4.55 (m, 1H), 4.52 – 4.47 (m, 2H), 4.20 – 4.12 (m, 1H), 3.83 – 3.77 (m, 6H), 3.57 – 3.47 (m, 2H), 3.24 – 3.14 (m, 1H), 3.03 – 2.93 (m, 1H), 2.88 – 2.77 (m, 1H), 2.68 – 2.50 (m, 2H), 2.46 – 2.40 (m, 2H), 2.29 – 2.20 (m, 1H), 2.10 – 2.02 (m, 1H), 1.81 – 1.56 (m, 5H), 1.51 – 1.29 (m, 3H), 1.24 – 1.02 (m, 9H), 0.91 – 0.76 (m, 3H). $^{13}C$ NMR (126 MHz, MeOD) δ 208.92, 170.85, 168.92, 159.07, 150.21, 148.63, 143.11, 138.60, 138.39, 134.99, 130.23, 130.10 (2C), 129.29 (2C), 127.60, 121.56, 120.90, 115.47, 114.33, 113.37, 113.03, 79.32, 79.06, 78.80, 68.14, 56.51, 56.41, 52.67, 50.93, 50.69, 47.60, 45.56, 39.06, 38.26, 36.45, 36.05, 33.45, 32.16, 27.25, 25.79, 23.82, 23.62, 22.03, 17.46, 17.35, 9.09. HRMS (ESI$^{+}$) m/z calcd for $C_{47}H_{61}N_{5}NaO_{11}^{+}$ [M+Na]$^{+}$: 894.4265, found 894.4210.

**JZ5:** *(R)-1-(3-(2-((3-((2-(((S)-1-amino-1-oxo-3-phenylpropan-2-yl)amino)-2-oxoethyl)(methyl)amino)-3-oxopropyl)amino)-2-oxoethoxy)phenyl)-3-(3,4-dimethoxyphenyl)propyl (S)-1-(3,3-dimethyl-2-oxopentanoyl)piperidine-2-carboxylate*

$^{1}H$ NMR (500 MHz, MeOD) δ 7.24-7.05 (m, 6H), 6.93-6.85 (m, 2H), 6.84 -6.81 (m, 1H), 6.76-6.73 (m, 1H), 6.70-6.65 (m, 1H), 6.62-6.59 (m, 1H), 5.68-5.62 (m, 1H), 4.66 – 4.51 (m, 1H), 4.42-4.37 (m, 2H), 3.96-3.76 (m, 2H), 3.70 (s, 3H), 3.69 (s, 3H), 3.45-3.36 (m, 2H), 3.31-3.27 (m, 1H), 3.15 – 3.05 (m, 2H), 2.84-2.72 (m, 3H), 2.65-2.63 (m, 1H), 2.58-2.42 (m, 3H), 2.37-2.09 (m, 3H), 2.02-1.90 (m, 1H), 1.71-1.47 (m, 5H), 1.43-1.17 (m, 3H), 1.13-0.99 (m, 6H), 0.81-0.67 (m, 3H). $^{13}C$ NMR (126 MHz, MeOD) δ 209.00, 175.96, 174.32, 170.95, 170.85, 169.03, 159.18, 150.36, 148.77, 143.26, 138.57, 135.14, 130.97, 130.27 (2C), 129.45 (2C), 127.73, 121.66, 120.99, 113.51, 113.18, 79.47, 79.21, 78.94, 77.95, 68.25, 58.23, 56.55, 56.45, 55.60, 52.77, 47.67, 45.67, 39.18, 38.69, 37.09, 36.10, 33.75, 33.54, 32.22, 27.31, 25.87, 23.84, 23.65, 22.11, 9.11. HRMS (ESI$^{+}$) m/z calcd for $C_{47}H_{61}N_{5}NaO_{11}^{+}$ [M+Na]$^{+}$: 894.4265, found 894.4211.

**JZ6:** *(R)-1-(3-(2-((3-((2-(((S)-1-amino-1-oxo-3-phenylpropan-2-yl)amino)-2-oxoethyl)amino)-3-oxopropyl)amino)-2-oxoethoxy)phenyl)-3-(3,4-dimethoxyphenyl)propyl (S)-1-(3,3-dimethyl-2-oxopentanoyl)piperidine-2-carboxylate*

$^{1}$H NMR (500 MHz, MeOD) δ 7.31-7.28 (m, 1H), 7.26-7.23 (m, 1H), 7.23-7.21 (m, 2H), 7.21-7.19 (m, 1H), 7.18-7.15 (m, 1H), 7.01-6.97 (m, 2H), 6.93-6.91 (m, 1H), 6.85-6.83 (m, 1H), 6.80-6.77 (m, 1H), 6.71-6.69 (m, 1H), 5.75-5.72 (m, 1H), 5.23-5.21 (m, 1H), 4.63-4.60 (m, 1H), 4.50 (s, 2H), 3.83-3.69 (m, 8H), 3.56-3.52 (m, 2H), 3.35 (s, 1H), 3.21-3.16 (m, 2H), 2.95-2.89 (m, 1H), 2.64-2.56 (m, 2H), 2.48-2.45 (m, 2H), 2.35-2.29 (m, 1H), 2.28-2.21 (m, 2H), 2.08-2.05 (m, 1H), 1.75-1.62 (m, 5H), 1.49-1.45 (m, 1H), 1.35-1.29 (m, 1H), 1.21-1.10 (m, 6H), 0.89-0.77 (m, 3H). $^{13}$C NMR (126 MHz, MeOD) δ 209.04, 176.02, 174.58, 171.40, 171.01, 169.09, 159.27, 150.41, 148.83, 143.31, 138.54, 135.20, 130.99, 130.28 (2C), 129.46 (2C), 127.75, 121.70, 120.97, 115.61, 114.36, 113.58, 113.25, 78.01, 68.25, 58.27, 56.55, 56.45, 55.72, 52.82, 47.69, 45.70, 43.65, 39.19, 38.70, 36.63, 36.43, 33.57, 32.24, 27.33, 25.88, 23.84, 23.64, 22.13, 9.12. HRMS (ESI$^{+}$) m/z calcd for $C_{46}H_{59}N_5NaO_{11}^{+}$ [M+Na]$^{+}$: 880.4109, found 880.4053.

**Synthesis of JZ7-19**

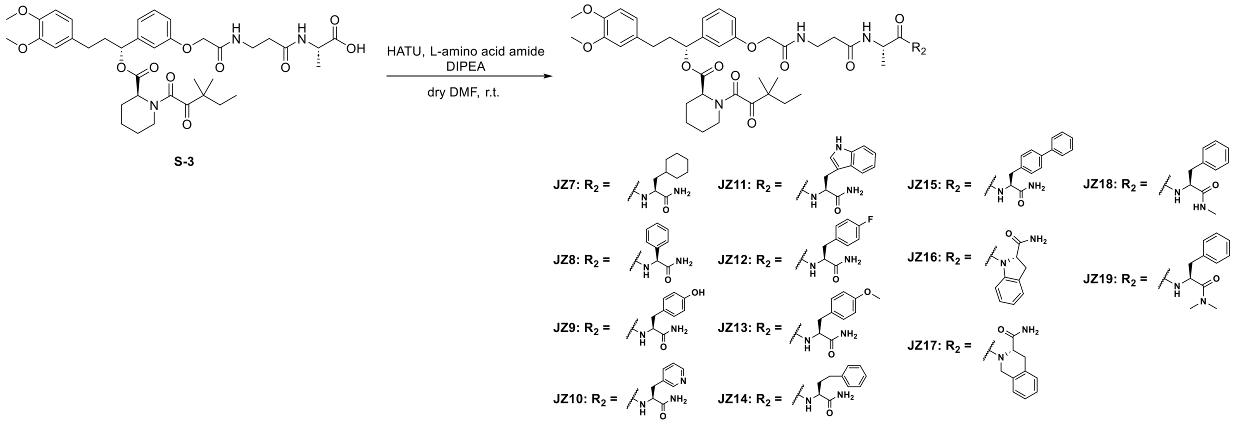


**JZ7:** *(R)-1-(3-(2-((3-(((S)-1-(((S)-1-amino-3-cyclohexyl-1-oxopropan-2-yl)amino)-1-oxopropan-2-yl)amino)-3-oxopropyl)amino)-2-oxoethoxy)phenyl)-3-(3,4-dimethoxyphenyl)propyl (S)-1-(3,3-dimethyl-2-oxopentanoyl)piperidine-2-carboxylate*

S-3 was synthesized according to the previously reported method[1]. S-3 (23 mg, 0.03 mmol, 1 eq), HATU (16 mg, 0.04 mmol, 1.3 eq), and (*S*)-2-amino-3-cyclohexylpropanamide hydrochloride (13 mg, 0.06 mmol, 2 eq) were dissolved in dry DMF (2.0 mL) at room temperature. DIPEA (27 μL, 0.16 mmol, 5.0 eq) was added to the reaction and stirred for 2 hours. The reaction was then concentrated and extracted with DCM (30 mL) and brine (30 mL). The organic layer was dried with anhydrous sodium sulfate. The reaction mixture was then concentrated and purified by flash chromatography to provide JZ7 as a yellow oil (16.3 mg, 0.02 mmol, 58.6% yield).

$^1$H NMR (500 MHz, $CDCl_3$) δ 7.39 (s, 1H), 7.30-7.27 (m, 1H), 7.00-6.92 (m, 3H), 6.85-6.82 (m, 1H), 6.80-6.77 (m, 1H), 6.70-6.67 (m, 3H), 6.56 (s, 1H), 5.78-5.73 (m, 1H), 5.52 (s, 1H), 5.28-5.27 (m, 1H), 4.49-4.45 (m, 3H), 4.34-4.29 (m, 1H), 3.86-3.84 (m, 6H), 3.63 (s, 2H), 3.38-3.35 (m, 1H), 3.23-3.17 (m, 1H), 2.63-2.47 (m, 4H), 2.40-2.38 (m, 1H), 2.27-2.20 (m, 1H), 2.09-2.03 (m, 1H), 1.81-1.50 (m, 12H), 1.36-1.10 (m, 15H), 0.93-0.80 (m, 5H). $^{13}$C NMR (126 MHz, $CDCl_3$) δ 208.09, 175.07, 172.64, 172.08, 169.80, 168.93, 167.56, 157.56, 149.05, 147.54, 141.95, 133.44, 130.17, 120.31, 120.22, 114.21, 113.54, 111.88, 111.50, 67.40, 56.07, 56.00, 51.54, 50.93, 46.86, 44.32, 39.15, 38.29, 36.00, 35.42, 34.31, 33.76, 32.59, 32.46, 31.43, 29.83, 26.62, 26.48, 26.35, 26.18, 25.05, 23.57, 23.34, 21.31, 18.05, 8.89. HRMS ($ESI^+$) m/z calcd for $C_{47}H_{67}N_5NaO_{11}^+$ $[M+Na]^+$: 900.4735, found 900.4685.

The synthetic route of JZ8-JZ19 were similar to JZ7.

**JZ8:** *(R)-1-(3-(2-((3-(((S)-1-(((S)-2-amino-2-oxo-1-phenylethyl)amino)-1-oxopropan-2-yl)amino)-3-oxopropyl)amino)-2-oxoethoxy)phenyl)-3-(3,4-dimethoxyphenyl)propyl (S)-1-(3,3-dimethyl-2-oxopentanoyl)piperidine-2-carboxylate*

$^1$H NMR (500 MHz, $CDCl_3$) δ 7.77 – 7.73 (m, 1H), 7.39-7.35 (m, 2H), 7.33-7.31 (m, 1H), 7.28-7.22 (m, 4H), 7.05-6.87 (m, 3H), 6.79-6.77 (m, 2H), 6.69-6.66 (m, 2H), 6.53 (s, 1H), 5.78-5.72 (m, 1H), 5.64 (s, 1H), 5.56-5.52 (m, 1H), 5.28-5.27 (m, 1H), 4.67-4.57 (m, 1H),

4.46-4.40 (m, 2H), 3.85-3.83 (m, 6H), 3.66-3.56 (m, 2H), 3.37-3.35 (m, 1H), 3.20-3.14 (m, 1H), 2.63-2.44 (m, 4H), 2.38-2.35 (m, 1H), 2.27-2.17 (m, 1H), 2.07-1.99 (m, 1H), 1.78-1.61 (m, 5H), 1.51-1.44 (m, 1H), 1.38-1.33 (m, 4H), 1.22-1.11 (m, 6H), 0.89-0.78 (m, 3H). $^{13}$C NMR (126 MHz, $CDCl_3$) δ 208.02, 172.02, 171.90, 171.33, 169.88, 168.77, 167.54, 157.57, 149.04, 147.53, 141.89, 137.81, 133.46, 130.08, 129.08, 128.51, 127.29, 120.30, 120.21, 114.11, 113.61, 111.88, 111.49, 67.34, 56.77, 56.06, 55.99, 51.50, 49.33, 46.84, 44.30, 38.24, 35.66, 35.44, 32.56, 31.42, 26.55, 25.03, 23.56, 23.28, 21.31, 18.97, 8.87. HRMS (ESI$^+$) m/z calcd for $C_{46}H_{59}N_5NaO_{11}^+$ [M+Na]$^+$: 880.4109, found 880.4056.

**JZ9:** *(R)-1-(3-(2-((3-(((S)-1-(((S)-1-amino-3-(4-hydroxyphenyl)-1-oxopropan-2-yl)amino)-1-oxopropan-2-yl)amino)-3-oxopropyl)amino)-2-oxoethoxy)phenyl)-3-(3,4-dimethoxyphenyl)propyl (S)-1-(3,3-dimethyl-2-oxopentanoyl)piperidine-2-carboxylate*

$^1$H NMR (500 MHz, $CDCl_3$) δ 7.27 – 7.22 (m, 2H), 7.95-7.85 (m, 4H), 6.82-6.79 (m, 1H), 6.75-6.72 (m, 1H), 6.68-6.61 (m, 4H), 5.72-5.28 (m, 1H), 5.26-5.20 (m, 1H), 4.49-4.37 (m, 3H), 4.22-4.15 (m, 1H), 3.80-3.78 (m, 6H), 3.46-3.39 (m, 2H), 3.32-3.29 (m, 1H), 3.17-3.04 (m, 3H), 2.97-2.87(m, 2H), 2.59-2.44 (m, 2H), 2.34-2.31 (m, 3H), 2.22-2.14 (m, 1H), 2.04-1.96 (m, 1H), 1.74-1.56 (m, 5H), 1.45-1.27 (m, 4H), 1.22-1.06 (m, 10H), 0.84-0.73 (m, 3H). $^{13}$C NMR (126 MHz, $CDCl_3$) δ 208.11, 172.54, 172.18, 169.77, 169.24, 167.53, 157.42, 155.79, 148.87, 147.35, 141.77, 141.76, 133.43, 130.33, 130.06, 127.23, 120.26, 120.09, 115.40, 114.25, 113.38, 111.81, 111.46, 67.12, 55.93, 55.84, 53.78, 51.44, 46.75, 46.74, 44.28, 38.11, 36.62, 35.24, 35.17, 32.43, 31.27, 26.42, 24.87, 23.37, 23.10, 21.12, 17.07, 8.67. HRMS (ESI$^+$) m/z calcd for $C_{47}H_{61}N_5NaO_{12}^+$ [M+Na]$^+$: 910.4214, found 910.4150.

**JZ10:** *(R)-1-(3-(2-((3-(((S)-1-(((S)-1-amino-1-oxo-3-(pyridin-3-yl)propan-2-yl)amino)-1-oxopropan-2-yl)amino)-3-oxopropyl)amino)-2-oxoethoxy)phenyl)-3-(3,4-dimethoxyphenyl)propyl (S)-1-(3,3-dimethyl-2-oxopentanoyl)piperidine-2-carboxylate*

$^1$H NMR (500 MHz, $CDCl_3$) δ 8.41 (s, 1H), 7.77 – 7.39 (m, 2H), 7.32 – 7.21 (m, 2H), 7.14 (s, 1H), 7.00 – 6.87 (m, 3H), 6.84 – 6.73 (m, 2H), 6.69 – 6.61 (m, 2H), 5.95 – 5.45 (m,

2H), 5.34 – 5.21 (m, 1H), 4.73 (s, 1H), 4.53 – 4.14 (m, 4H), 3.88 – 3.79 (m, 6H), 3.59 (s, 2H), 3.46 (s, 2H), 3.39 – 3.31 (m, 1H), 3.25 – 3.14 (m, 1H), 3.11 – 2.78 (m, 1H), 2.69 – 2.30 (m, 5H), 2.31 – 2.15 (m, 1H), 2.12 – 1.98 (m, 1H), 1.82 – 1.57 (m, 5H), 1.54 – 1.42 (m, 1H), 1.40 – 1.30 (m, 1H), 1.29 – 1.12 (m, 9H), 0.91 – 0.78 (m, 3H). $^{13}$C NMR (126 MHz, $CDCl_3$) δ 208.06, 173.00, 172.66, 172.13, 169.82, 169.03, 167.59, 166.71, 157.57, 149.03, 147.52, 141.92, 133.42, 130.13, 120.29, 120.18, 114.13, 113.64, 111.87, 111.48, 67.41, 56.84, 56.05, 55.98, 53.55, 51.53, 46.84, 44.30, 39.06, 38.23, 36.05, 35.60, 32.57, 31.41, 29.81, 27.75, 26.58, 25.01, 23.78, 23.52, 23.33, 21.29, 17.82, 14.24, 8.87. HRMS (ESI$^+$) m/z calcd for $C_{46}H_{61}N_6O_{11}^+$ [M+H]$^+$: 873.4398, found 873.4350.

**JZ11:** *(R)-1-(3-(2-((3-(((S)-1-(((S)-1-amino-3-(1H-indol-3-yl)-1-oxopropan-2-yl)amino)-1-oxopropan-2-yl)amino)-3-oxopropyl)amino)-2-oxoethoxy)phenyl)-3-(3,4-dimethoxyphenyl)propyl (S)-1-(3,3-dimethyl-2-oxopentanoyl)piperidine-2-carboxylate*

$^1$H NMR (500 MHz, $CDCl_3$) δ 8.87 (s, 1H), 7.60 (d, *J* = 7.8 Hz, 1H), 7.33 – 7.18 (m, 4H), 7.12 (t, *J* = 7.5 Hz, 1H), 7.07 (t, *J* = 7.3 Hz, 1H), 7.01 (s, 1H), 6.98 – 6.93 (m, 2H), 6.89 (s, 1H), 6.83 – 6.79 (m, 1H), 6.77 (d, *J* = 8.6 Hz, 1H), 6.70 – 6.63 (m, 2H), 6.48 (s, 1H), 5.80 – 5.64 (m, 2H), 5.33 – 5.24 (m, 1H), 4.78 – 4.65 (m, 1H), 4.47 – 4.39 (m, 2H), 4.29 – 4.18 (m, 1H), 3.87 – 3.81 (m, 6H), 3.53 – 3.25 (m, 4H), 3.24 – 3.11 (m, 2H), 2.68 – 2.46 (m, 2H), 2.36 (d, *J* = 13.7 Hz, 1H), 2.28 – 2.16 (m, 2H), 2.10 – 1.96 (m, 2H), 1.83 – 1.59 (m, 5H), 1.52 – 1.42 (m, 1H), 1.39 – 1.29 (m, 1H), 1.28 – 1.12 (m, 9H), 0.92 – 0.80 (m, 3H). $^{13}$C NMR (126 MHz, $CDCl_3$) δ 208.17, 174.17, 172.17, 172.01, 169.87, 168.84, 167.61, 166.78, 157.52, 149.04, 147.54, 141.94, 136.37, 133.43, 130.20, 127.65, 123.76, 122.21, 120.31, 119.67, 118.73, 114.45, 113.23, 111.90, 111.68, 111.51, 109.90, 67.36, 56.07, 56.00, 53.51, 51.53, 50.00, 46.87, 44.33, 38.31, 35.28, 35.12, 32.58, 31.40, 29.83, 27.36, 26.57, 25.02, 23.54, 23.34, 21.28, 17.47, 8.88. HRMS (ESI$^+$) m/z calcd for $C_{49}H_{62}N_6NaO_{11}^+$ [M+Na]$^+$: 933.4374, found 933.4303.

**JZ12:** *(R)-1-(3-(2-((3-(((S)-1-(((S)-1-amino-3-(4-fluorophenyl)-1-oxopropan-2-yl)amino)-1-oxopropan-2-yl)amino)-3-oxopropyl)amino)-2-oxoethoxy)phenyl)-3-(3,4-dimethoxyphenyl)propyl (S)-1-(3,3-dimethyl-2-oxopentanoyl)piperidine-2-carboxylate*

$^{1}H$ NMR (500 MHz, MeOD) δ 7.33 – 7.27 (m, 1H), 7.24 – 7.19 (m, 2H), 7.03 – 6.89 (m, 5H), 6.84 (d, *J* = 8.0 Hz, 1H), 6.79 – 6.75 (m, 1H), 6.73 – 6.68 (m, 1H), 5.78 – 5.71 (m, 1H), 5.26 – 5.19 (m, 1H), 4.61 – 4.53 (m, 1H), 4.50 (d, *J* = 6.4 Hz, 2H), 4.22 – 4.15 (m, 1H), 3.84 – 3.76 (m, 6H), 3.59 – 3.47 (m, 2H), 3.28 – 3.13 (m, 2H), 2.99 – 2.77 (m, 2H), 2.67 – 2.51 (m, 2H), 2.47 – 2.40 (m, 2H), 2.27 – 2.19 (m, 1H), 2.10 – 2.01 (m, 1H), 1.79 – 1.59 (m, 5H), 1.49 – 1.30 (m, 3H), 1.21 – 1.07 (m, 9H), 0.91 – 0.76 (m, 3H). $^{13}C$ NMR (126 MHz, MeOD) δ 208.93, 170.87, 168.94, 164.02, 162.09, 159.09, 150.23, 148.65, 143.12, 135.02, 131.99, 131.93, 131.87, 131.81, 130.91, 121.57, 120.91, 115.98, 115.81, 114.35, 113.39, 113.06, 79.35, 79.09, 78.83, 68.16, 56.51, 55.34, 52.69, 50.92, 50.70, 47.61, 45.57, 39.08, 37.49, 36.46, 36.08, 33.46, 32.17, 27.25, 25.80, 23.82, 23.62, 22.04, 17.53, 17.39, 9.09. HRMS (ESI$^{+}$) m/z calcd for $C_{47}H_{60}FN_5NaO_{11}^{+}$ [M+Na]$^{+}$: 912.4171, found 912.4108.

**JZ13:** *(R)-1-(3-(2-((3-(((S)-1-(((S)-1-amino-3-(4-methoxyphenyl)-1-oxopropan-2-yl)amino)-1-oxopropan-2-yl)amino)-3-oxopropyl)amino)-2-oxoethoxy)phenyl)-3-(3,4-dimethoxyphenyl)propyl (S)-1-(3,3-dimethyl-2-oxopentanoyl)piperidine-2-carboxylate*

$^{1}H$ NMR (500 MHz, MeOD) δ 7.30 (t, *J* = 7.9 Hz, 1H), 7.14 – 7.09 (m, 2H), 7.02 – 6.95 (m, 2H), 6.94 – 6.89 (m, 1H), 6.84 – 6.73 (m, 4H), 6.72 – 6.67 (m, 1H), 5.79 – 5.70 (m, 1H), 5.22 (d, *J* = 5.6 Hz, 1H), 4.59 – 4.45 (m, 3H), 4.23 – 4.17 (m, 1H), 3.80 (d, *J* = 5.9 Hz, 6H), 3.71 (s, 3H), 3.58 – 3.47 (m, 2H), 3.23 – 3.15 (m, 1H), 3.12 – 3.03 (m, 1H), 2.98 (s, 1H), 2.94 – 2.87 (m, 1H), 2.66 – 2.52 (m, 2H), 2.47 – 2.42 (m, 2H), 2.36 – 2.31 (m, 1H), 2.28 – 2.19 (m, 1H), 2.10 – 2.01 (m, 1H), 1.79 – 1.59 (m, 5H), 1.37 – 1.27 (m, 2H), 1.24 – 1.06 (m, 9H), 0.90 – 0.77 (m, 3H). $^{13}C$ NMR (126 MHz, MeOD) δ 208.90, 175.84, 174.58, 174.11, 170.83, 168.89, 159.77, 159.02, 150.17, 148.60, 143.08, 134.97, 131.58, 131.21, 130.89 (2C), 130.16 (2C), 121.54, 120.88, 115.43, 114.70, 114.34, 113.33, 113.00, 79.29, 79.03, 78.77, 77.88, 68.12, 56.40, 55.58, 52.65, 50.87, 47.58, 45.53, 39.04, 37.51, 36.50, 36.39, 33.43, 32.14, 27.23, 25.76, 23.60, 22.01, 17.46, 9.09. HRMS (ESI$^{+}$) m/z calcd for $C_{48}H_{63}N_5NaO_{12}^{+}$ [M+Na]$^{+}$: 924.4371, found 924.5178.

**JZ14:** *(R)-1-(3-(2-((3-(((S)-1-(((S)-1-amino-1-oxo-4-phenylbutan-2-yl)amino)-1-oxopropan-2-yl)amino)-3-oxopropyl)amino)-2-oxoethoxy)phenyl)-3-(3,4-dimethoxyphenyl)propyl (S)-1-(3,3-dimethyl-2-oxopentanoyl)piperidine-2-carboxylate*

$^1$H NMR (500 MHz, MeOD) δ 7.31-7.27 (m, 1H), 7.24-7.20 (m, 2H), 7.18 – 7.11 (m, 3H), 7.01-6.93 (m, 2H), 6.92-6.88 (m, 1H), 6.85-6.83 (m, 1H), 6.80-6.78 (m, 1H), 6.73-6.69 (m, 1H), 5.77-5.72 (m, 1H), 5.23-5.21 (m, 1H), 4.50-4.41 (m, 2H), 4.36-4.28 (m, 2H), 3.80 (s, 3H), 3.79 (s, 3H), 3.62-3.52 (m, 2H), 3.25-3.13 (m, 1H), 2.76-2.54 (m, 4H), 2.54-2.44 (m, 2H), 2.38-2.19 (m, 2H), 2.17-2.01 (m, 2H), 2.01-1.91 (m, 1H), 1.79-1.58 (m, 5H), 1.53-1.40 (m, 1H), 1.39-1.25 (m, 5H), 1.24-1.18 (m, 5H), 1.10 (s, 1H), 0.93-0.76 (m, 3H). $^{13}$C NMR (126 MHz, MeOD) δ 207.62, 175.35, 173.83, 172.81, 169.63, 169.61, 167.71, 157.85, 149.04, 147.46, 141.93, 141.06, 133.82, 129.59, 128.15 (2C), 128.08 (2C), 125.68, 120.31, 119.58, 114.22, 113.00, 112.20, 111.87, 76.60, 66.86, 55.16, 55.06, 52.63, 51.43, 49.58, 46.31, 44.31, 37.81, 35.30, 35.09, 33.55, 32.18, 31.67, 30.86, 25.94, 24.50, 22.44, 22.24, 20.74, 16.33, 7.71. HRMS (ESI$^+$) m/z calcd for $C_{48}H_{63}N_5NaO_{11}^+$ [M+Na]$^+$: 908.4422, found 908.4360.

**JZ15:** *(R)-1-(3-(2-((3-(((S)-1-(((S)-3-([1,1'-biphenyl]-4-yl)-1-amino-1-oxopropan-2-yl)amino)-1-oxopropan-2-yl)amino)-3-oxopropyl)amino)-2-oxoethoxy)phenyl)-3-(3,4-dimethoxyphenyl)propyl (S)-1-(3,3-dimethyl-2-oxopentanoyl)piperidine-2-carboxylate*

$^1$H NMR (500 MHz, MeOD) δ 7.56-7.52 (m, 2H), 7.51-7.49 (m, 2H), 7.39-7.36 (m, 2H), 7.32-7.26 (m, 4H), 7.02-6.94 (m, 2H), 6.93-6.90 (m, 1H), 6.85-6.81 (m, 1H), 6.79-6.74 (m, 1H), 6.71-6.67 (m, 1H), 5.76-5.71 (m, 1H), 5.23-5.19 (m, 1H), 4.66-4.61 (m, 1H), 4.50-4.45 (m, 2H), 4.26-4.17 (m, 1H), 3.79 (s, 3H), 3.78 (s, 3H), 3.59-3.44 (m, 2H), 3.26-3.13 (m, 2H), 3.05-2.98 (m, 2H), 2.65-2.52 (m, 2H), 2.46-2.40 (m, 2H), 2.36-2.30 (m, 1H), 2.26-2.19 (m, 1H), 2.08-2.01 (m, 1H), 1.73-1.61 (m, 4H), 1.31-1.26 (m, 3H), 1.24-1.22 (m, 3H), 1.21-1.12 (m, 6H), 0.89-0.78 (m, 3H). $^{13}$C NMR (126 MHz, MeOD) δ 207.64, 174.54, 173.46, 172.89, 169.61, 167.70, 163.45, 157.84, 149.00, 147.42, 141.91, 140.67, 139.45, 136.29, 133.79, 129.51 (2C), 128.47 (2C), 126.87, 126.57 (2C),

126.44 (2C), 120.31, 119.62, 114.19, 113.08, 112.16, 111.84, 78.12, 77.86, 77.60, 76.63, 66.85, 55.19, 54.03, 51.43, 49.61, 46.32, 44.30, 37.80, 36.75, 35.29, 35.17, 32.19, 30.88, 25.95, 24.51, 22.49, 20.75, 16.13, 7.76. HRMS (ESI$^+$) m/z calcd for $C_{53}H_{65}N_5NaO_{11}^+$ [M+Na]$^+$: 970.4578, found 970.4512.

**JZ16:** *(R)-1-(3-(2-((3-(((S)-1-((S)-2-carbamoylindolin-1-yl)-1-oxopropan-2-yl)amino)-3-oxopropyl)amino)-2-oxoethoxy)phenyl)-3-(3,4-dimethoxyphenyl)propyl (S)-1-(3,3-dimethyl-2-oxopentanoyl)piperidine-2-carboxylate*

$^1$H NMR (500 MHz, MeOD) δ 8.15 – 8.01 (m, 1H), 7.35 – 7.28 (m, 1H), 7.28 – 7.22 (m, 1H), 7.14 (s, 1H), 7.07 (t, *J* = 7.5 Hz, 1H), 7.04 – 6.98 (m, 2H), 6.98 – 6.93 (m, 1H), 6.92 – 6.84 (m, 1H), 6.82 – 6.76 (m, 1H), 6.76 – 6.69 (m, 1H), 5.79 – 5.72 (m, 1H), 5.25 – 5.21 (m, 1H), 5.14 – 5.00 (m, 1H), 4.63 – 4.33 (m, 3H), 3.87 – 3.76 (m, 6H), 3.68 – 3.58 (m, 1H), 3.58 – 3.45 (m, 2H), 3.25 – 3.15 (m, 1H), 2.70 – 2.41 (m, 4H), 2.38 – 2.31 (m, 1H), 2.32 – 2.16 (m, 1H), 2.13 – 1.99 (m, 1H), 1.79 – 1.59 (m, 5H), 1.54 – 1.45 (m, 2H), 1.44 – 1.36 (m, 3H), 1.35 – 1.28 (m, 2H), 1.26 – 1.08 (m, 6H), 0.95 – 0.77 (m, 3H). $^{13}$C NMR (126 MHz, MeOD) δ 209.02, 173.52, 170.98, 169.09, 168.43, 159.29, 150.43, 148.85, 143.30, 135.24, 130.98, 128.38, 125.56, 121.70, 120.99, 120.97, 119.04, 118.68, 115.57, 113.60, 113.27, 78.25, 78.02, 68.28, 62.59, 58.28, 56.56, 56.46, 52.82, 47.70, 45.71, 40.15, 39.15, 36.63, 36.12, 33.65, 33.58, 32.24, 28.59, 27.34, 25.89, 25.48, 23.84, 23.63, 22.14, 22.05, 9.16. HRMS (ESI$^+$) m/z calcd for $C_{47}H_{59}N_5NaO_{11}^+$ [M+Na]$^+$: 892.4109, found 892.4057.

**JZ17:** *(R)-1-(3-(2-((3-(((S)-1-((S)-3-carbamoyl-3,4-dihydroisoquinolin-2(1H)-yl)-1-oxopropan-2-yl)amino)-3-oxopropyl)amino)-2-oxoethoxy)phenyl)-3-(3,4-dimethoxyphenyl)propyl (S)-1-(3,3-dimethyl-2-oxopentanoyl)piperidine-2-carboxylate*

$^1$H NMR (500 MHz, MeOD) δ 7.37 – 7.27 (m, 1H), 7.26 – 7.15 (m, 4H), 7.07 – 6.97 (m, 2H), 6.95 – 6.90 (m, 1H), 6.90 – 6.83 (m, 1H), 6.83 – 6.77 (m, 1H), 6.76 – 6.68 (m, 1H), 5.80 – 5.73 (m, 1H), 5.27 – 5.21 (m, 1H), 5.07 – 4.98 (m, 1H), 4.95 – 4.89 (m, 2H), 4.68 (d, *J* = 14.9 Hz, 1H), 4.55 (s, 1H), 4.49 (s, 1H), 3.84 – 3.78 (m, 6H), 3.64 – 3.47 (m, 2H),

3.46 – 3.38 (m, 1H), 3.24 – 3.17 (m, 1H), 3.17 – 3.13 (m, 1H), 2.70 – 2.52 (m, 2H), 2.52 – 2.42 (m, 2H), 2.39 – 2.32 (m, 1H), 2.30 – 2.20 (m, 1H), 2.13 – 2.02 (m, 1H), 1.81 – 1.58 (m, 5H), 1.55 – 1.44 (m, 1H), 1.42 – 1.38 (m, 2H), 1.33 – 1.29 (m, 3H), 1.28 – 1.09 (m, 6H), 0.95 – 0.77 (m, 3H). $^{13}$C NMR (126 MHz, MeOD) δ 209.02, 176.11, 174.37, 173.16, 171.00, 169.10, 159.24, 150.44, 148.84, 143.32, 135.23, 134.76, 130.99, 129.47, 128.76, 128.62, 128.02, 127.07, 121.70, 120.97, 115.60, 114.36, 113.60, 113.27, 77.99, 68.23, 56.55, 56.45, 55.76, 52.82, 47.70, 47.29, 46.94, 45.71, 44.80, 39.23, 36.57, 36.12, 33.57, 32.57, 32.25, 27.33, 25.89, 23.84, 23.64, 22.14, 17.33, 9.11. HRMS (ESI$^+$) m/z calcd for $C_{48}H_{61}N_5NaO_{11}^+$ [M+Na]$^+$: 906.4265 found 906.4210.

**JZ18:** *(R)-1-(3-(((4S,7S)-4-benzyl-7-methyl-3,6,9,13-tetraoxo-2,5,8,12-tetraazatetradecan-14-yl)oxy)phenyl)-3-(3,4-dimethoxyphenyl)propyl (S)-1-(3,3-dimethyl-2-oxopentanoyl)piperidine-2-carboxylate*

$^1$H NMR (500 MHz, MeOD) δ 7.30-7.28 (m, 1H), 7.27-7.22 (m, 2H), 7.21-7.15 (m, 3H), 7.04-6.96 (m, 2H), 6.95-6.90 (m, 1H), 6.85-6.83 (m, 1H), 6.80-6.76 (m, 1H), 6.72-6.68 (m, 1H), 5.78-5.71 (m, 1H), 5.26-5.20 (m, 1H), 5.03-4.97 (m, 1H), 4.51 (s, 2H), 4.37-4.29 (m, 1H), 3.79 (s, 3H), 1.78 (s, 3H), 3.61-3.48 (m, 2H), 3.02-2.88 (m, 2H), 2.80-2.73 (m, 3H), 2.74-2.71 (m, 3H), 2.68-2.51 (m, 2H), 2.50-2.43 (m, 2H), 2.37-2.18 (m, 2H), 2.09-2.01 (m, 1H), 1.80-1.57 (m, 5H), 1.53-1.40 (m, 1H), 1.29-1.25 (m, 3H), 1.23-1.09 (m, 6H), 0.90-0.75 (m, 3H). $^{13}$C NMR (126 MHz, MeOD) δ 207.62, 172.98, 172.22, 171.58, 169.58, 167.68, 157.92, 149.06, 147.47, 141.97, 136.45, 133.81, 129.61, 129.11 (2C), 128.09 (2C), 126.62, 120.30, 119.60, 114.28, 112.99, 112.21, 111.88, 76.57, 66.95, 55.17, 55.07, 51.42, 50.48, 48.99, 46.31, 44.30, 38.02, 37.85, 37.50, 36.03, 35.24, 34.83, 34.65, 32.19, 30.87, 25.96, 24.52, 22.47, 22.27, 16.49, 7.75. HRMS (ESI$^+$) m/z calcd for $C_{48}H_{63}N_5NaO_{11}^+$ [M+Na]$^+$: 908.4422 found 908.4372.

**JZ19:** *(R)-1-(3-(((4S,7S)-4-benzyl-2,7-dimethyl-3,6,9,13-tetraoxo-2,5,8,12-tetraazatetradecan-14-yl)oxy)phenyl)-3-(3,4-dimethoxyphenyl)propyl (S)-1-(3,3-dimethyl-2-oxopentanoyl)piperidine-2-carboxylate*

$^{1}H$ NMR (500 MHz, MeOD) δ 7.32-7.28 (m, 1H), 7.27-7.22 (m, 2H), 7.21-7.15 (m, 3H), 7.03-6.96 (m, 2H), 6.95-6.90 (m, 1H), 6.86-6.82 (m, 1H), 6.80-6.76 (m, 1H), 6.72-6.68 (m, 1H), 5.76-5.71 (m, 1H), 5.26-5.20 (m, 1H), 4.02-4.96 (m, 1H), 4.51 (s, 2H), 4.36-4.30 (m, 1H), 3.79 (s, 3H), 3.78 (s, 3H), 3.60-3.48 (m, 2H), 3.22-3.14 (m, 1H), 3.01-2.89 (m, 2H), 2.81 (s, 1H), 2.80-2.72 (m, 6H), 2.68-2.52 (m, 2H), 2.50-2.45 (m, 2H), 2.38-2.17 (m, 2H), 2.12-1.98 (m, 1H), 1.78-1.57 (m, 5H), 1.51-1.29 (m, 2H), 1.32-1.25 (m, 3H), 1.23-1.10 (m, 6H), 0.91-0.77 (m, 3H). $^{13}C$ NMR (126 MHz, MeOD) δ 207.62, 172.98, 172.22, 171.58, 169.58, 167.68, 157.92, 149.06, 147.47, 141.97, 136.45, 133.81, 129.61, 129.11 (2C), 128.09 (2C), 126.62, 120.30, 119.60, 114.28, 112.99, 112.21, 111.88, 76.57, 66.95, 55.17, 55.07, 51.42, 50.48, 48.99, 46.31, 44.30, 38.02, 37.85, 37.50, 36.03, 35.24, 34.83, 34.65, 32.19, 30.87, 25.96, 24.52, 22.47, 22.27, 20.77, 16.49, 7.75. HRMS (ESI$^+$) m/z calcd for $C_{49}H_{65}N_5NaO_{11}^+$ [M+Na]$^+$: 922.4578, found 922.4534.

**Synthesis of JZ20**

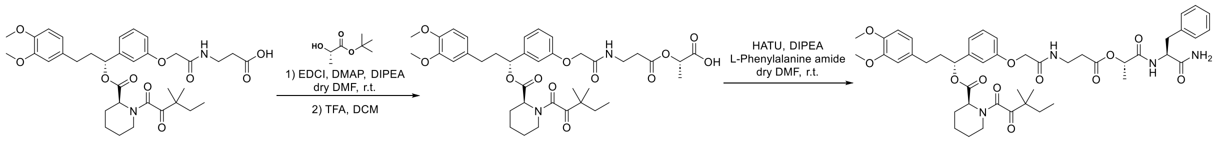


**JZ20:** *(R)-1-(3-(2-((3-(((S)-1-(((S)-1-amino-1-oxo-3-phenylpropan-2-yl)amino)-1-oxopropan-2-yl)oxy)-3-oxopropyl)amino)-2-oxoethoxy)phenyl)-3-(3,4-dimethoxyphenyl)propyl (S)-1-(3,3-dimethyl-2-oxopentanoyl)piperidine-2-carboxylate*

S-4 was synthesized according to the previously reported method[1]. S-4 (81.4 mg, 0.124 mmol, 1.0 eq), DMAP (1.52 mg, 0.01 mmol, 0.1 eq), and *tert*-butyl (*S*)-2-hydroxypropanoate (23.6 mg, 0.162 mmol, 1.3 eq) were dissolved in dry DMF (3.0 mL) at room temperature. EDCI (31.0 mg, 0.162 mmol, 1.3 eq) was added to the reaction and stirred for overnight. Upon completion, the reaction was concentrated, extracted with DCM (30 mL) and washed with brine (1 x 30 mL). The organic layer was dried by Na2SO4, concentrated, and the resulting residue was used without further purification.

The crude product was dissolved in DCM (5.0 mL), and TFA (1.0 mL) was added to the reaction and stirred for 2 h. The reaction was concentrated used without further purifications as S-5. S-5 (80.0 mg, 0.11 mmol, 1eq), HATU (54.4 mg, 0.14 mmol, 1.3 eq), and L-Phenylalanine amide (26.2 mg, 0.22 mmol, 2 eq) were dissolved in dry DMF (2.0 mL) at room temperature. DIPEA (95.0 µL, 0.55 mmol, 5.0 eq) was added to the reaction and stirred for 2 hours. The reaction was then concentrated and extracted with DCM (30 mL) and brine (30 mL). The organic layer was dried with anhydrous sodium sulfate. The reaction mixture was then concentrated and purified by flash chromatography to provide JZ20 as a colorless oil (53.1 mg, 0.06 mmol, 48.4% yield over 3 steps).

$^{1}$H NMR (500 MHz, MeOD) δ 7.96 (s, 1H), 7.30 (t, *J* = 7.9 Hz, 1H), 7.26 – 7.15 (m, 5H), 7.02 – 6.94 (m, 2H), 6.93 – 6.89 (m, 1H), 6.83 (d, *J* = 8.1 Hz, 1H), 6.78 – 6.74 (m, 1H), 6.72 – 6.68 (m, 1H), 5.74 (s, 1H), 5.03 – 4.97 (m, 1H), 4.66 – 4.57 (m, 1H), 4.50 (s, 2H), 3.80 (d, *J* = 6.6 Hz, 6H), 3.74 – 3.64 (m, 1H), 3.63 – 3.50 (m, 2H), 3.23 – 3.15 (m, 3H), 2.70 – 2.52 (m, 4H), 2.38 – 2.19 (m, 2H), 2.11 – 2.00 (m, 1H), 1.80 – 1.43 (m, 6H), 1.26 (d, *J* = 7.0 Hz, 4H), 1.22 – 1.05 (m, 6H), 0.83 (dt, *J* = 42.0, 7.4 Hz, 3H). $^{13}$C NMR (126 MHz, MeOD) δ 208.96, 175.41, 172.39, 170.85, 168.88, 164.64, 159.02, 150.16, 148.59, 143.08, 138.02, 134.97, 130.31 (2C), 129.30 (2C), 127.69, 121.54, 115.42, 114.30, 113.33, 113.00, 79.29, 79.04, 78.78, 77.88, 71.70, 68.11, 55.70, 55.01, 54.64, 52.65, 47.59, 45.53, 43.68, 39.05, 38.61, 35.84, 35.00, 33.44, 32.15, 27.24, 25.77, 22.02, 17.87, 13.14, 9.13. HRMS (ESI$^{+}$) m/z calcd for $C_{47}H_{60}N_4NaO_{12}^{+}$ [M+Na]$^{+}$: 895.4105, found 895.4066.

**Synthesis of JZ21-JZ25, JZ27**

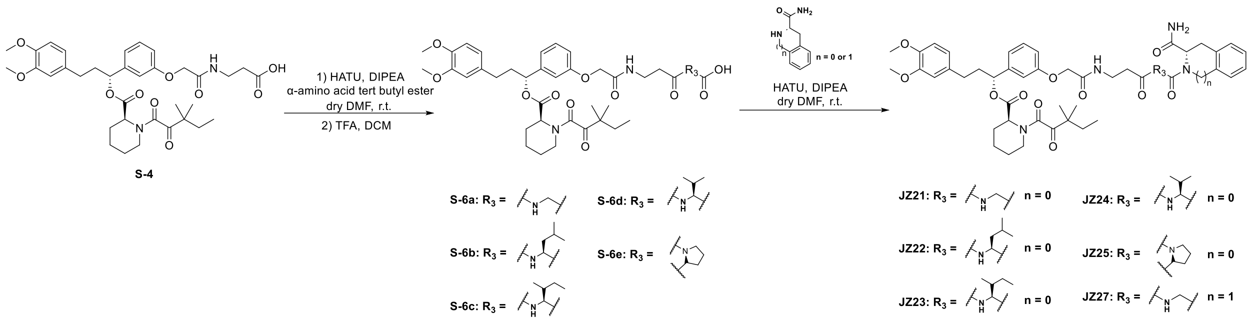

**JZ21:** *(R)-1-(3-(2-((3-((2-((S)-2-carbamoylindolin-1-yl)-2-oxoethyl)amino)-3-oxopropyl)amino)-2-oxoethoxy)phenyl)-3-(3,4-dimethoxyphenyl)propyl (S)-1-(3,3-dimethyl-2-oxopentanoyl)piperidine-2-carboxylate*

Step 1: S-4 (80.0 mg, 0.12 mmol, 1.0 eq), HATU (60.4 mg, 0.16 mmol, 1.3 eq), and tert-butyl glycinate hydrochloride (41.0 mg, 0.24 mmol, 2 eq) were dissolved in dry DMF (2.0 mL) at room temperature. DIPEA (105 µL, 0.61 mmol, 5.0 eq) was added to the reaction and stirred for 2 hours. The reaction was then concentrated and extracted with DCM (30 mL) and brine (30 mL). The organic layer was dried with anhydrous sodium sulfate. The concentrated resulting residue was dissolved in DCM (5.0 mL), and TFA (1.0 mL) was added and stirred for 2 hours. The reaction mixture was then concentrated and purified by flash chromatography to provide S-6a as a colorless oil (53.7 mg, 0.08 mmol, 66.7% over 2 steps).

$^1$H NMR (500 MHz, $CDCl_3$) δ 7.55 (s, 1H), 7.26 – 7.23 (m, 1H), 7.10 (s, 1H), 6.99 – 6.89 (m, 2H), 6.85 – 6.74 (m, 2H), 6.72 – 6.62 (m, 2H), 5.79 – 5.70 (m, 1H), 5.28 (s, 1H), 4.48 (s, 2H), 3.95 (s, 2H), 3.87 – 3.81 (m, 6H), 3.63 (s, 2H), 3.47 (s, 1H), 3.35 (d, *J* = 13.1 Hz, 1H), 3.24 – 3.15 (m, 1H), 2.68 – 2.41 (m, 4H), 2.37 (d, *J* = 13.4 Hz, 1H), 2.27 – 2.17 (m, 1H), 2.10 – 1.98 (m, 1H), 1.82 – 1.55 (m, 5H), 1.43 – 1.34 (m, 2H), 1.22 – 1.08 (m, 6H), 0.90 – 0.75 (m, 3H).

Step 2: S-6a (26.3 mg, 0.04 mmol, 1 eq), HATU (18.3 mg, 0.05 mmol, 1.3 eq), and (2S)-Indoline-2-carboxamide (6.0 mg, 0.04 mmol, 1 eq) were dissolved in dry DMF (2.0 mL) at room temperature. DIPEA (19.1 µL, 0.11 mmol, 3.0 eq) was added to the reaction and stirred for 2 hours. The reaction was then concentrated and extracted with DCM (30 mL) and brine (30 mL). The organic layer was dried with anhydrous sodium sulfate. The reaction mixture was then concentrated and purified by flash chromatography to provide JZ21 as a white solid (22.3 mg, 0.03 mmol, 70.5% yield).

$^1$H NMR (500 MHz, $CDCl_3$) δ 8.11 (s, 1H), 7.52 – 7.31 (m, 1H), 7.21 – 7.02 (m, 3H), 6.98 – 6.89 (m, 2H), 6.85 – 6.47 (m, 6H), 5.83 – 5.67 (m, 1H), 5.50 (s, 1H), 5.29 (d, *J* = 8.9 Hz, 1H), 5.14 (s, 1H), 4.85 (s, 1H), 4.49 (s, 3H), 4.22 – 3.90 (m, 1H), 3.84 (d, *J* = 6.1 Hz, 6H), 3.76 – 3.54 (m, 3H), 3.49 – 3.25 (m, 2H), 3.22 – 3.10 (m, 1H), 2.68 – 2.44 (m, 4H), 2.36 (d, *J* = 13.8 Hz, 1H), 2.28 – 2.14 (m, 1H), 2.10 – 1.95 (m, 1H), 1.72 – 1.60 (m, 6H), 1.28 – 1.14 (m, 7H), 0.86 (t, *J* = 7.4 Hz, 3H). $^{13}$C NMR (126 MHz, $CDCl_3$) δ 208.16, 169.93, 168.59, 167.54, 157.63, 149.05, 147.52, 141.96, 133.49, 130.11, 128.03, 125.11, 120.31, 120.01, 111.89, 111.50, 77.41, 77.16, 76.91, 67.48, 56.08, 56.01, 55.62, 51.50, 46.85, 44.33, 43.60, 38.32, 35.69, 35.29, 32.67, 32.58, 31.41, 29.84, 27.82, 26.60, 25.06, 24.58, 23.55, 23.29, 21.32, 12.54, 8.94, 8.88. HRMS (ESI$^+$) m/z calcd for $C_{46}H_{57}N_5NaO_{11}^+$ [M+Na]$^+$: 878.3952, found 878.3901.

The synthetic route of JZ22-JZ25, JZ27 were similar to JZ21.

**JZ22:** *(R)-1-(3-(2-((3-(((S)-1-((S)-2-carbamoylindolin-1-yl)-4-methyl-1-oxopentan-2-yl)amino)-3-oxopropyl)amino)-2-oxoethoxy)phenyl)-3-(3,4-dimethoxyphenyl)propyl (S)-1-(3,3-dimethyl-2-oxopentanoyl)piperidine-2-carboxylate*

$^1$H NMR (500 MHz, $CDCl_3$) δ 8.11 – 8.00 (m, 1H), 7.44 (d, *J* = 25.7 Hz, 1H), 7.19 (d, *J* = 7.2 Hz, 1H), 7.14 – 7.03 (m, 2H), 7.00 – 6.89 (m, 2H), 6.84 (s, 1H), 6.79 – 6.75 (m, 1H), 6.72 – 6.45 (m, 3H), 5.83 – 5.66 (m, 1H), 5.45 – 5.23 (m, 1H), 4.78 (s, 1H), 4.63 – 4.37 (m, 3H), 3.93 – 3.82 (m, 6H), 3.78 – 3.47 (m, 4H), 3.41 – 3.31 (m, 1H), 3.18 (s, 1H), 2.65 – 2.44 (m, 4H), 2.35 (d, *J* = 13.8 Hz, 1H), 2.26 – 2.15 (m, 1H), 2.09 – 1.97 (m, 1H), 1.91 – 1.54 (m, 13H), 1.21 (d, *J* = 7.7 Hz, 5H), 1.15 – 1.09 (m, 1H), 1.01 – 0.77 (m, 9H). $^{13}$C NMR (126 MHz, $CDCl_3$) δ 208.03, 169.78, 167.50, 157.68, 149.05, 147.54, 141.96, 141.66, 133.50, 130.15, 129.95, 127.82, 125.23, 124.81, 120.29, 118.26, 114.39,

113.51, 111.88, 111.47, 67.61, 61.67, 56.83, 56.08, 56.00, 51.67, 51.47, 46.85, 44.30, 41.10, 39.03, 38.28, 35.95, 35.37, 34.22, 32.70, 32.61, 31.41, 29.84, 26.57, 25.13, 25.06, 23.72, 23.59, 23.55, 23.31, 21.61, 21.30, 14.26, 8.90. HRMS (ESI$^+$) m/z calcd for $C_{50}H_{65}N_5NaO_{11}^+$ [M+Na]$^+$: 934.4578, found 934.4529.

**JZ23:** *(1R)-1-(3-(2-((3-(((3R)-1-((S)-2-carbamoylindolin-1-yl)-3-methyl-1-oxopentan-2-yl)amino)-3-oxopropyl)amino)-2-oxoethoxy)phenyl)-3-(3,4-dimethoxyphenyl)propyl (2S)-1-(3,3-dimethyl-2-oxopentanoyl)piperidine-2-carboxylate*

$^1$H NMR (500 MHz, $CDCl_3$) δ 8.16 – 8.09 (m, 1H), 7.51 – 7.39 (m, 1H), 7.22 – 7.14 (m, 2H), 7.11 – 7.04 (m, 1H), 6.99 – 6.87 (m, 2H), 6.83 (d, *J* = 8.2 Hz, 1H), 6.78 (d, *J* = 1.9 Hz, 1H), 6.70 – 6.64 (m, 2H), 6.43 – 6.34 (m, 1H), 5.82 – 5.71 (m, 1H), 5.50 – 5.11 (m, 2H), 4.94 (d, *J* = 9.1 Hz, 1H), 4.58 – 4.37 (m, 3H), 3.88 – 3.83 (m, 6H), 3.74 – 3.58 (m, 2H), 3.52 – 3.32 (m, 3H), 3.20 (d, *J* = 10.6 Hz, 1H), 2.67 – 2.44 (m, 4H), 2.37 (d, *J* = 13.9 Hz, 1H), 2.21 (s, 1H), 2.11 – 1.98 (m, 1H), 1.65 – 1.54 (m, 8H), 1.21 (s, 6H), 1.14 – 1.10 (m, 2H), 1.03 – 0.95 (m, 2H), 0.91 – 0.69 (m, 9H). $^{13}$C NMR (126 MHz, $CDCl_3$) δ 208.02, 172.85, 172.02, 170.55, 169.73, 168.66, 167.62, 157.72, 149.07, 147.56, 141.91, 141.78, 133.49, 130.17, 127.88, 125.18, 124.83, 120.29, 118.29, 114.22, 113.67, 111.88, 111.48, 67.74, 61.90, 57.02, 56.09, 56.01, 51.52, 46.86, 44.30, 38.76, 38.25, 38.07, 36.13, 35.47, 34.11, 32.60, 31.43, 29.85, 26.62, 25.06, 24.59, 23.58, 23.33, 21.32, 15.89, 14.26, 11.47, 8.90. HRMS (ESI$^+$) m/z calcd for $C_{50}H_{65}N_5NaO_{11}^+$ [M+Na]$^+$: 934.4578, found 934.4533.

**JZ24:** *(1R)-1-(3-(2-((3-((1-((S)-2-carbamoylindolin-1-yl)-3-methyl-1-oxobutan-2-yl)amino)-3-oxopropyl)amino)-2-oxoethoxy)phenyl)-3-(3,4-dimethoxyphenyl)propyl (2S)-1-(3,3-dimethyl-2-oxopentanoyl)piperidine-2-carboxylate*

$^1$H NMR (500 MHz, $CDCl_3$) δ 8.12 (d, *J* = 7.9 Hz, 1H), 7.52 – 7.38 (m, 1H), 7.26 – 7.11 (m, 5H), 7.10 – 7.04 (m, 1H), 6.98 – 6.86 (m, 2H), 6.85 – 6.75 (m, 2H), 6.70 – 6.64 (m, 2H), 6.52 (s, 1H), 5.79 – 5.71 (m, 1H), 5.35 – 5.24 (m, 1H), 4.99 – 4.92 (m, 1H), 4.58 – 4.36 (m, 3H), 3.88 – 3.83 (m, 6H), 3.76 – 3.57 (m, 2H), 3.54 – 3.29 (m, 3H), 2.57 – 2.48

(m, 3H), 2.41 – 2.33 (m, 1H), 2.12 – 1.97 (m, 2H), 1.85 – 1.55 (m, 8H), 1.28 – 1.17 (m, 8H), 1.05 – 0.92 (m, 6H), 0.91 – 0.84 (m, 3H). $^{13}$C NMR (126 MHz, $CDCl_3$) δ 208.03, 172.95, 172.08, 170.39, 169.75, 168.69, 167.59, 157.69, 149.06, 147.54, 141.90, 141.78, 133.49, 130.15, 130.07, 127.87, 124.83, 120.29, 118.22, 114.22, 113.65, 111.87, 111.48, 67.69, 61.88, 57.53, 56.85, 56.08, 56.00, 51.50, 46.85, 44.29, 38.24, 36.11, 35.48, 34.12, 32.60, 31.46, 31.41, 29.84, 26.60, 25.05, 23.58 (2C), 23.31, 21.31, 19.75, 17.92, 8.89. HRMS (ESI$^+$) m/z calcd for $C_{49}H_{63}N_5NaO_{11}^+$ [M+Na]$^+$: 920.4422, found 920.4373.

**JZ25:** *(R)-1-(3-(2-((3-((R)-2-((S)-2-carbamoylindoline-1-carbonyl)pyrrolidin-1-yl)-3-oxopropyl)amino)-2-oxoethoxy)phenyl)-3-(3,4-dimethoxyphenyl)propyl (S)-1-(3,3-dimethyl-2-oxopentanoyl)piperidine-2-carboxylate*

$^1$H NMR (500 MHz, $CDCl_3$) δ 8.01 (d, *J* = 8.0 Hz, 1H), 7.48 – 7.39 (m, 1H), 7.20 (d, *J* = 7.4 Hz, 1H), 7.13 (t, *J* = 7.7 Hz, 1H), 7.06 (t, *J* = 7.3 Hz, 1H), 7.00 – 6.81 (m, 4H), 6.79 – 6.60 (m, 4H), 5.80 – 5.69 (m, 1H), 5.34 – 5.24 (m, 2H), 4.87 – 4.74 (m, 1H), 4.57 – 4.39 (m, 3H), 3.86 – 3.83 (m, 6H), 3.74 – 3.45 (m, 7H), 3.43 – 3.26 (m, 3H), 2.80 – 2.43 (m, 5H), 2.40 – 2.09 (m, 6H), 2.08 – 1.87 (m, 6H), 1.28 – 1.21 (m, 5H), 0.87 (t, *J* = 7.3 Hz, 3H). $^{13}$C NMR (126 MHz, $CDCl_3$) δ 207.86, 170.78, 169.72, 168.22, 167.27, 157.54, 148.91, 147.38, 141.85, 133.40, 130.00, 127.60, 125.10, 124.65, 120.15, 119.93, 118.28, 114.57, 113.11, 111.74, 111.32, 67.45, 61.65, 59.43, 57.91, 55.94, 55.86, 53.43, 51.27, 47.86, 46.70, 46.32, 46.29, 44.15, 38.13, 34.64, 34.08, 32.47, 31.26, 29.71, 28.95, 26.47, 26.40, 24.95, 23.46, 23.15, 21.20, 14.13, 8.76. HRMS (ESI$^+$) m/z calcd for $C_{49}H_{61}N_5NaO_{11}^+$ [M+Na]$^+$: 918.4265, found 918.4201.

**JZ27:** *(R)-1-(3-(2-((3-((2-((S)-3-carbamoyl-3,4-dihydroisoquinolin-2(1H)-yl)-2-oxoethyl)amino)-3-oxopropyl)amino)-2-oxoethoxy)phenyl)-3-(3,4-dimethoxyphenyl)propyl (S)-1-(3,3-dimethyl-2-oxopentanoyl)piperidine-2-carboxylate*

$^1$H NMR (500 MHz, MeOD) δ 7.33 – 7.26 (m, 1H), 7.23 – 7.09 (m, 4H), 7.02 – 6.90 (m, 3H), 6.86 – 6.82 (m, 1H), 6.80 – 6.75 (m, 1H), 6.73 – 6.68 (m, 1H), 5.78 – 5.71 (m, 1H), 5.24 – 5.18 (m, 1H), 5.06 – 5.01 (m, 1H), 4.81 – 4.62 (m, 2H), 4.60 – 4.48 (m, 2H), 4.32 – 4.23 (m, 1H), 4.15 (s, 1H), 3.83 – 3.74 (m, 6H), 3.58 (t, *J* = 6.6 Hz, 2H), 3.41 – 3.36

(m, 1H), 3.27 – 3.14 (m, 2H), 3.07 (dd, $J$ = 15.4, 6.0 Hz, 1H), 2.69 – 2.47 (m, 4H), 2.38 – 2.17 (m, 2H), 2.13 – 1.98 (m, 1H), 1.80 – 1.56 (m, 5H), 1.51 – 1.26 (m, 2H), 1.24 – 1.18 (m, 5H), 1.10 (s, 1H), 0.92 – 0.74 (m, 3H). $^{13}C$ NMR (126 MHz, MeOD) δ 209.05, 175.95, 174.36, 171.01, 170.94, 169.07, 159.28, 150.42, 148.83, 143.30, 135.20, 134.65, 133.83, 131.00, 129.11, 128.57, 128.01, 127.28, 121.69, 120.93, 115.61, 114.38, 113.58, 113.24, 78.01, 68.26, 58.27, 56.54, 56.44, 55.31, 52.81, 47.69, 46.19, 45.70, 45.19, 42.74, 39.20, 36.66, 36.28, 33.56, 32.25, 27.32, 25.88, 23.83, 23.63, 22.13, 9.11. HRMS (ESI$^+$) m/z calcd for $C_{47}H_{59}N_5NaO_{11}^+$ [M+Na]$^+$: 892.4109, found 892.4069.

**Synthesis of JZ26, JZ28, JZ29**

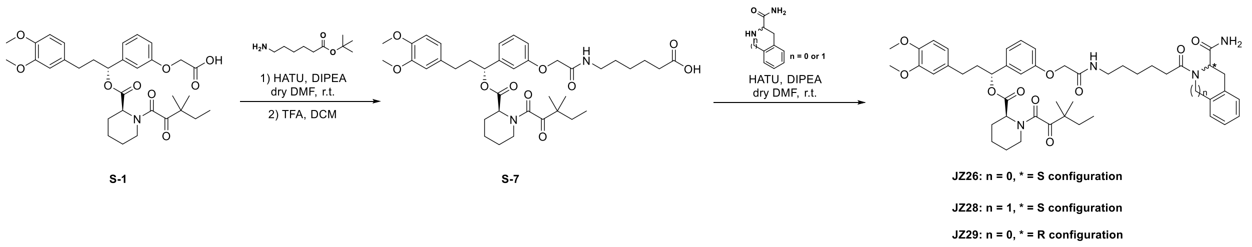


**JZ26:** *(R)-1-(3-(2-((6-((S)-2-carbamoylindolin-1-yl)-6-oxohexyl)amino)-2-oxoethoxy)phenyl)-3-(3,4-dimethoxyphenyl)propyl (S)-1-(3,3-dimethyl-2-oxopentanoyl)piperidine-2-carboxylate*

Step 1: S-1 (80.0 mg, 0.14 mmol, 1.0 eq), HATU (67.8 mg, 0.18 mmol, 1.3 eq), and tert-butyl 6-aminohexanoate (38.5 mg, 0.21 mmol, 1.5 eq) were dissolved in dry DMF (10 mL) at room temperature. DIPEA (118 μL, 0.68 mmol, 5.0 eq) was added to the reaction and stirred for 2 hours. The reaction was then concentrated and extracted with DCM (30 mL) and brine (30 mL). The organic layer was dried with anhydrous sodium sulfate. The concentrated resulting residue was dissolved in DCM (10.0 mL), and TFA (2.0 mL) was added and stirred for 2 hours. The reaction mixture was then concentrated and purified by flash chromatography to provide S-7 as a colorless oil (68.3 mg, 0.10 mmol, 71.5% over 2 steps).

$^{1}H$ NMR (500 MHz, MeOD) δ 7.35 – 7.27 (m, 1H), 7.03 – 6.96 (m, 2H), 6.94 (d, *J* = 8.5 Hz, 1H), 6.85 (d, *J* = 8.2 Hz, 1H), 6.81 – 6.76 (m, 1H), 6.74 – 6.68 (m, 1H), 5.78 – 5.72 (m, 1H), 5.24 – 5.21 (m, 1H), 4.51 (s, 2H), 3.83 – 3.77 (m, 6H), 3.26 (t, *J* = 7.1 Hz, 2H), 3.23 – 3.15 (m, 1H), 2.87 – 2.80 (m, 1H), 2.69 – 2.54 (m, 2H), 2.35 (d, *J* = 13.9 Hz, 1H), 2.30 – 2.21 (m, 3H), 2.13 – 2.00 (m, 1H), 1.80 – 1.51 (m, 9H), 1.45 – 1.27 (m, 4H), 1.24 – 1.07 (m, 6H), 0.93 – 0.77 (m, 3H).

Step 2: S-7 (47.0 mg, 0.07 mmol, 1 eq), HATU (30.8 mg, 0.08 mmol, 1.2 eq), and (2S)-Indoline-2-carboxamide (12.0 mg, 0.07 mmol, 1.1 eq) were dissolved in dry DMF (5.0 mL) at room temperature. DIPEA (58.2 µL, 0.33 mmol, 5.0 eq) was added to the reaction and stirred for 2 hours. The reaction was then concentrated and extracted with DCM (30 mL) and brine (30 mL). The organic layer was dried with anhydrous sodium sulfate. The reaction mixture was then concentrated and purified by flash chromatography to provide JZ26 as a white solid (36.4 mg, 0.04 mmol, 64.2 % yield).

$^{1}H$ NMR (500 MHz, MeOD) δ 8.13 (d, *J* = 8.1 Hz, 1H), 7.29 (t, *J* = 7.9 Hz, 1H), 7.22 – 7.12 (m, 2H), 7.04 – 6.95 (m, 3H), 6.94 – 6.90 (m, 1H), 6.86 – 6.80 (m, 1H), 6.79 – 6.75 (m, 1H), 6.73 – 6.65 (m, 1H), 5.74 (s, 1H), 5.22 (d, *J* = 5.7 Hz, 1H), 5.13 – 4.95 (m, 1H), 4.55 – 4.44 (m, 2H), 3.83 – 3.76 (m, 6H), 3.43 – 3.34 (m, 1H), 3.30 – 3.24 (m, 2H), 3.22 – 3.04 (m, 2H), 2.69 – 2.37 (m, 3H), 2.36 – 2.31 (m, 1H), 2.30 – 2.14 (m, 2H), 2.11 – 2.01 (m, 1H), 1.80 – 1.60 (m, 7H), 1.58 – 1.43 (m, 3H), 1.39 – 1.23 (m, 4H), 1.23 – 1.07 (m, 6H), 0.93 – 0.76 (m, 3H). $^{13}C$ NMR (126 MHz, MeOD) δ 208.95, 173.79, 170.84, 170.71, 168.92, 159.15, 150.22, 148.64, 143.11, 135.01, 130.87, 128.37, 125.39, 125.09, 121.57, 120.93, 115.48, 114.18, 113.38, 113.04, 79.34, 79.08, 78.82, 77.87, 68.19, 62.53, 58.11, 56.51, 56.42, 52.68, 47.61, 45.57, 39.82, 39.10, 35.86, 35.54, 33.47, 32.15, 30.07, 28.50, 27.24, 25.79, 25.19, 23.82, 23.64, 22.03, 9.10. HRMS ($ESI^+$) m/z calcd for $C_{47}H_{60}N_4NaO_{10}^+$ $[M+Na]^+$: 863.4207, found 863.4160.

The synthetic route of JZ28, JZ29 were similar to JZ26.

**JZ28:** *(R)-1-(3-(2-((6-((S)-3-carbamoyl-3,4-dihydroisoquinolin-2(1H)-yl)-6-oxohexyl)amino)-2-oxoethoxy)phenyl)-3-(3,4-dimethoxyphenyl)propyl (S)-1-(3,3-dimethyl-2-oxopentanoyl)piperidine-2-carboxylate*

$^1$H NMR (500 MHz, MeOD) δ 7.30 (s, 1H), 7.23 – 7.16 (m, 4H), 7.02 – 6.96 (m, 2H), 6.95 – 6.91 (m, 1H), 6.84 (d, *J* = 8.2 Hz, 1H), 6.80 – 6.76 (m, 1H), 6.72 – 6.69 (m, 1H), 5.74 (s, 1H), 5.22 (d, *J* = 5.7 Hz, 1H), 4.99 (t, *J* = 5.4 Hz, 1H), 4.68 (d, *J* = 6.0 Hz, 1H), 4.51 (s, 2H), 3.82 – 3.77 (m, 6H), 3.42 – 3.37 (m, 1H), 3.30 – 3.24 (m, 2H), 3.22 – 3.15 (m, 2H), 3.13 – 2.85 (m, 1H), 2.68 – 2.18 (m, 6H), 2.13 – 1.99 (m, 1H), 1.81 – 1.44 (m, 10H), 1.42 – 1.26 (m, 4H), 1.23 – 1.08 (m, 6H), 0.93 – 0.76 (m, 3H). $^{13}$C NMR (126 MHz, MeOD) δ 209.04, 175.71, 170.97, 169.08, 159.36, 150.43, 148.84, 143.33, 135.19, 134.82, 134.72, 130.96, 128.91, 128.50, 127.96, 127.15, 121.69, 121.00, 115.62, 114.22, 113.58, 113.25, 77.96, 68.31, 56.55, 56.44, 55.22, 52.81, 47.69, 47.23, 45.69, 45.04, 39.92, 39.24, 34.58, 33.57, 32.62, 32.25, 30.23, 27.51, 27.33, 25.89, 25.49, 23.83, 23.65, 22.13, 9.12. HRMS (ESI$^+$) m/z calcd for $C_{48}H_{62}N_4NaO_{10}^+$ [M+Na]$^+$: 877.4364, found 877.4319.

**JZ29:** *(R)-1-(3-(2-((6-((R)-2-carbamoylindolin-1-yl)-6-oxohexyl)amino)-2-oxoethoxy)phenyl)-3-(3,4-dimethoxyphenyl)propyl (S)-1-(3,3-dimethyl-2-oxopentanoyl)piperidine-2-carboxylate*

$^1$H NMR (500 MHz, MeOD) δ 8.19 – 8.11 (m, 1H), 7.34 – 7.12 (m, 3H), 7.06 – 6.95 (m, 3H), 6.94 – 6.88 (m, 1H), 6.87 – 6.82 (m, 1H), 6.80 – 6.77 (m, 1H), 6.74 – 6.66 (m, 1H), 5.78 – 5.69 (m, 1H), 5.24 – 5.11 (m, 1H), 5.05 – 4.97 (m, 1H), 4.65 – 4.56 (m, 1H), 4.50 (d, *J* = 8.9 Hz, 2H), 3.83 – 3.76 (m, 6H), 3.71 – 3.42 (m, 2H), 3.30 – 3.09 (m, 3H), 2.69 – 2.40 (m, 3H), 2.37 – 2.14 (m, 3H), 2.10 – 1.89 (m, 2H), 1.83 – 1.59 (m, 7H), 1.42 – 1.26 (m, 6H), 1.23 – 1.17 (m, 5H), 0.93 – 0.85 (m, 3H). $^{13}$C NMR (126 MHz, MeOD) δ 209.04, 173.98, 170.98, 169.11, 166.42, 159.36, 150.44, 148.86, 143.33, 135.22, 130.97, 128.43, 121.71, 121.02, 118.36, 115.64, 113.59, 113.27, 87.02, 77.97, 74.15, 68.32, 62.66, 60.57, 56.56, 54.80, 52.83, 47.70, 45.72, 44.17, 43.66, 39.90, 39.23, 35.96, 35.65, 33.58, 32.24, 31.08, 30.19, 29.89, 27.38, 26.01, 25.18, 23.64, 22.14,

20.93, 9.10. HRMS (ESI+) m/z calcd for $C_{47}H_{60}N_4NaO_{10}^+$ [M+Na]+: 863.4207, found 863.4162.

**Synthesis of JZ30, JZ31**

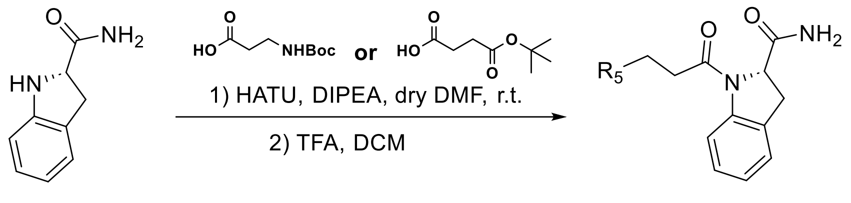


**JZ30: R$_5$ = NH$_2$**

**JZ31: R$_5$ = COOH**

**JZ30:** *(S)-1-(3-aminopropanoyl)indoline-2-carboxamide*

(2S)-Indoline-2-carboxamide (50.0 mg, 0.31 mmol, 1 eq), HATU (152 mg, 0.40 mmol, 1.3 eq), and 3-((*tert*-butoxycarbonyl)amino)propanoic acid (117 mg, 0.62 mmol, 2 eq) were dissolved in dry DMF (6.0 mL) at room temperature. DIPEA (287 µL, 1.54 mmol, 5.0 eq) was added to the reaction and stirred for 2 hours. The reaction was then concentrated and extracted with DCM (30 mL) and brine (30 mL). The organic layer was dried with anhydrous sodium sulfate. The concentrated resulting residue was dissolved in DCM (10.0 mL), and TFA (2.0 mL) was added and stirred for 2 hours. The reaction mixture was then concentrated and purified by flash chromatography to provide JZ30 as a white solid (67.3 mg, 0.29 mmol, 93 % yield).

$^1$H NMR (500 MHz, Methanol-*d*$_4$) δ 8.09 (d, *J* = 8.0 Hz, 1H), 7.29-7.18 (m, 2H), 7.08 (t, *J* = 7.5 Hz, 1H), 5.08 (d, *J* = 10.0 Hz, 1H), 4.56 (q, *J* = 7.0 Hz, 1H), 3.73-3.65 (m, 1H), 3.23-3.14 (m, 2H), 2.69-2.62 (m, 2H). HRMS (ESI+) m/z calcd for $C_{12}H_{16}N_3O_2^+$ [M+H]+: 234.1243, found 234.1243.

The synthetic route of JZ31 was similar to JZ30.

**JZ31:** *(S)-4-(2-carbamoylindolin-1-yl)-4-oxobutanoic acid*

$^1$H NMR (500 MHz, DMSO-$d_6$) δ 12.11 (s, 1H), 8.04 (d, $J$ = 8.0 Hz, 1H), 7.72 (s, 1H), 7.28 (s, 1H), 7.20 (d, $J$ = 7.5 Hz, 1H), 7.15 (t, $J$ = 7.5 Hz, 1H), 6.98 (t, $J$ = 7.5 Hz, 1H), 5.00 (dd, $J$ = 11.5, 3.0 Hz, 1H), 3.58 (dd, $J$ = 16.5, 11.0 Hz, 1H), 3.04 (dd, $J$ = 16.5, 3.0 Hz, 1H), 2.74-2.67 (m, 1H), 2.58-2.53 (m, 2H), 2.37-2.29 (m, 1H). $^{13}$C NMR (126 MHz, DMSO-$d_6$) δ 173.80, 172.88, 170.21, 143.35, 129.80, 127.00, 124.41, 123.16, 116.02, 60.41, 34.17, 29.65, 28.66. HRMS (ESI$^+$) m/z calcd for $C_{13}H_{14}N_2NaO_4^+$ [M+Na]$^+$: 285.0851, found 285.0841.

**Synthesis of JZ32-JZ34**

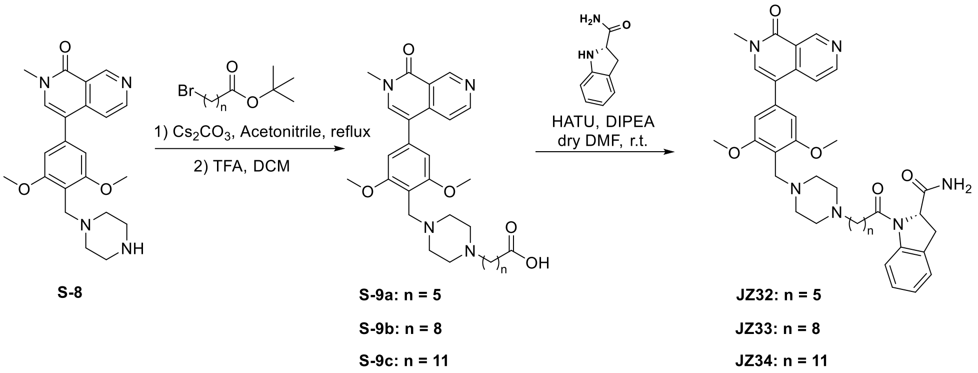


**JZ32:** *(S)-1-(6-(4-(2,6-dimethoxy-4-(2-methyl-1-oxo-1,2-dihydro-2,7-naphthyridin-4-yl)benzyl)piperazin-1-yl)hexanoyl)indoline-2-carboxamide*

Step 1: S-8 was synthesized according to the previously reported method[4]. S-8 (57.0 mg, 0.14 mmol, 1.0 eq), $Cs_2CO_3$ (118 mg, 0.36 mmol, 2.5 eq), and tert-butyl 6-bromohexanoate (54.4 mg, 0.22 mmol, 1.5 eq) were dissolved in acetonitrile (10 mL) and refluxed overnight. Upon completion, the reaction was concentrated, extracted with DCM (30 mL) and washed with brine (1 x 20 mL). The organic layer was dried by $Na_2SO_4$, concentrated, and the resulting residue was used without further purification. The crude product was dissolved in DCM (5.0 mL), TFA (1.0 mL) was added to the reaction and stirred for 1 h. The reaction mixture was then concentrated and purified by flash chromatography to provide S-9a as light yellow oil (46.2 mg, 0.09 mmol, 64.9 % over 2 steps).

$^1$H NMR (500 MHz, MeOD) δ 9.52 (s, 1H), 8.67 (d, *J* = 5.6 Hz, 1H), 7.77 (d, *J* = 1.5 Hz, 1H), 7.64 (d, *J* = 5.7 Hz, 1H), 6.84 (s, 2H), 4.28 (s, 2H), 3.93 (s, 6H), 3.70 (d, *J* = 1.3 Hz, 3H), 3.37 – 3.33 (m, 4H), 3.19 (s, 4H), 2.92 – 2.85 (m, 2H), 2.32 (t, *J* = 7.3 Hz, 2H), 1.73 – 1.61 (m, 4H), 1.42 (qd, *J* = 9.0, 6.3 Hz, 2H).

Step 2: S-9a (46.2 mg, 0.09 mmol, 1 eq), HATU (44.9 mg, 0.12 mmol, 1.3 eq), and (2S)-Indoline-2-carboxamide (17.7 mg, 0.11 mmol, 1.2 eq) were dissolved in dry DMF (5.0 mL) at room temperature. DIPEA (78.4 µL, 0.45 mmol, 5.0 eq) was added to the reaction and stirred for 2 hours. The reaction was then concentrated and extracted with DCM (30 mL) and brine (30 mL). The organic layer was dried with anhydrous sodium sulfate. The reaction mixture was then concentrated and purified by flash chromatography to provide JZ32 as a white solid (35.8 mg, 0.05 mmol, 60.4% yield).

$^1$H NMR (500 MHz, MeOD) δ 7.96 (s, 1H), 7.12 (d, *J* = 5.8 Hz, 1H), 6.60 (d, *J* = 8.1 Hz, 1H), 6.18 (s, 1H), 6.06 (d, *J* = 5.7 Hz, 1H), 5.68 – 5.59 (m, 2H), 5.48 (t, *J* = 7.5 Hz, 1H), 5.24 (s, 2H), 3.52 – 3.46 (m, 1H), 2.49 (s, 2H), 2.36 (s, 6H), 2.14 (s, 3H), 1.68 – 1.17 (m, 10H), 1.09 (t, *J* = 7.8 Hz, 2H), 1.01 – 0.91 (m, 1H), 0.83 – 0.71 (m, 1H), 0.23 – 0.16 (m, 2H), 0.12 – 0.04 (m, 2H), -0.07 – -0.15 (m, 2H). $^{13}$C NMR (126 MHz, MeOD) δ 176.46, 173.96, 163.04, 160.97, 151.68, 151.21, 144.27, 143.52, 138.96, 138.61, 131.00, 128.41, 125.52, 125.20, 121.68, 119.24, 119.11, 118.27, 106.63, 62.59, 58.52, 56.55, 52.57, 52.41, 49.77, 37.43, 36.95, 35.81, 35.65, 27.65, 26.45, 25.23. HRMS (ESI$^+$) m/z calcd for $C_{37}H_{45}N_6O_5^+$ [M+H]$^+$: 653.3451, found 653.3430.

The synthetic route of JZ33 and JZ34 were similar to JZ32.

**JZ33:** *(S)-1-(9-(4-(2,6-dimethoxy-4-(2-methyl-1-oxo-1,2-dihydro-2,7-naphthyridin-4-yl)benzyl)piperazin-1-yl)nonanoyl)indoline-2-carboxamide*

$^1$H NMR (500 MHz, MeOD) δ 9.51 (s, 1H), 8.70 – 8.65 (m, 1H), 8.15 (d, *J* = 8.1 Hz, 1H), 7.73 (s, 1H), 7.63 (d, *J* = 5.7 Hz, 1H), 7.23 – 7.15 (m, 2H), 7.03 (t, *J* = 7.4 Hz, 1H), 6.81 (s, 2H), 5.07 – 5.01 (m, 1H), 4.15 (s, 2H), 3.92 (s, 6H), 3.70 (s, 3H), 3.17 – 2.96 (m, 8H), 2.81 – 2.74 (m, 2H), 2.55 – 2.42 (m, 1H), 2.37 – 2.26 (m, 1H), 1.74 – 1.68 (m, 2H), 1.66 – 1.58 (m, 2H), 1.46 – 1.29 (m, 10H). $^{13}$C NMR (126 MHz, MeOD) δ 176.48, 174.20, 163.01, 160.94, 151.62, 151.17, 144.30, 143.45, 139.13, 139.06, 131.01, 129.32, 128.40, 125.51, 125.15, 121.67, 119.25, 118.95, 118.24, 109.40, 106.68, 62.58, 58.29, 56.64, 52.06, 51.81, 49.76, 37.45, 36.05, 35.64, 30.10, 30.03, 27.76, 26.23, 25.59. HRMS (ESI$^+$) m/z calcd for $C_{40}H_{51}N_6O_5^+$ [M+H]$^+$: 695.3921, found 695.3897.

**JZ34:** *(S)-1-(12-(4-(2,6-dimethoxy-4-(2-methyl-1-oxo-1,2-dihydro-2,7-naphthyridin-4-yl)benzyl)piperazin-1-yl)dodecanoyl)indoline-2-carboxamide*

$^1$H NMR (500 MHz, MeOD) δ 9.51 (s, 1H), 8.68 (d, *J* = 5.7 Hz, 1H), 8.15 (d, *J* = 8.1 Hz, 1H), 7.73 (s, 1H), 7.63 (d, *J* = 5.7 Hz, 1H), 7.22 – 7.16 (m, 2H), 7.07 – 7.00 (m, 1H), 6.82 (s, 2H), 5.06 – 5.01 (m, 1H), 4.14 (s, 2H), 3.93 (s, 6H), 3.70 (s, 3H), 3.26 – 2.64 (m, 10H), 2.54 – 2.43 (m, 1H), 2.36 – 2.26 (m, 1H), 1.77 – 1.67 (m, 2H), 1.65 – 1.58 (m, 2H), 1.45 – 1.31 (m, 16H). $^{13}$C NMR (126 MHz, MeOD) δ 176.48, 174.23, 163.02, 160.95, 151.66, 151.23, 144.29, 143.45, 139.02, 130.99, 128.40, 125.50, 125.14, 121.66, 119.22, 118.98, 118.26, 106.69, 62.60, 58.40, 56.64, 52.19, 49.77, 37.44, 36.19, 35.64, 30.49, 30.45, 30.27, 30.25, 27.93, 26.39, 25.75. HRMS (ESI$^+$) m/z calcd for $C_{43}H_{57}N_6O_5^+$ [M+H]$^+$: 737.4390, found 737.4361.

### Synthesis of JZ35-JZ37

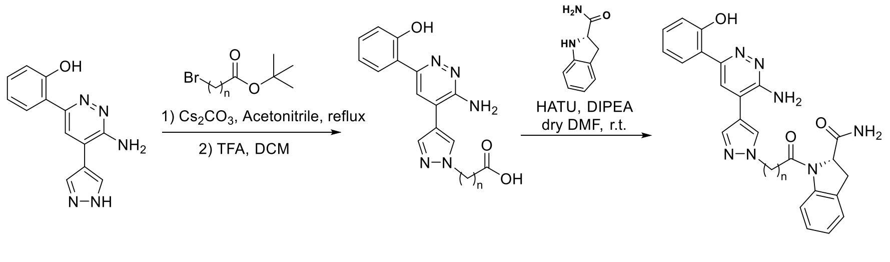


**S-10**

**S-11a: n = 5**
**S-11b: n = 8**
**S-11c: n = 11**

**JZ35: n = 5**
**JZ36: n = 8**
**JZ37: n = 11**

**JZ35:** *(S)-1-(6-(4-(3-amino-6-(2-hydroxyphenyl)pyridazin-4-yl)-1H-pyrazol-1-yl)hexanoyl)indoline-2-carboxamide*

S-10 was synthesized according to the previously reported method[5]. S-10 (42.0 mg, 0.17 mmol, 1.0 eq), $Cs_2CO_3$ (297 mg, 0.91 mmol, 5.5 eq), and tert-butyl 6-bromohexanoate (83.3 mg, 0.33 mmol, 2.0 eq) were dissolved in acetonitrile (10 mL) and refluxed overnight. Upon completion, the reaction was concentrated, extracted with DCM (30 mL) and washed with brine (1 x 20 mL). The organic layer was dried by $Na_2SO_4$, concentrated, and the resulting residue was used without further purification. The crude product was dissolved in DCM (5.0 mL), TFA (1.0 mL) was added to the reaction and stirred for 1 h. The reaction mixture was then concentrated and used without further purification to provide S-11a. S-11a (23.2 mg, 0.06 mmol, 1 eq), HATU (31.2 mg, 0.08 mmol, 1.3 eq), and (2S)-Indoline-2-carboxamide (12.3 mg, 0.108 mmol, 1.2 eq) were dissolved in dry DMF (5.0 mL) at room temperature. DIPEA (54.5 µL, 0.32 mmol, 5.0 eq) was added to the reaction and stirred for 2 hours. The reaction was then concentrated and extracted with DCM (30 mL) and brine (30 mL). The organic layer was dried with anhydrous $Na_2SO_4$. The reaction mixture was then concentrated and purified by flash chromatography to provide JZ35 as a light yellow oil (21.5 mg, 0.04 mmol, 23.5% yield over 3 steps).

$^1$H NMR (500 MHz, $CDCl_3$) δ 7.92 (s, 1H), 7.83 (s, 1H), 7.77 (s, 1H), 7.61 (d, *J* = 7.9 Hz, 1H), 7.23 (d, *J* = 7.4 Hz, 1H), 7.16 – 7.12 (m, 2H), 7.02 – 6.97 (m, 2H), 6.87 (t, *J* = 7.5 Hz, 1H), 4.78 (s, 1H), 4.19 (t, *J* = 6.8 Hz, 3H), 2.32 – 2.22 (m, 1H), 1.97 – 1.87 (m, 2H), 1.79 – 1.66 (m, 3H), 1.24 – 1.11 (m, 4H). $^{13}$C NMR (126 MHz, $CDCl_3$) δ 174.31, 172.00, 158.49, 156.31, 154.02, 138.38, 130.97, 129.27, 125.42, 124.75, 122.54, 122.30, 119.18, 118.26, 117.38, 115.05, 68.22, 62.02, 53.47, 52.30, 51.64, 33.66, 29.77, 29.70, 25.88, 24.14, 23.67, 14.07. HRMS (ESI$^+$) m/z calcd for $C_{28}H_{30}N_7O_3^+$ $[M+H]^+$: 512.2410, found 512.2375.

The synthetic route of JZ36 and JZ37 were similar to JZ35.

**JZ36:** *(S)-1-(9-(4-(3-amino-6-(2-hydroxyphenyl)pyridazin-4-yl)-1H-pyrazol-1-yl)nonanoyl)indoline-2-carboxamide*

$^{1}$H NMR (500 MHz, $CDCl_3$) δ 7.86 – 7.82 (m, 2H), 7.76 (s, 1H), 7.62 – 7.57 (m, 1H), 7.29 – 7.20 (m, 2H), 7.18 – 7.11 (m, 2H), 7.04 – 6.95 (m, 2H), 6.91 – 6.82 (m, 1H), 4.78 (s, 1H), 4.19 – 4.12 (m, 2H), 3.70 – 3.45 (m, 1H), 3.38 – 3.26 (m, 1H), 2.65 (s, 1H), 2.43 – 2.18 (m, 1H), 1.94 – 1.82 (m, 2H), 1.65 (s, 2H), 1.30 (s, 8H). $^{13}$C NMR (126 MHz, $CDCl_3$) δ 158.49, 156.29, 154.06, 138.21, 130.99, 128.84, 127.92, 125.39, 124.66, 122.63, 122.32, 119.18, 118.28, 117.35, 115.07, 77.42, 77.16, 76.91, 62.13, 52.78, 35.36, 34.54, 30.16, 29.70, 29.12, 29.00, 28.80, 26.46, 24.47. HRMS (ESI$^+$) m/z calcd for $C_{31}H_{36}N_7O_3^+$ $[M+H]^+$: 554.2880, found 554.2844.

**JZ37:** *(S)-1-(12-(4-(3-amino-6-(2-hydroxyphenyl)pyridazin-4-yl)-1H-pyrazol-1-yl)dodecanoyl)indoline-2-carboxamide*

$^{1}$H NMR (500 MHz, $CDCl_3$) δ 7.82 (d, *J* = 9.0 Hz, 2H), 7.74 (s, 1H), 7.58 (d, *J* = 8.0 Hz, 1H), 7.24 – 7.19 (m, 1H), 7.16 – 7.00 (m, 3H), 6.99 – 6.92 (m, 2H), 6.85 (t, *J* = 7.6 Hz, 1H), 4.77 (s, 1H), 4.13 (s, 2H), 3.34 – 3.21 (m, 3H), 2.63 (s, 1H), 2.44 – 2.01 (m, 2H), 1.91 – 1.78 (m, 2H), 1.71 – 1.54 (m, 2H), 1.31 – 1.18 (m, 12H). $^{13}$C NMR (126 MHz, $CDCl_3$) δ 158.37, 156.29, 153.95, 142.13, 140.34, 138.14, 133.21, 130.96, 128.84, 127.83, 125.38, 124.59, 122.64, 122.36, 119.19, 118.18, 117.32, 115.00, 62.06, 55.90, 52.78, 35.42, 34.53, 31.88, 30.19, 29.64, 29.36, 29.34, 29.32, 29.17, 28.99, 26.52, 24.56, 13.99. HRMS (ESI+) m/z calcd for $C_{34}H_{42}N_7O_3^+$ $[M+H]^+$: 596.3349, found 596.3311.

### $^{1}$H NMR spectrum of JZ1

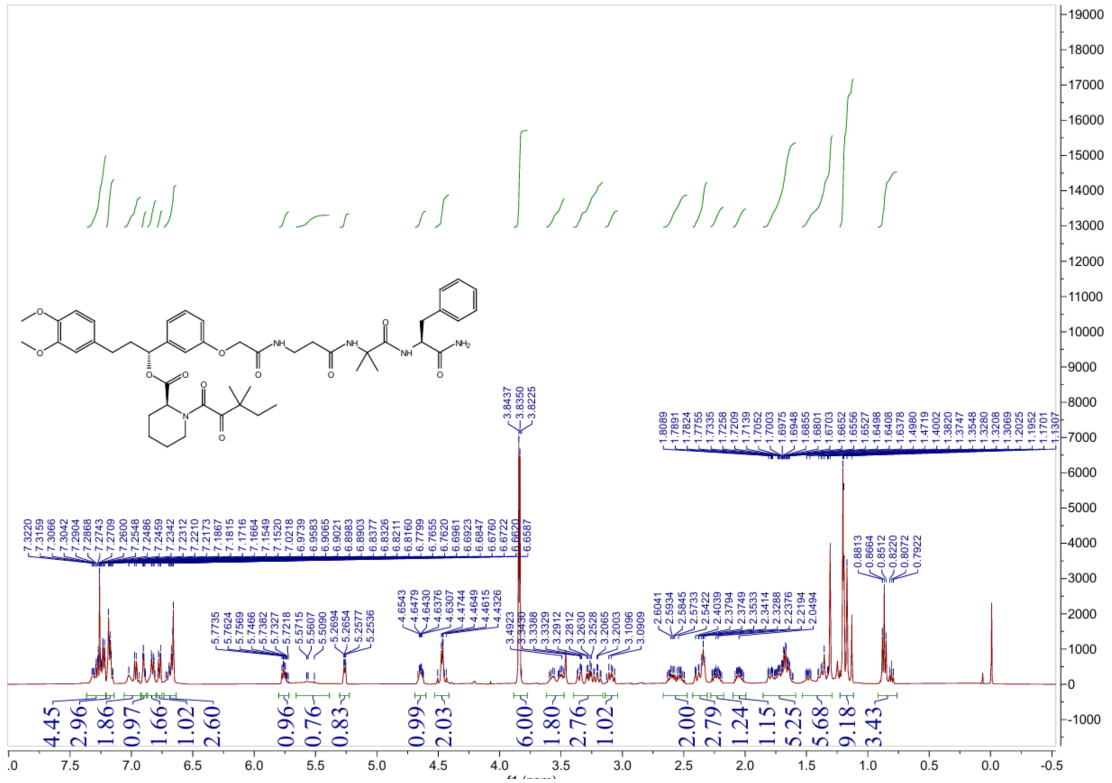

### $^{13}$C NMR spectrum of JZ1

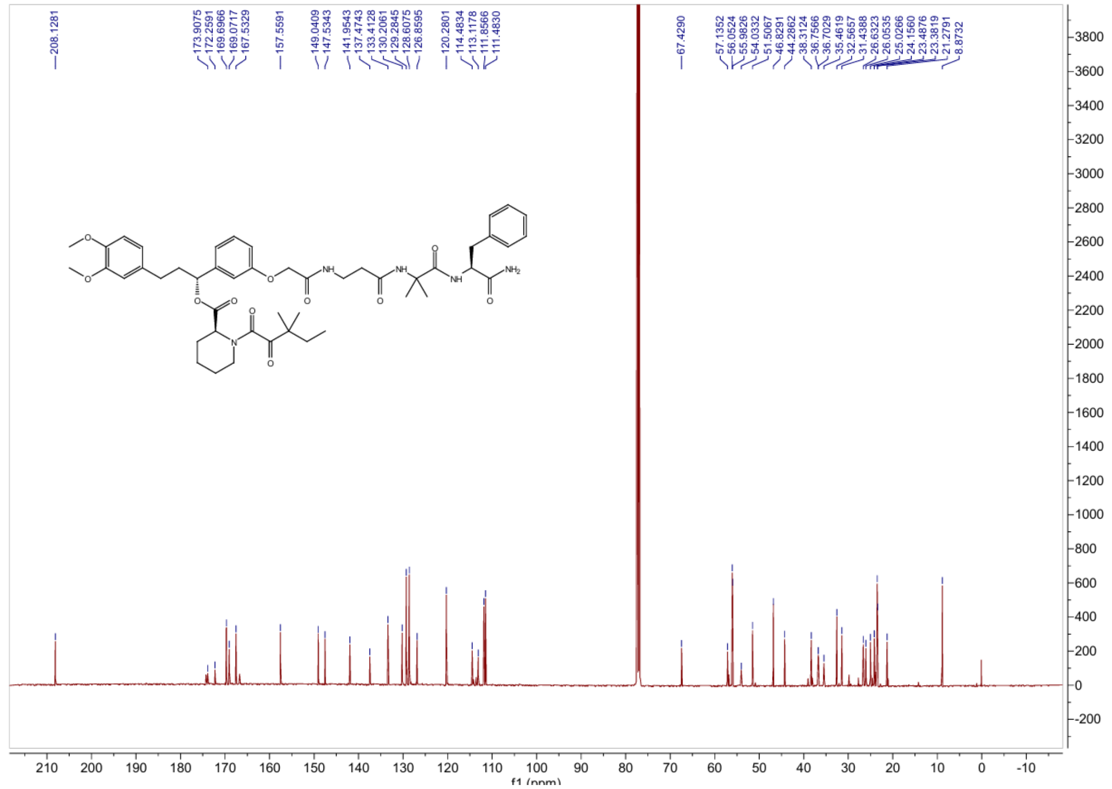

### $^{1}$H NMR spectrum of JZ2

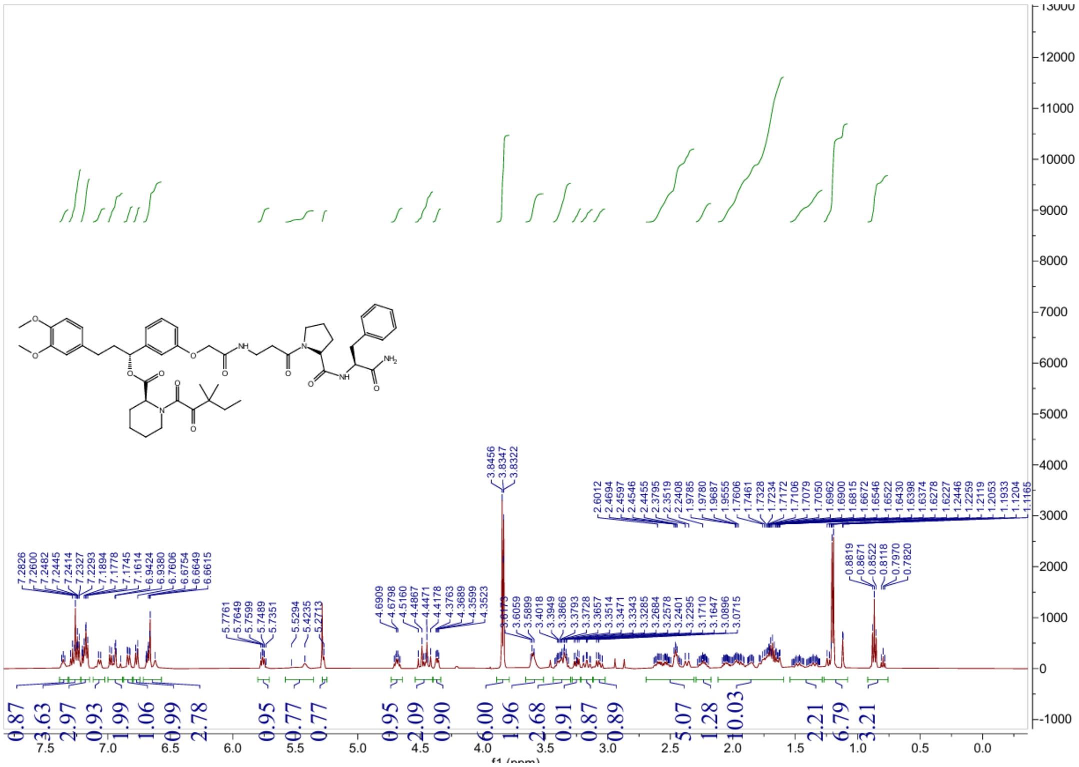


### $^{13}$C NMR spectrum of JZ2

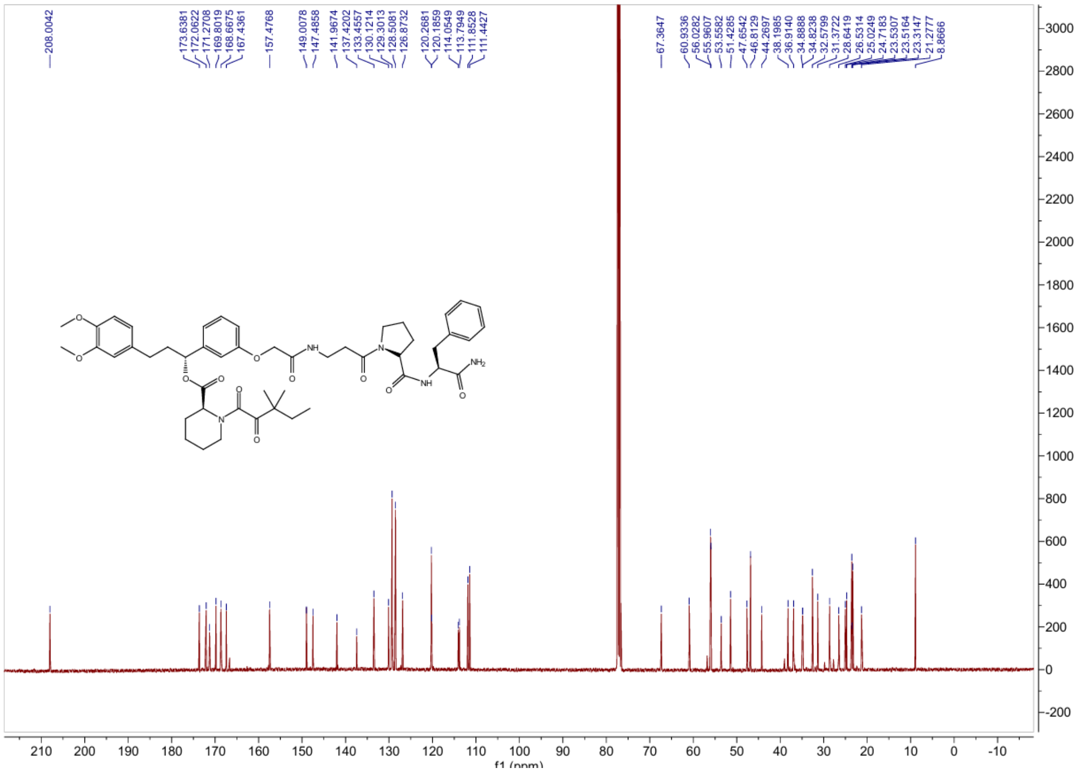

### [1]H NMR spectrum of JZ3

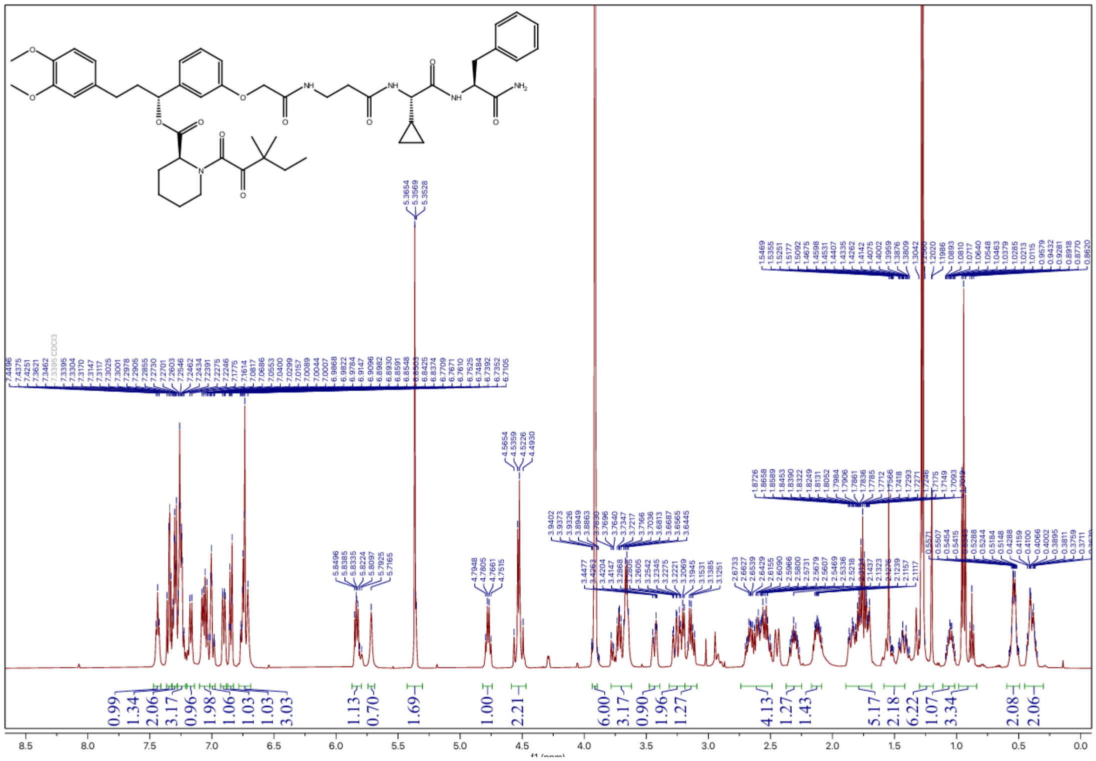

### [13]C NMR spectrum of JZ3

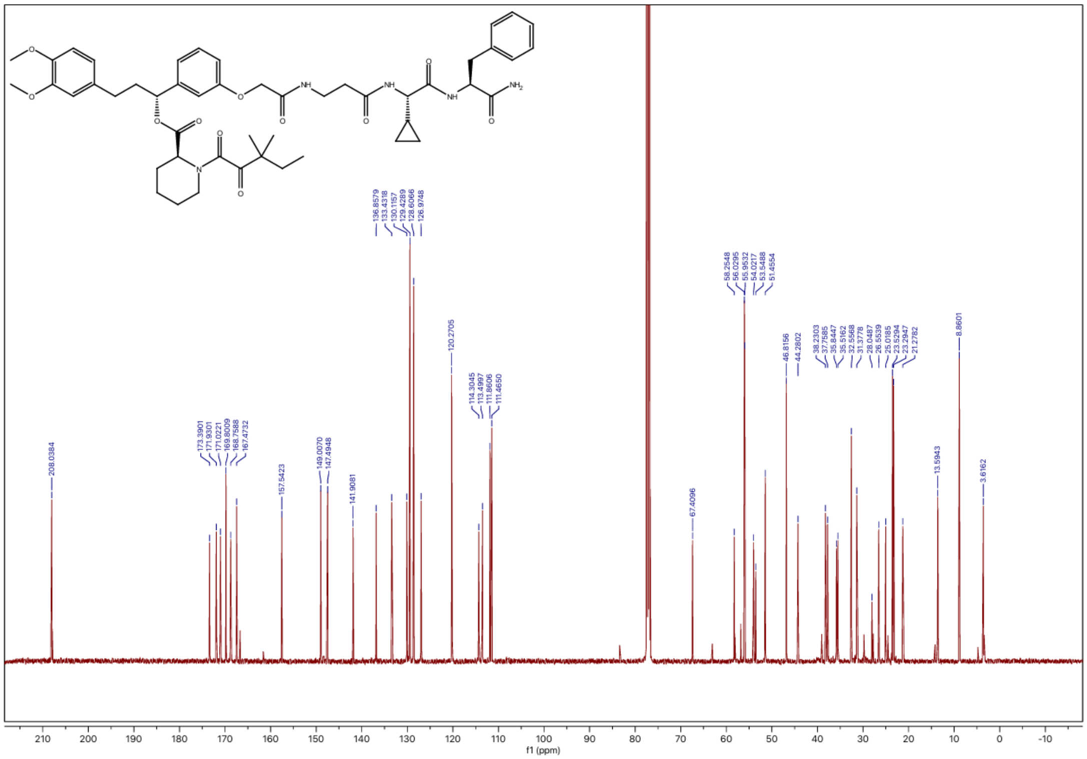

### $^1$H NMR spectrum of JZ4

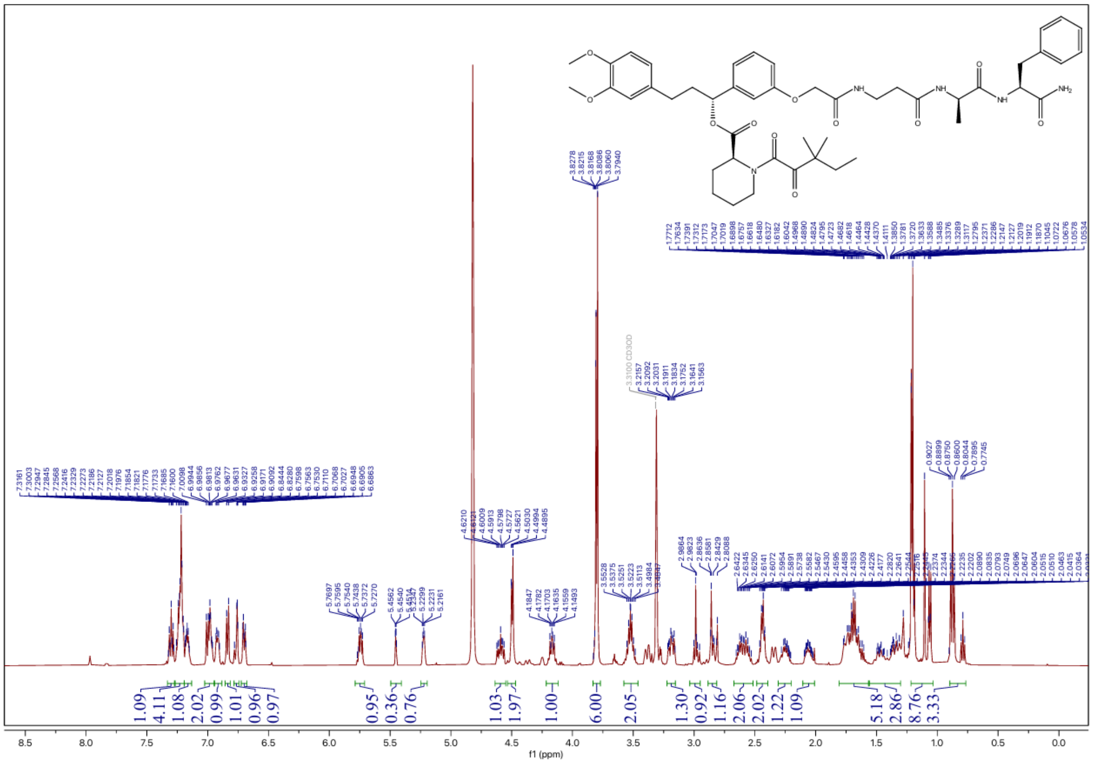


### $^{13}$C NMR spectrum of JZ4

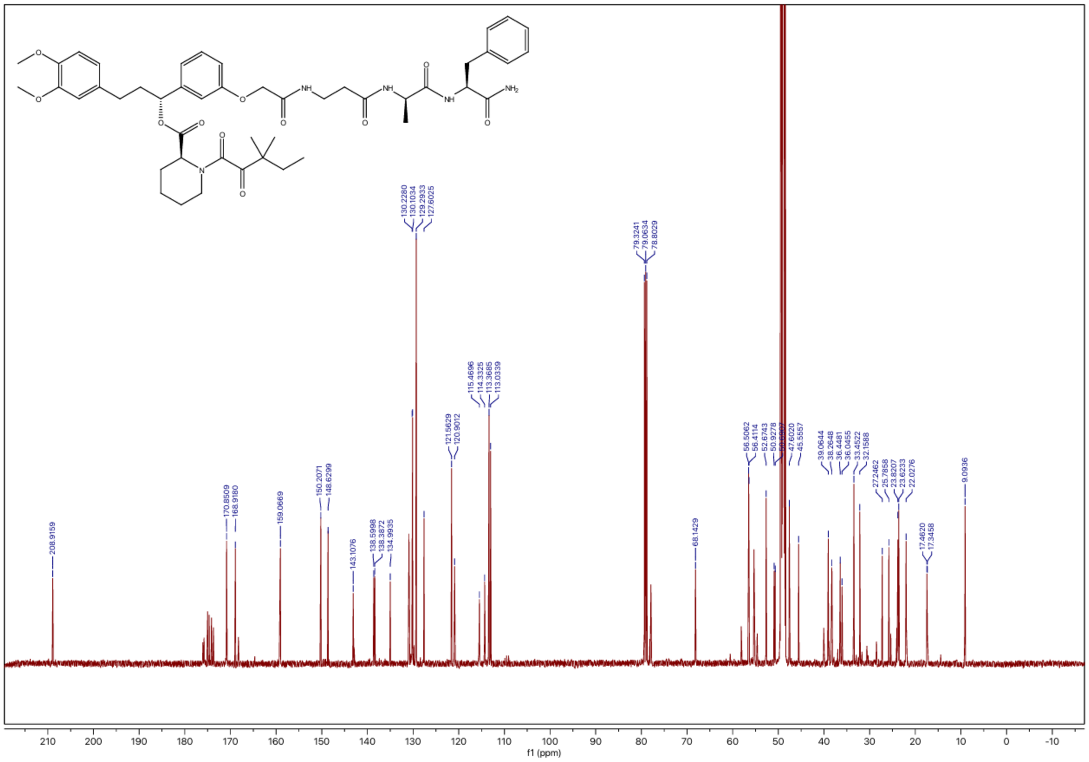

### [1]H NMR spectrum of JZ5

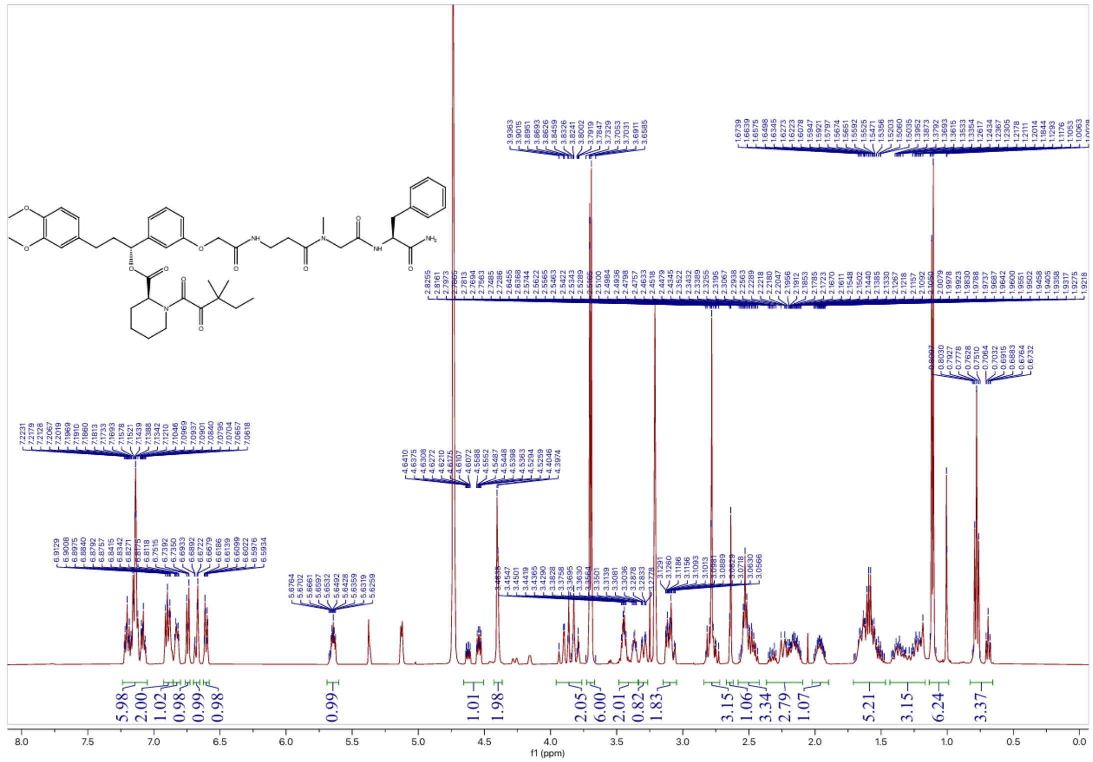

### [13]C NMR spectrum of JZ5

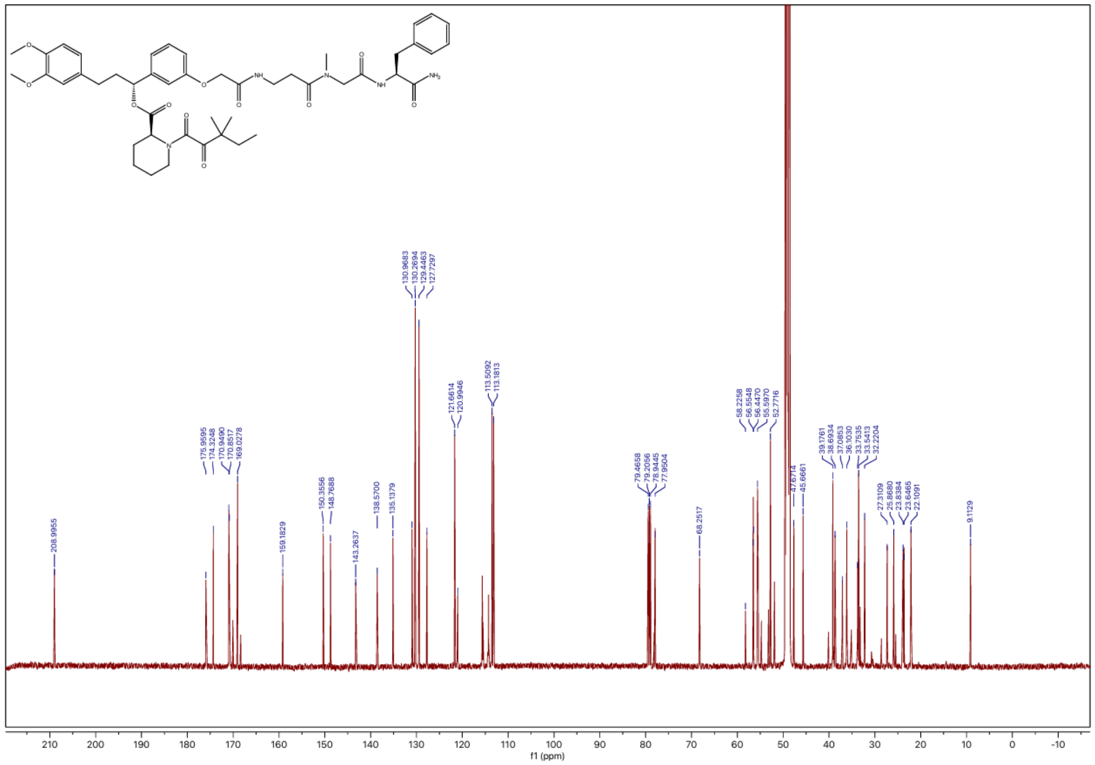

$^{1}H$ NMR spectrum of JZ6

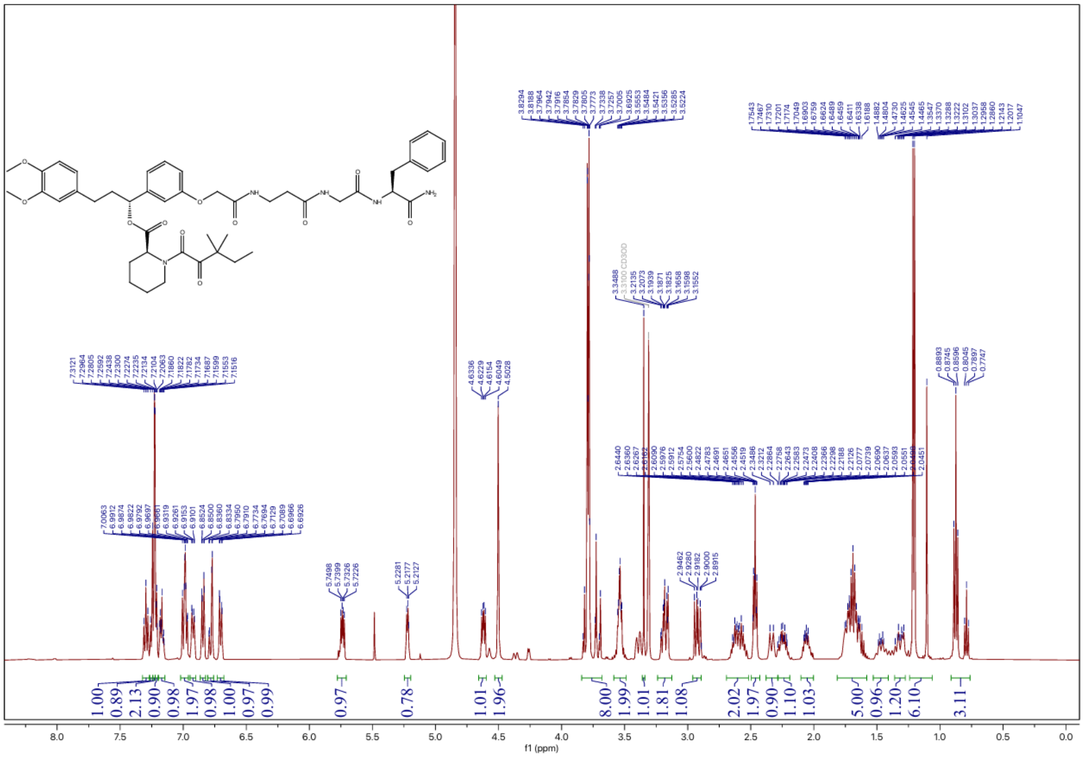

$^{13}C$ NMR spectrum of JZ6

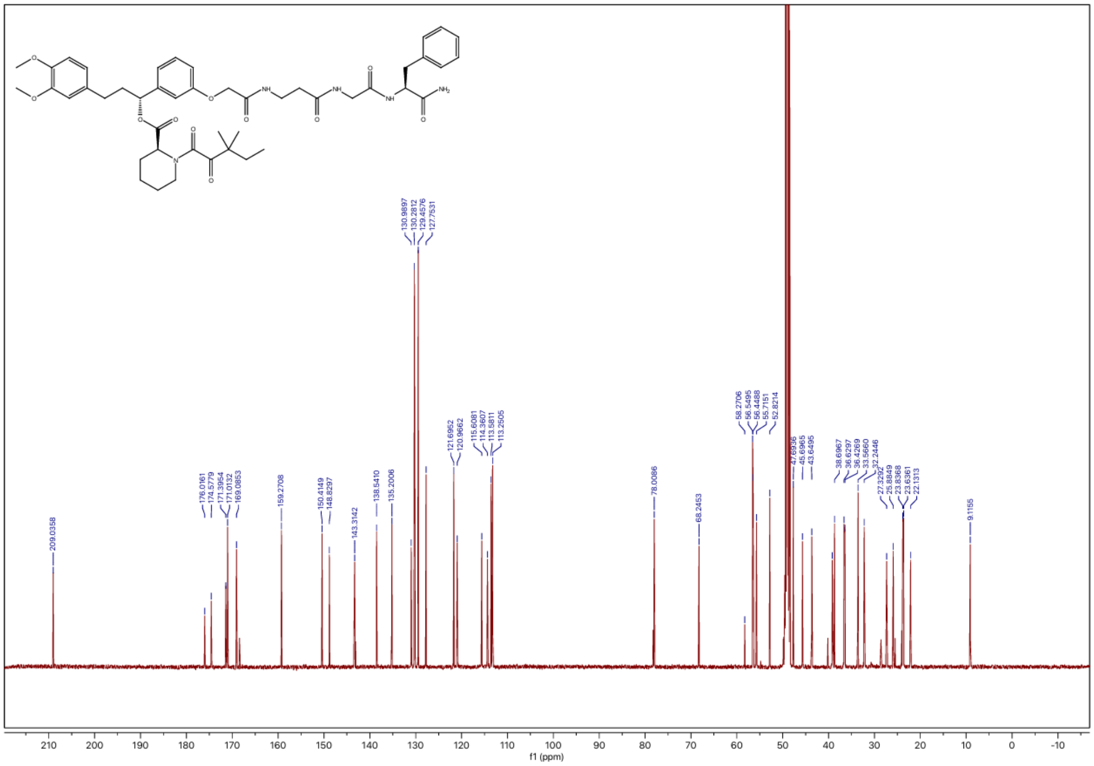

[1]H NMR spectrum of JZ7

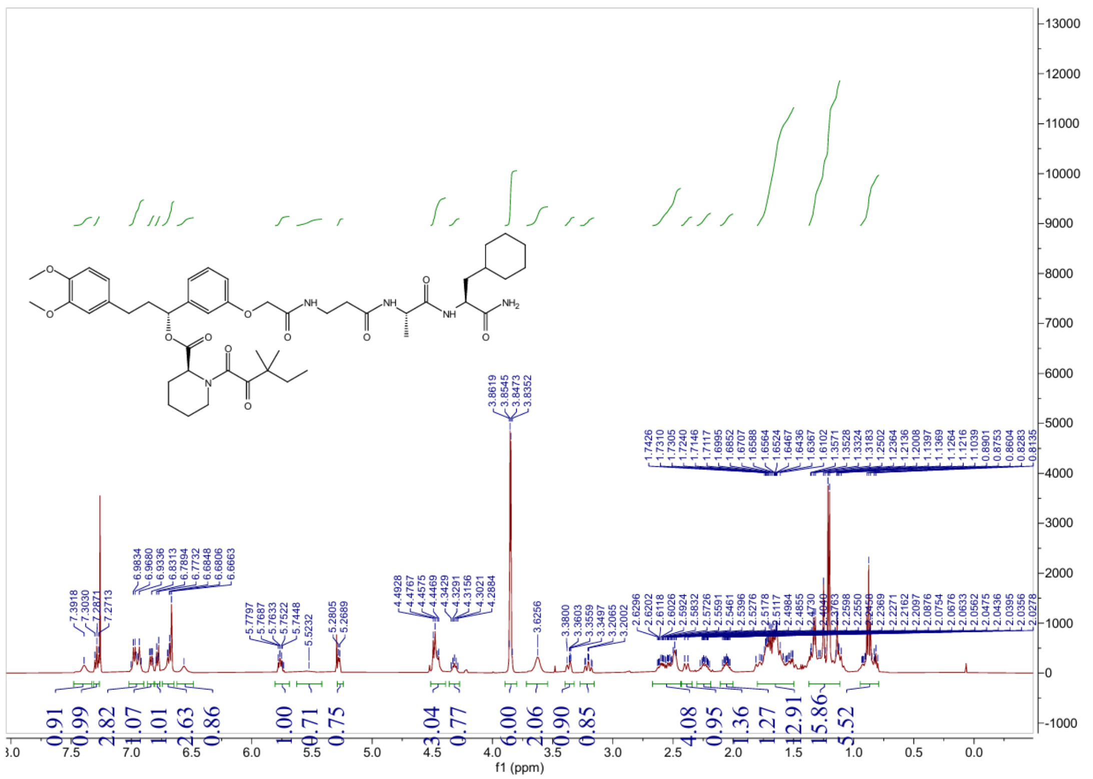

[13]C NMR spectrum of JZ7

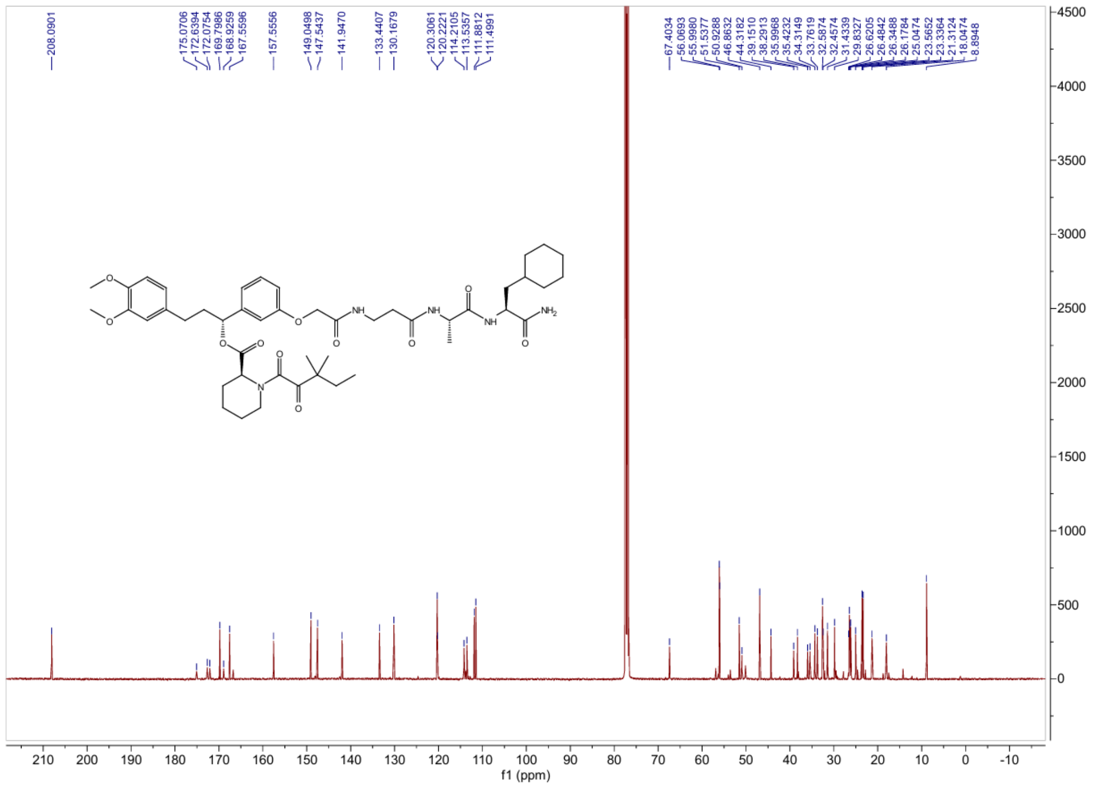

### $^{1}H$ NMR spectrum of JZ8

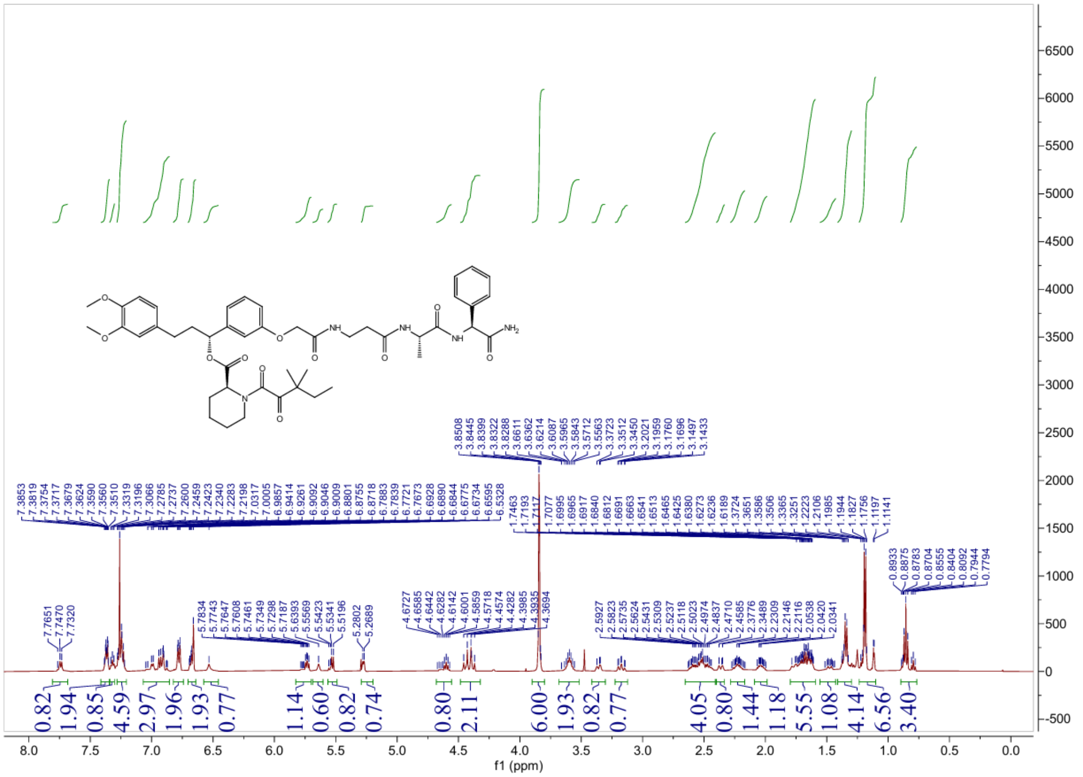

### $^{13}C$ NMR spectrum of JZ8

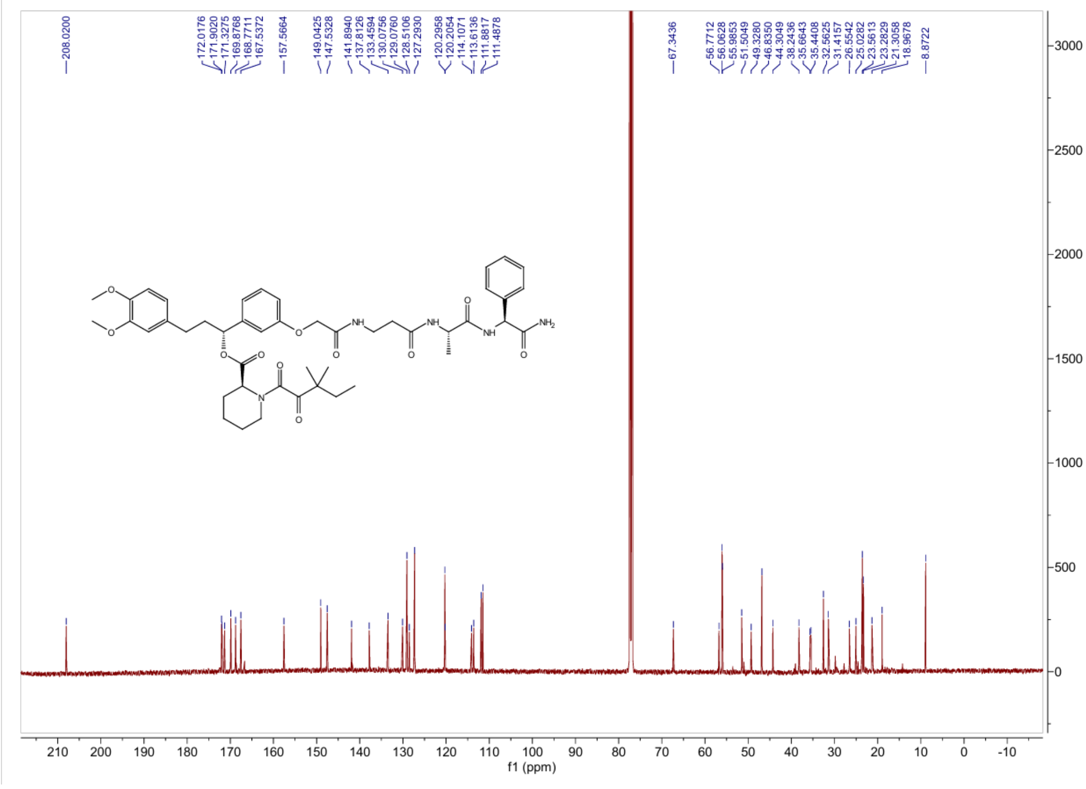

### $^{1}$H NMR spectrum of JZ9

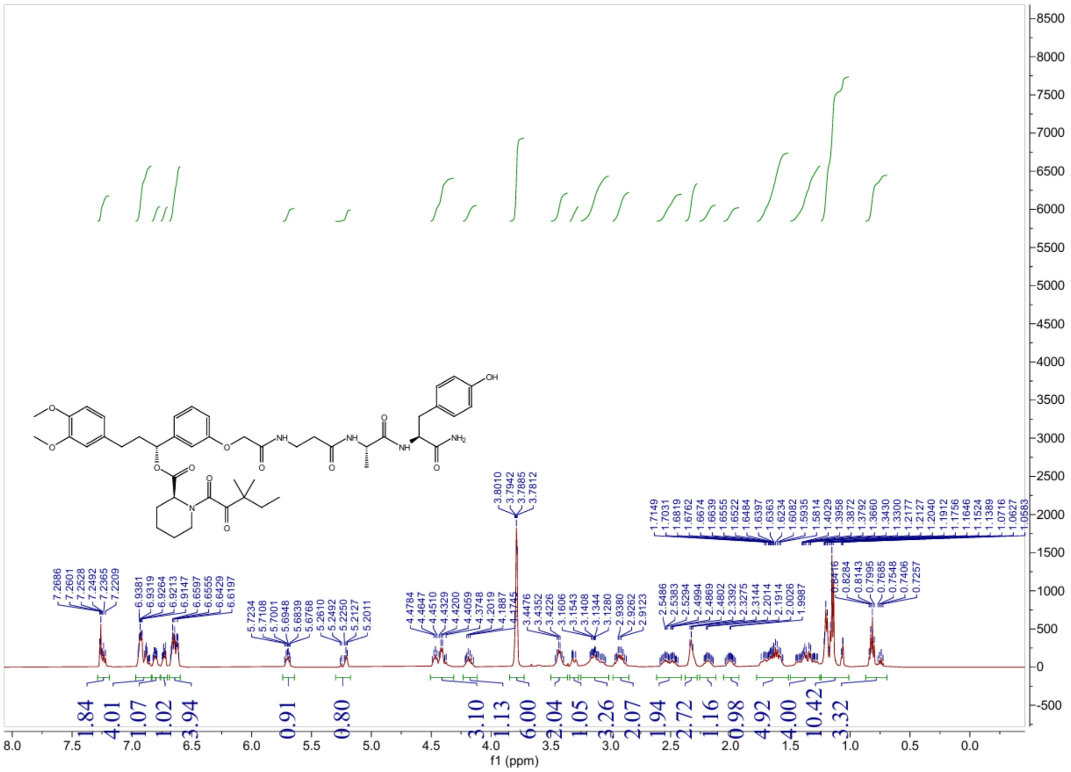

### $^{13}$C NMR spectrum of JZ9

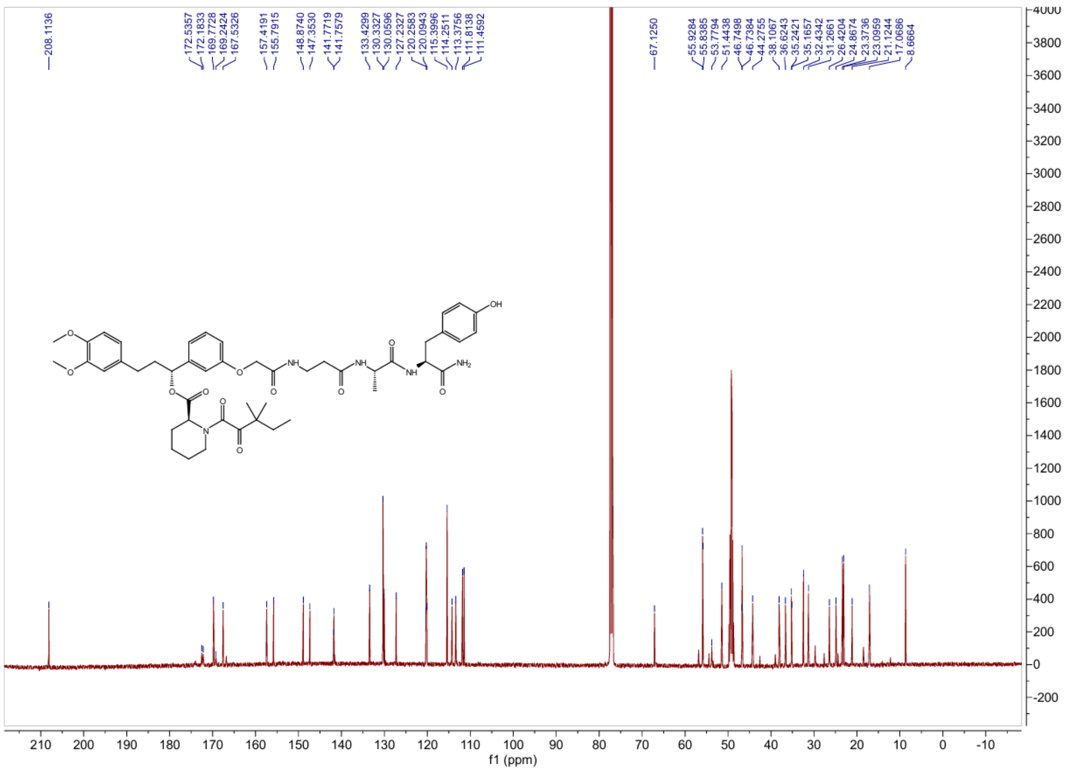

### $^{1}$H NMR spectrum of JZ10

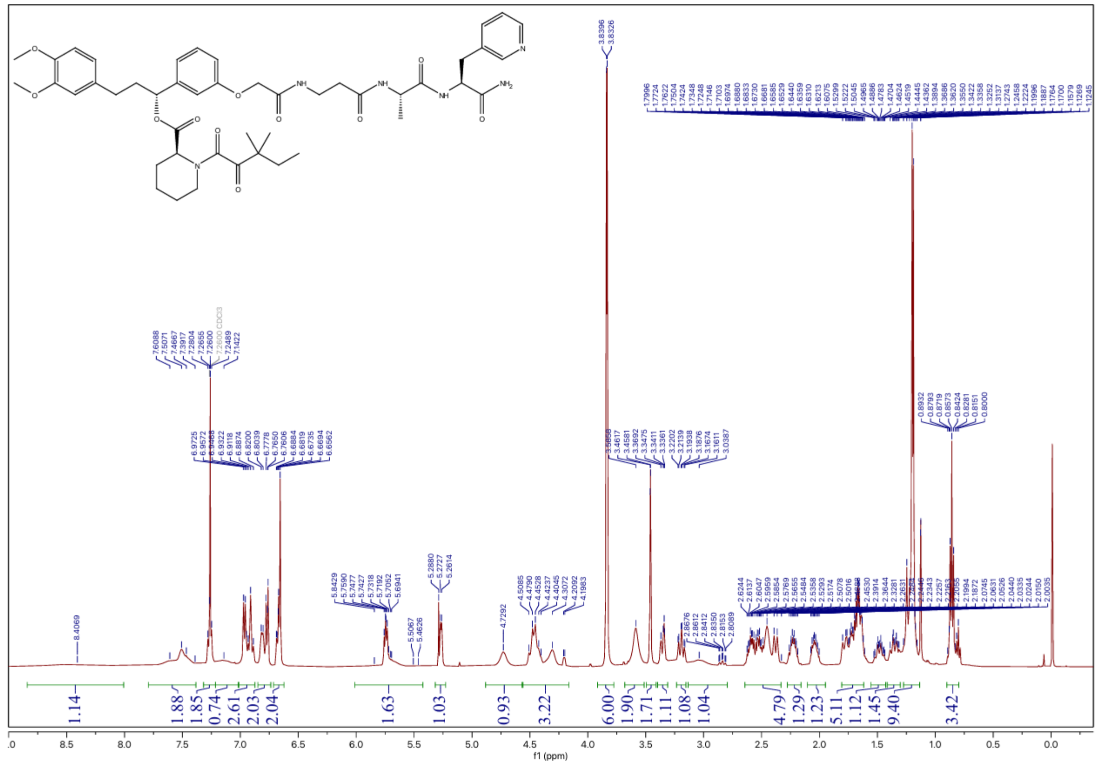

### $^{13}$C NMR spectrum of JZ10

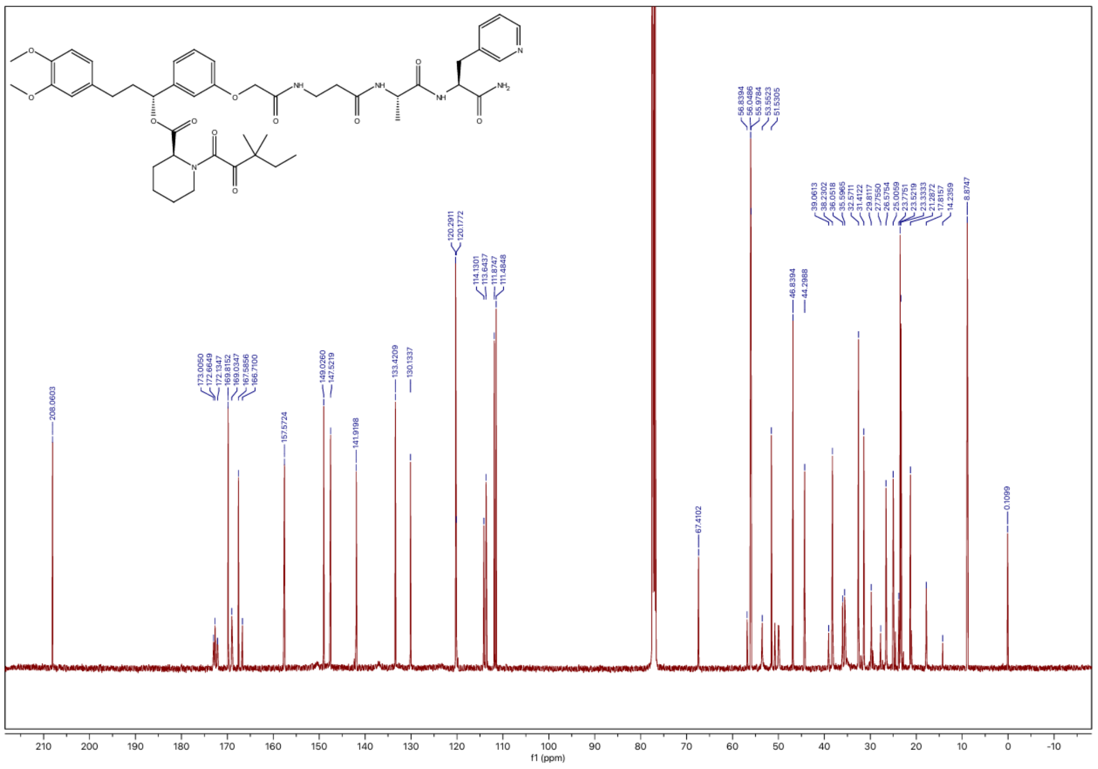

### $^{1}$H NMR spectrum of JZ11

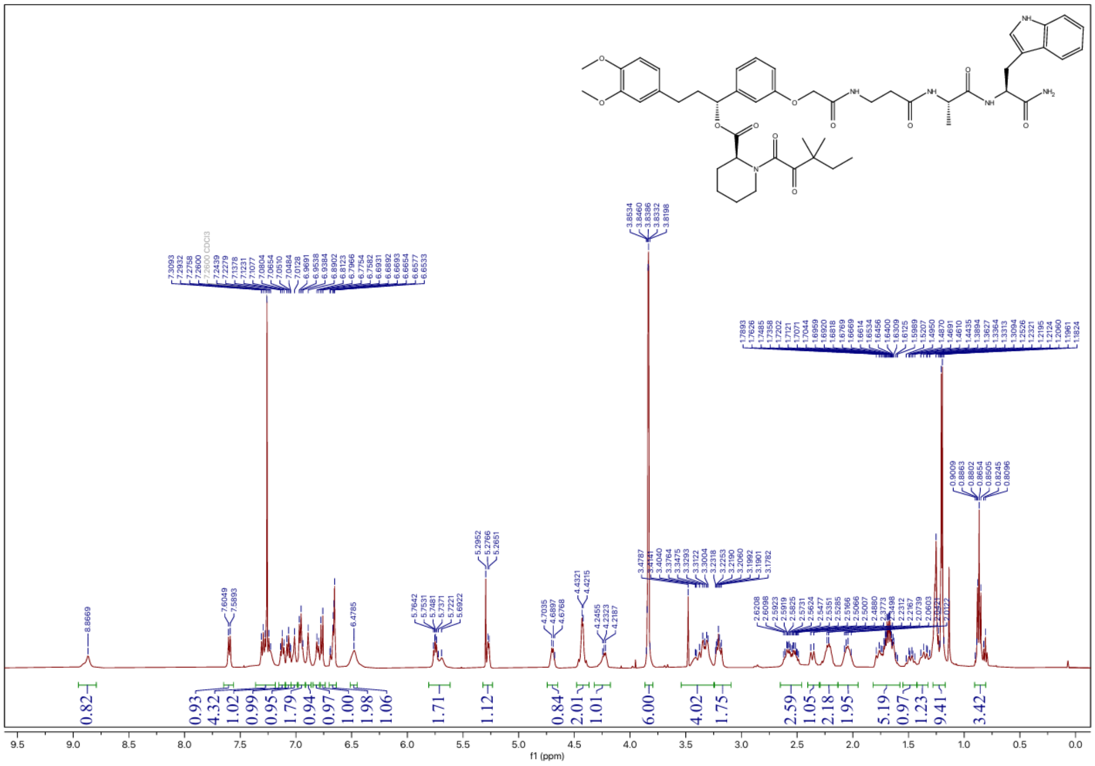

### $^{13}$C NMR spectrum of JZ11

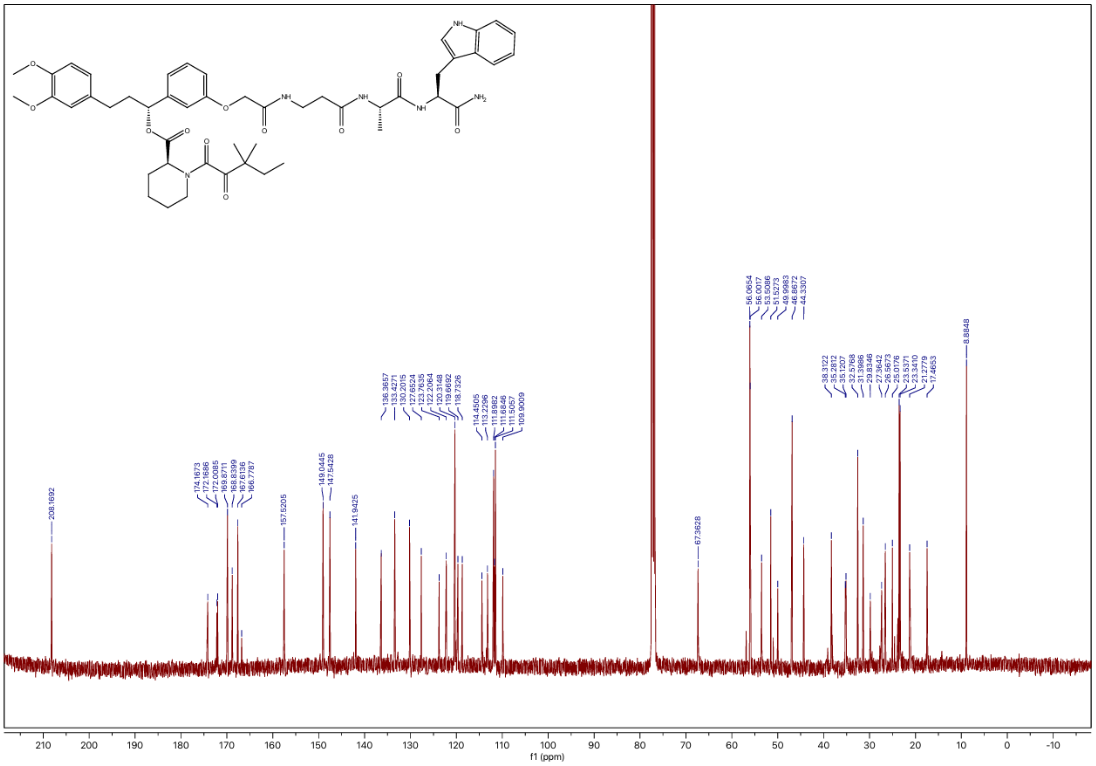

### $^1$H NMR spectrum of JZ12

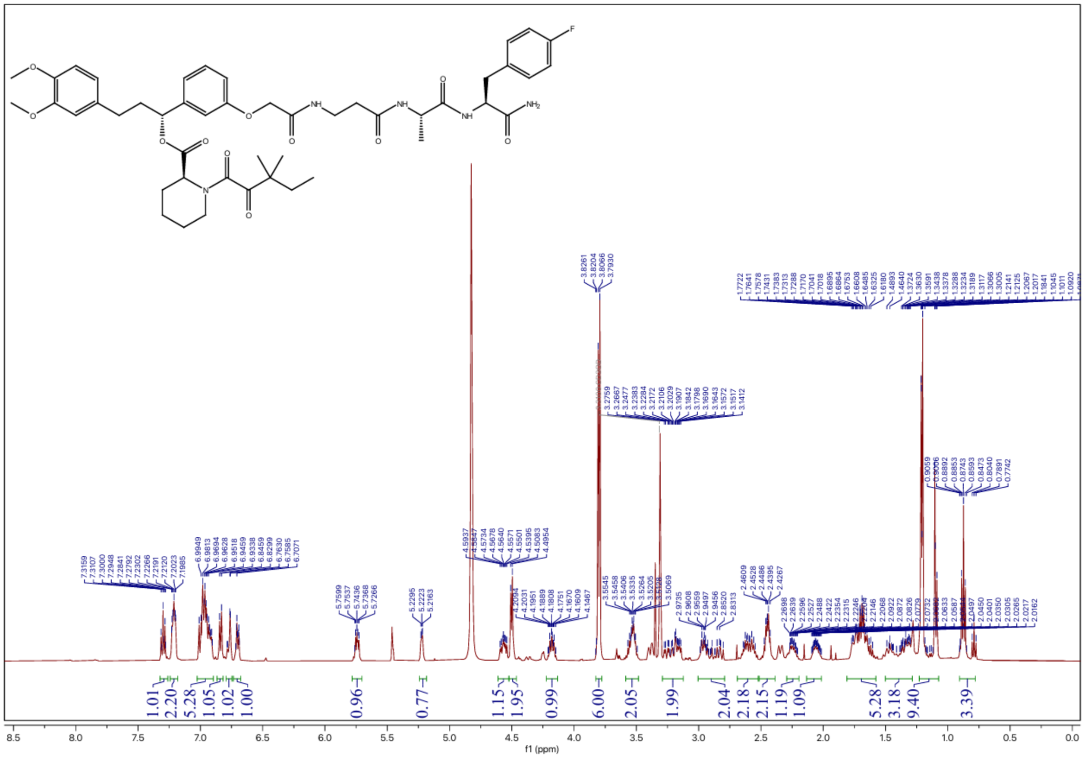

### $^{13}$C NMR spectrum of JZ12

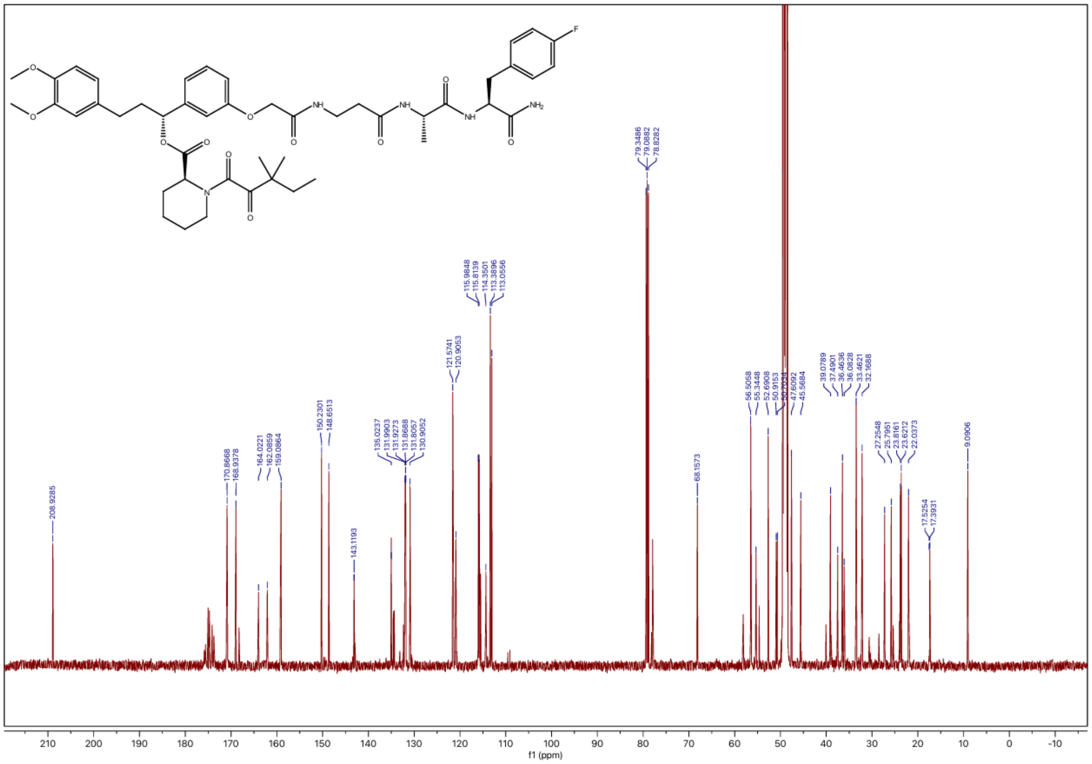

### $^{1}$H NMR spectrum of JZ13

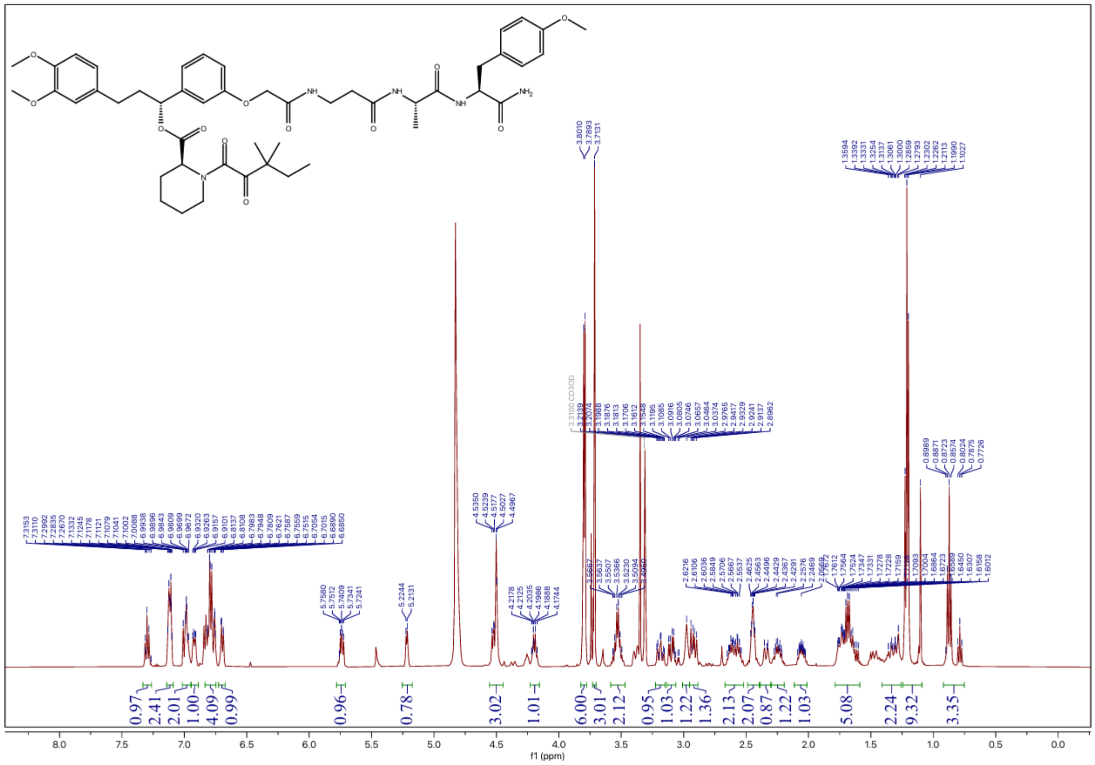

### $^{13}$C NMR spectrum of JZ13

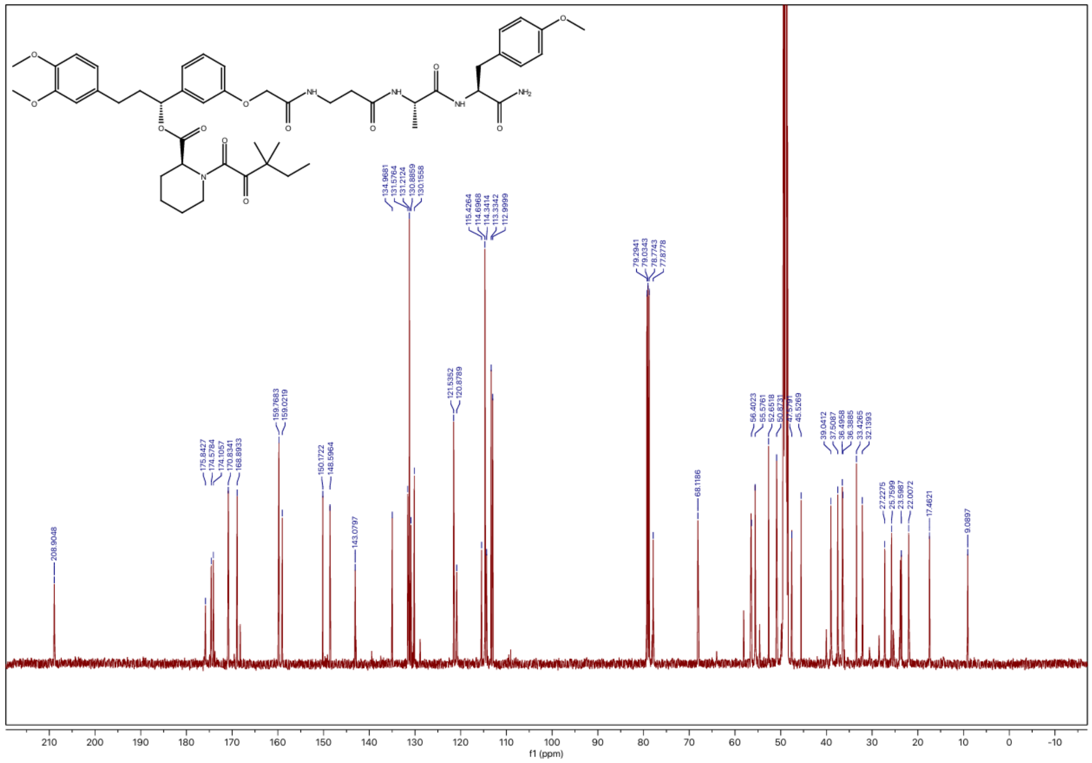

### $^{1}H$ NMR spectrum of JZ14

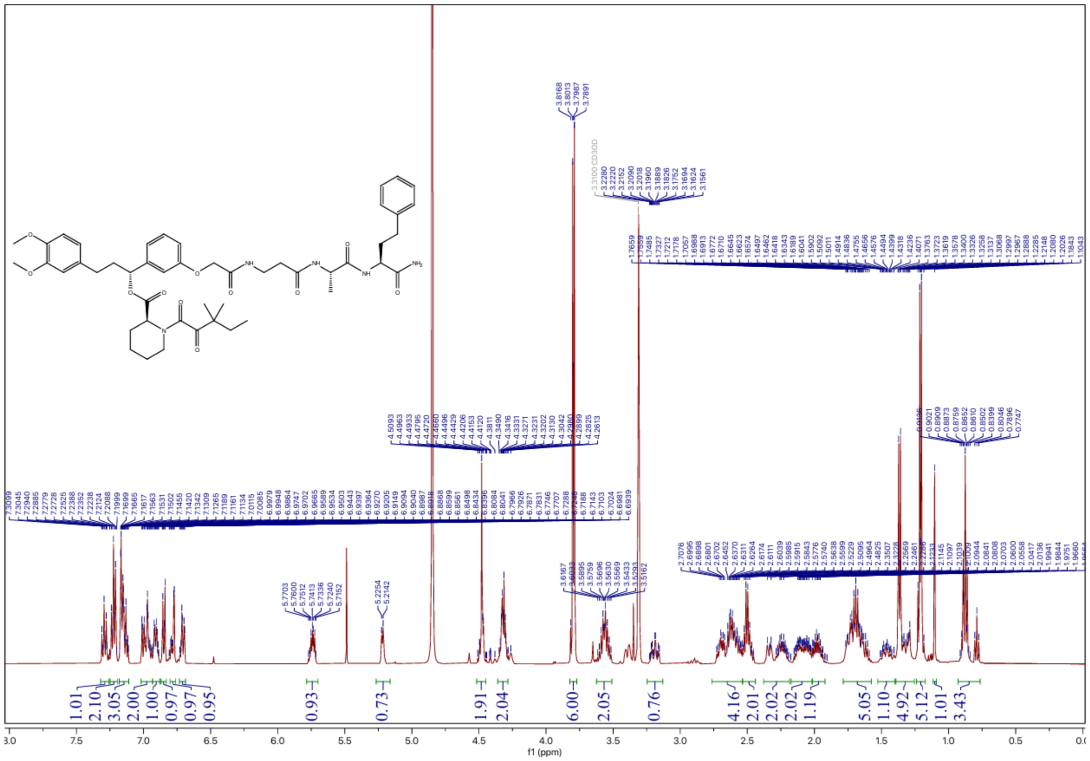

### $^{13}C$ NMR spectrum of JZ14

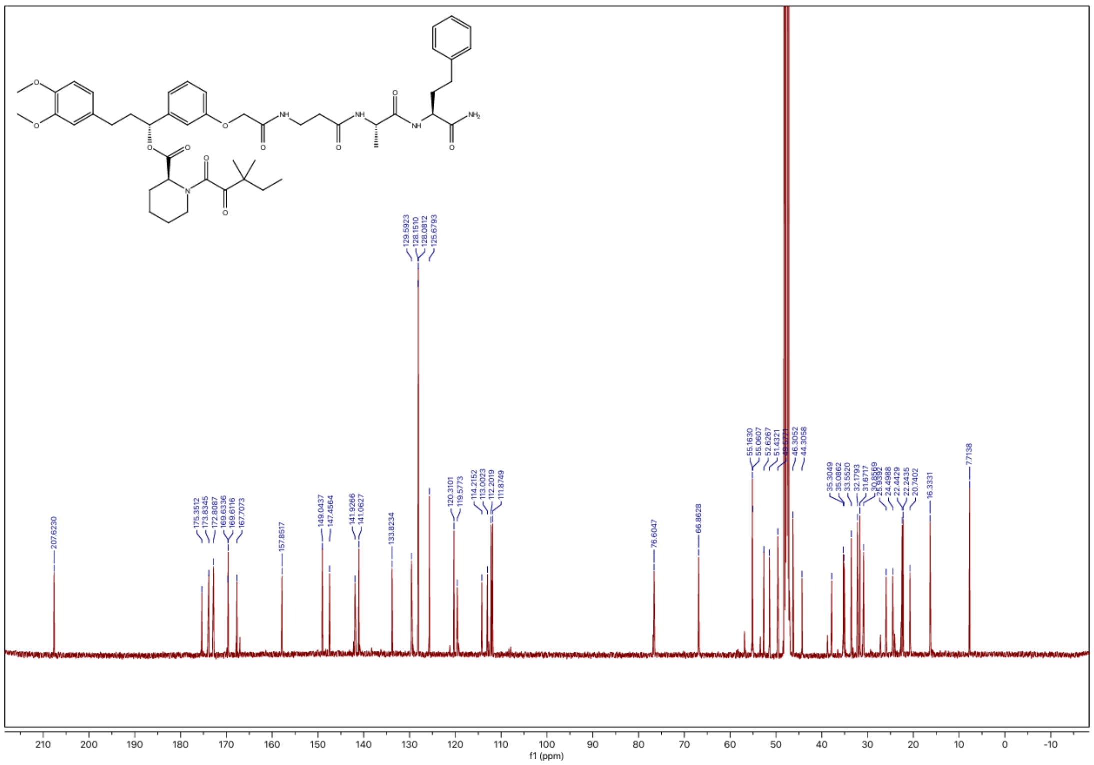

#### $^1$H NMR spectrum of JZ15

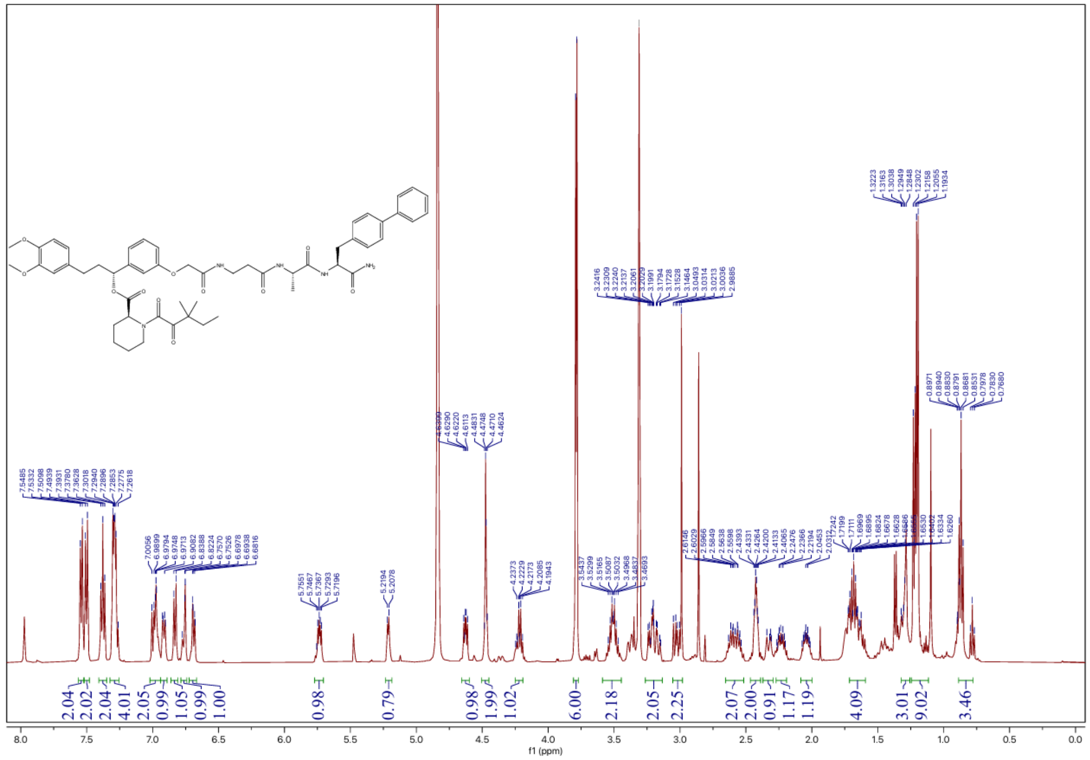

#### $^{13}$C NMR spectrum of JZ15

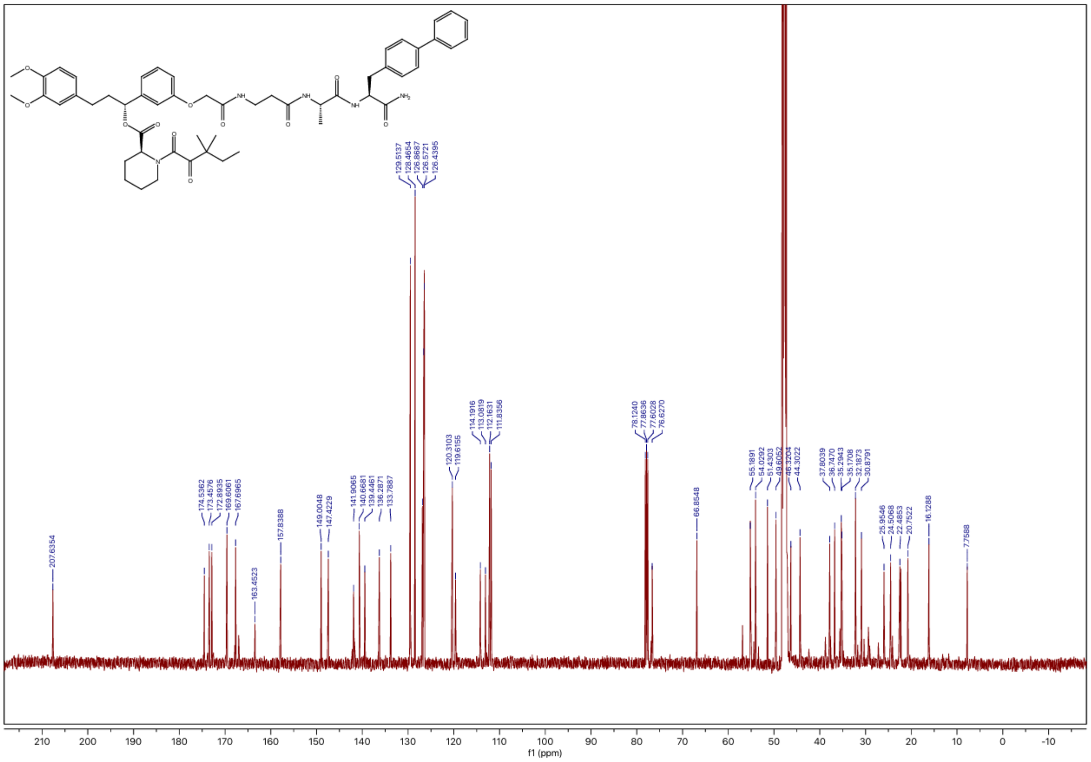

### $^{1}H$ NMR spectrum of JZ16

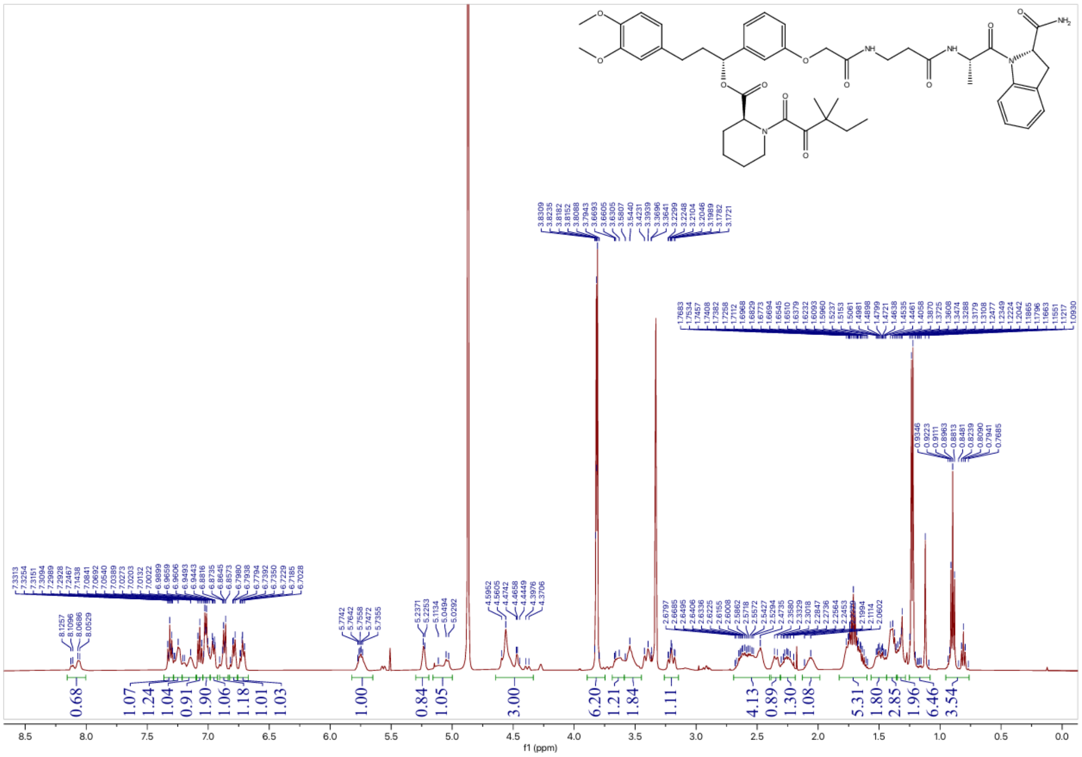


### $^{13}C$ NMR spectrum of JZ16

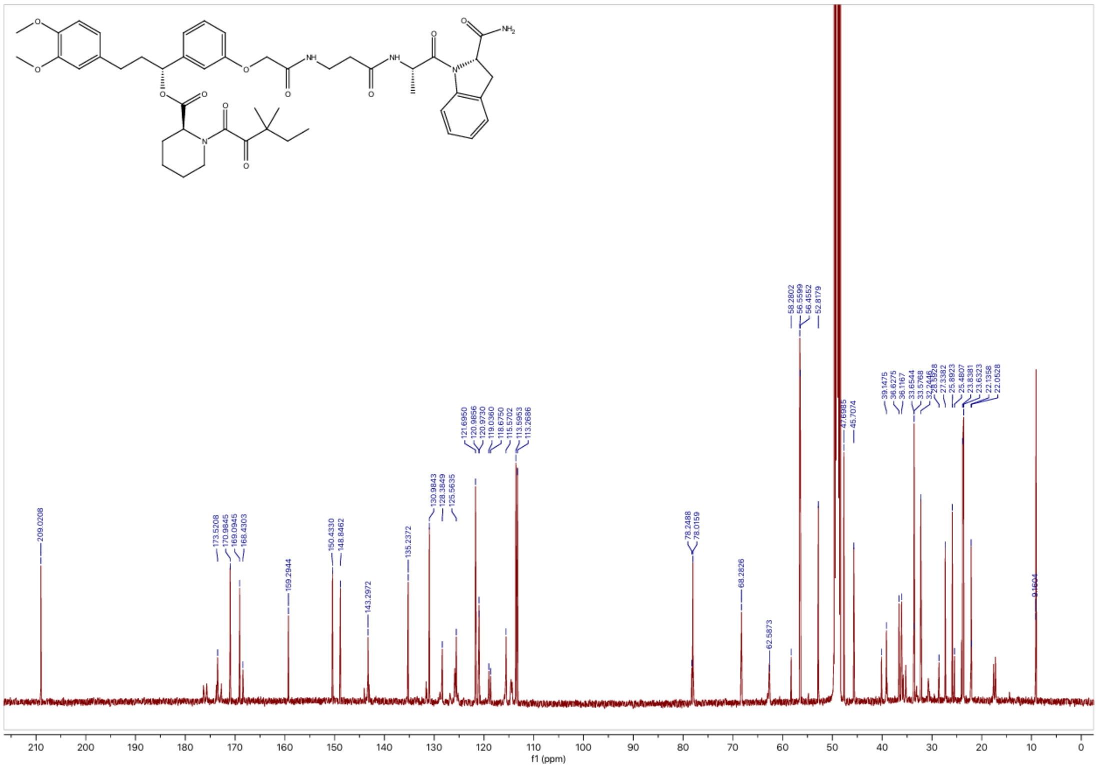

### $^{1}$H NMR spectrum of JZ17

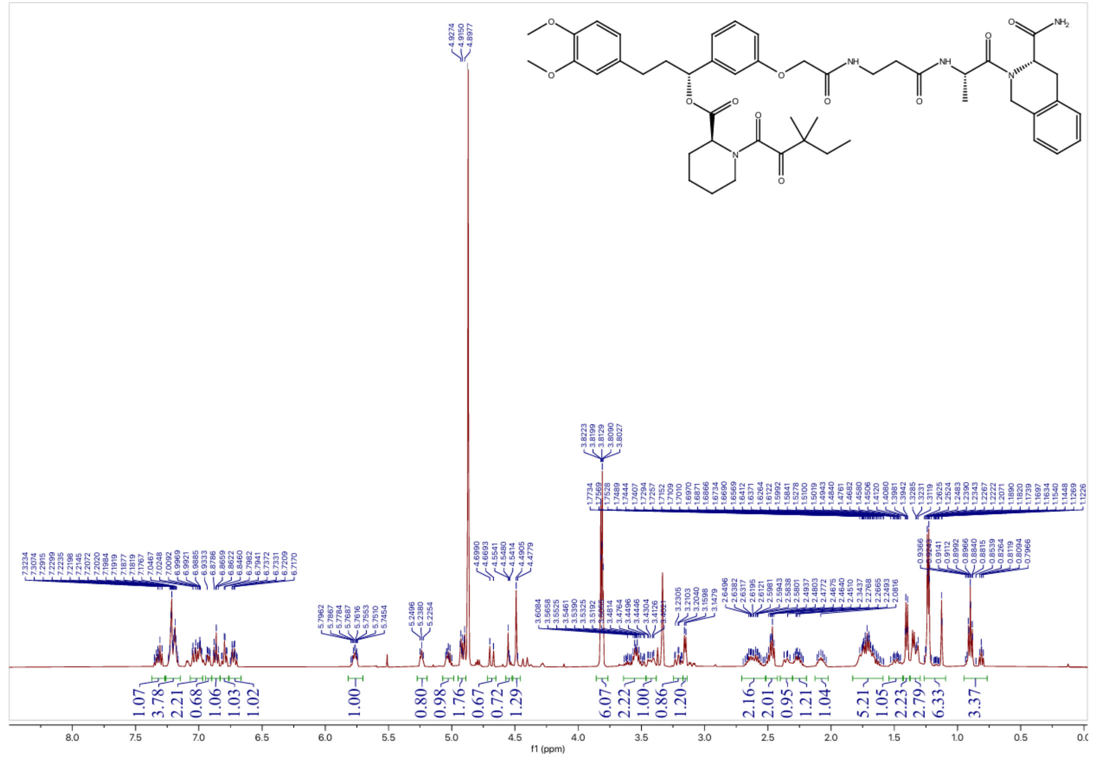

### $^{13}$C NMR spectrum of JZ17

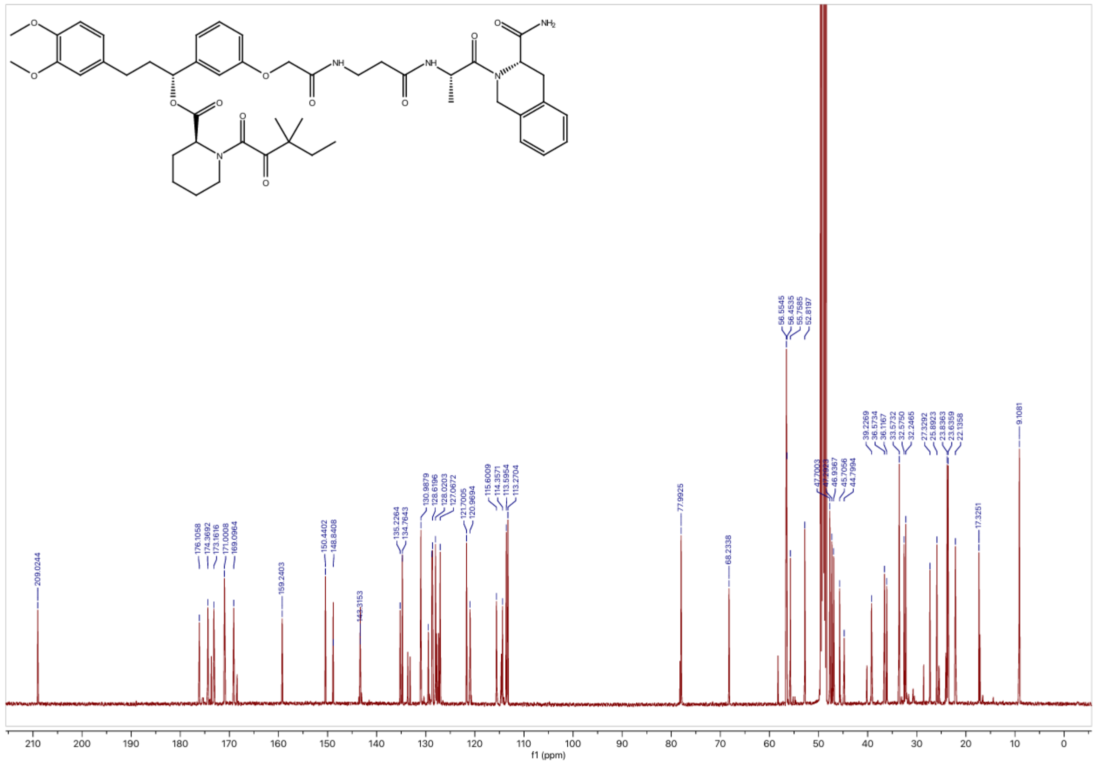

### $^{1}H$ NMR spectrum of JZ18

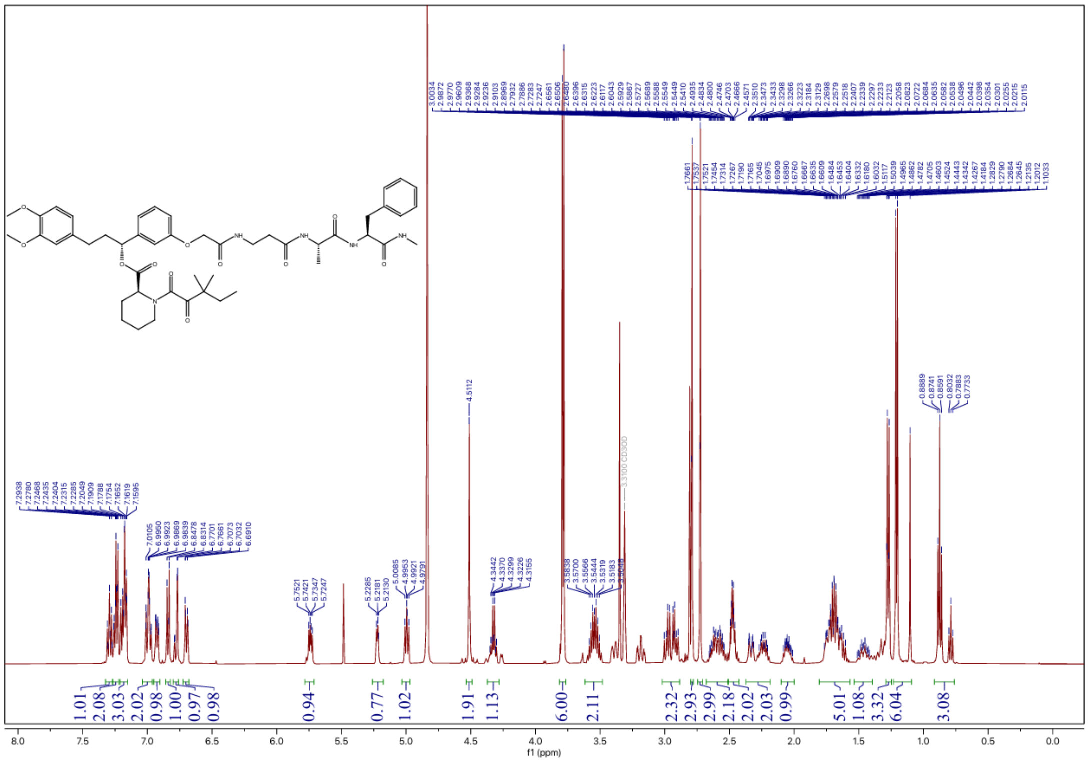

### $^{13}C$ NMR spectrum of JZ18

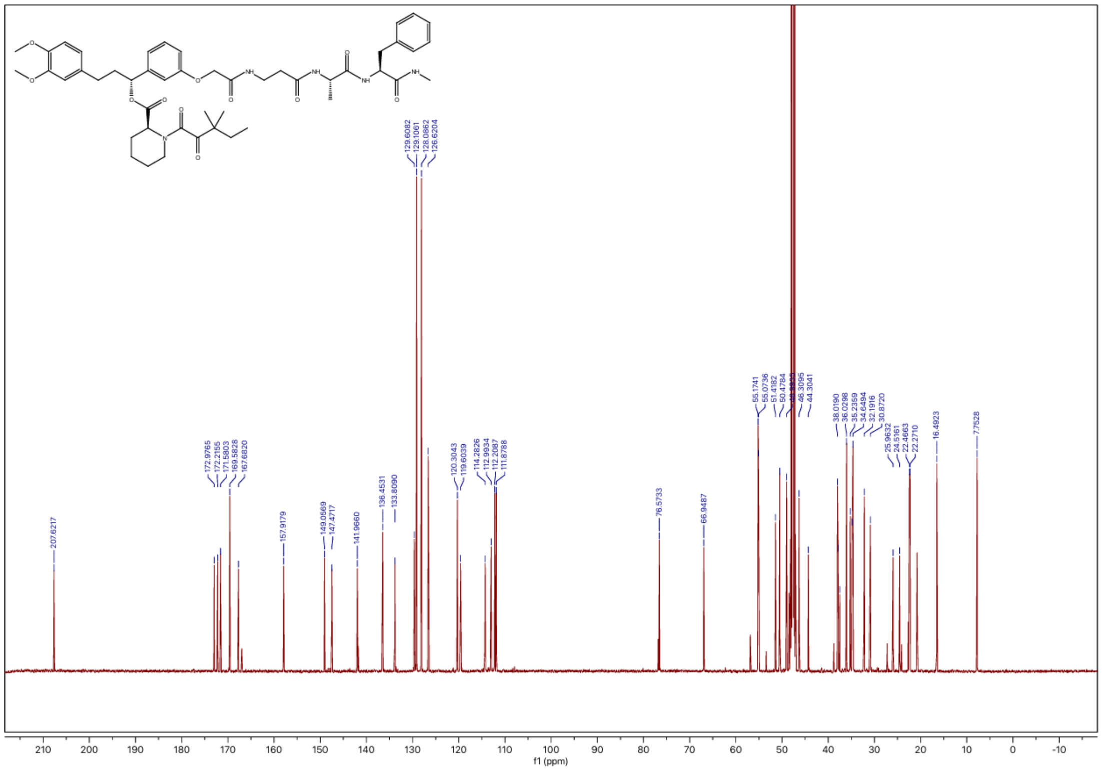

### $^{1}$H NMR spectrum of JZ19

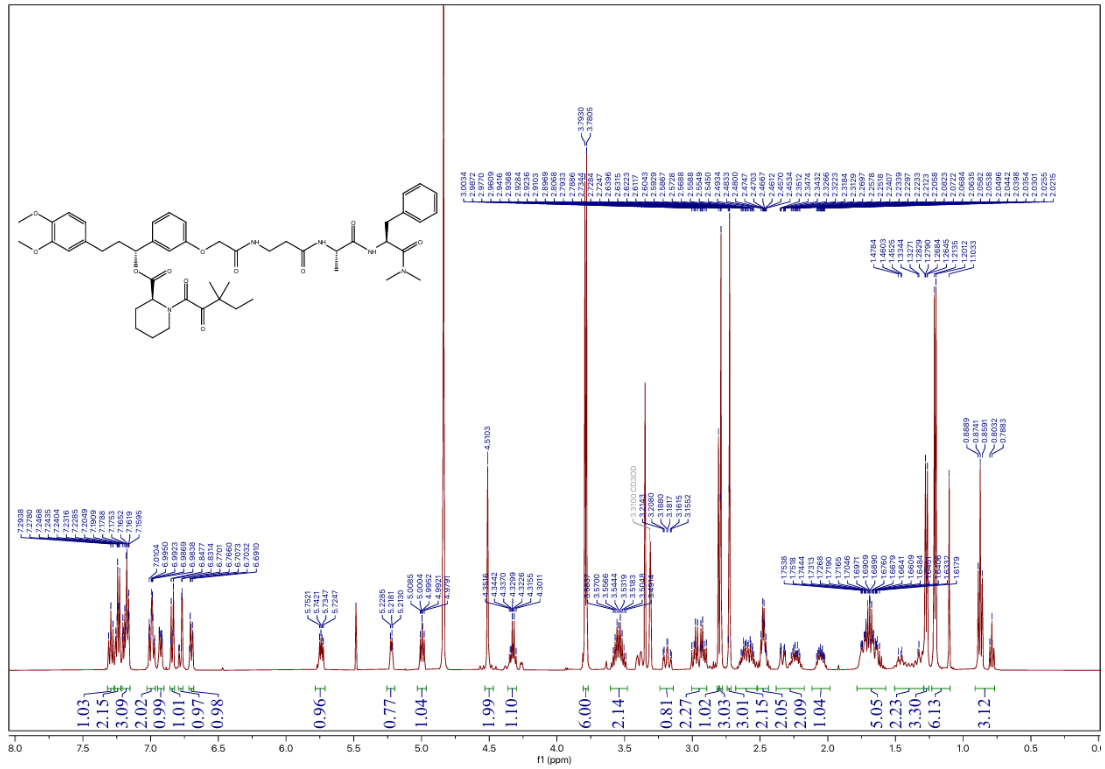

### $^{13}$C NMR spectrum of JZ19

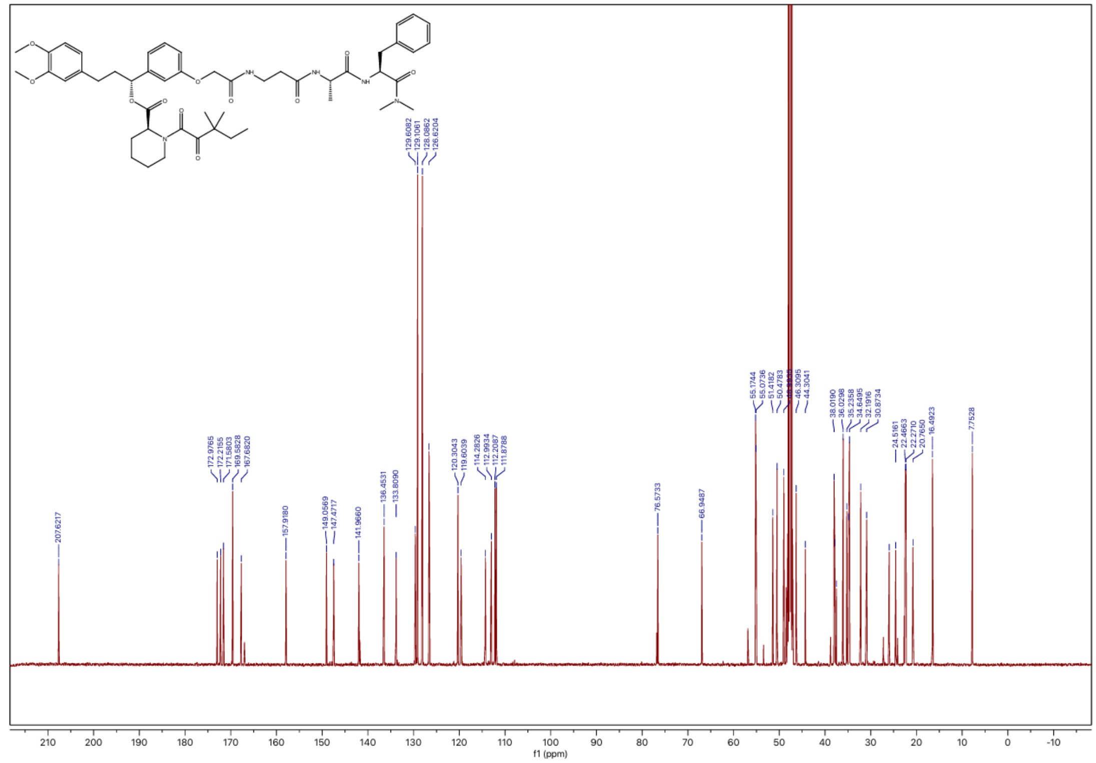

### $^{1}$H NMR spectrum of JZ20

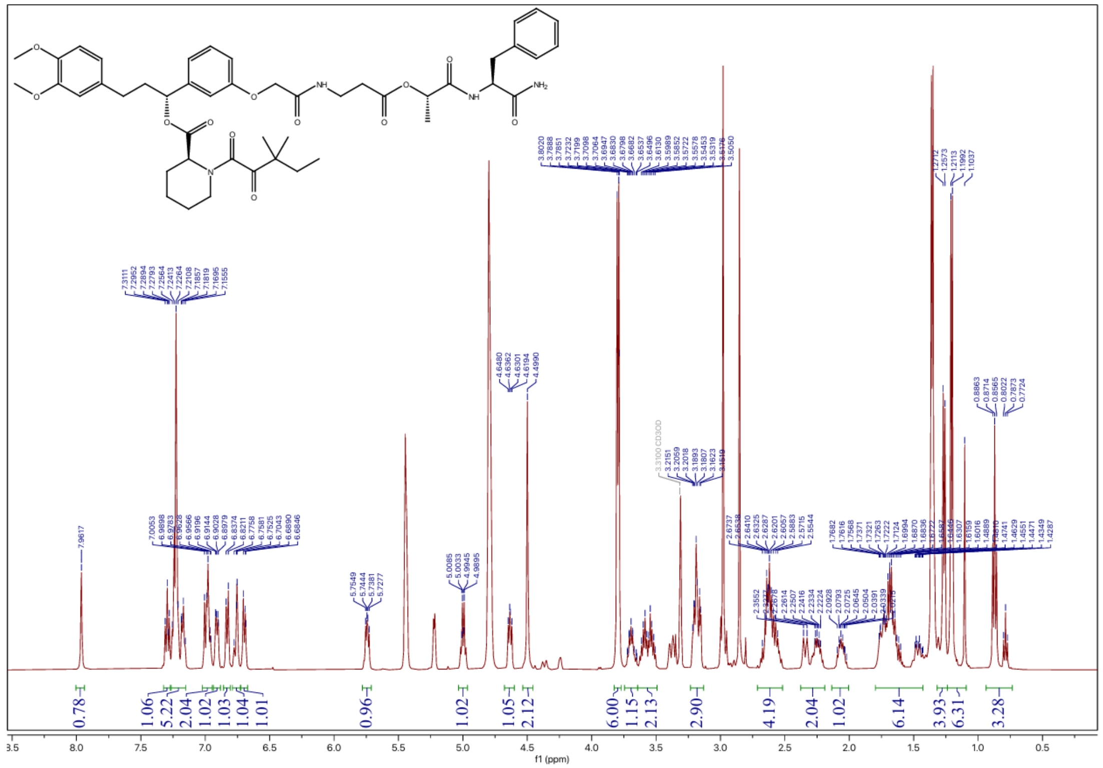

### $^{13}$C NMR spectrum of JZ20

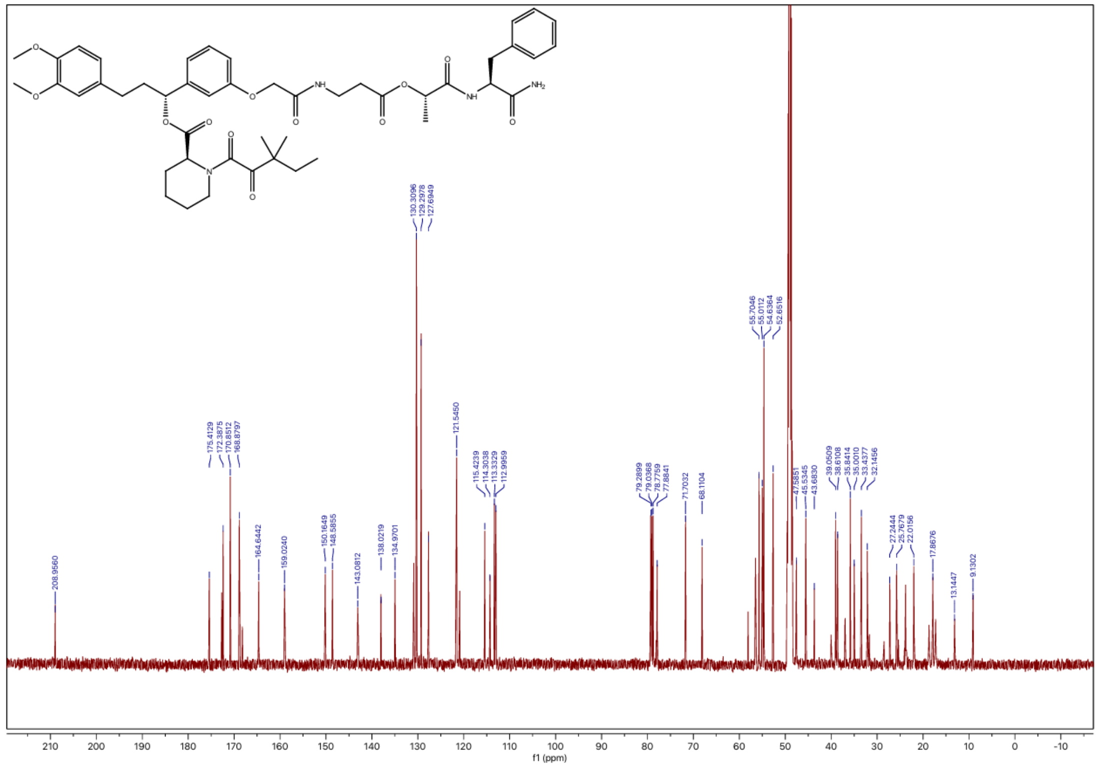

### $^{1}$H NMR spectrum of JZ21

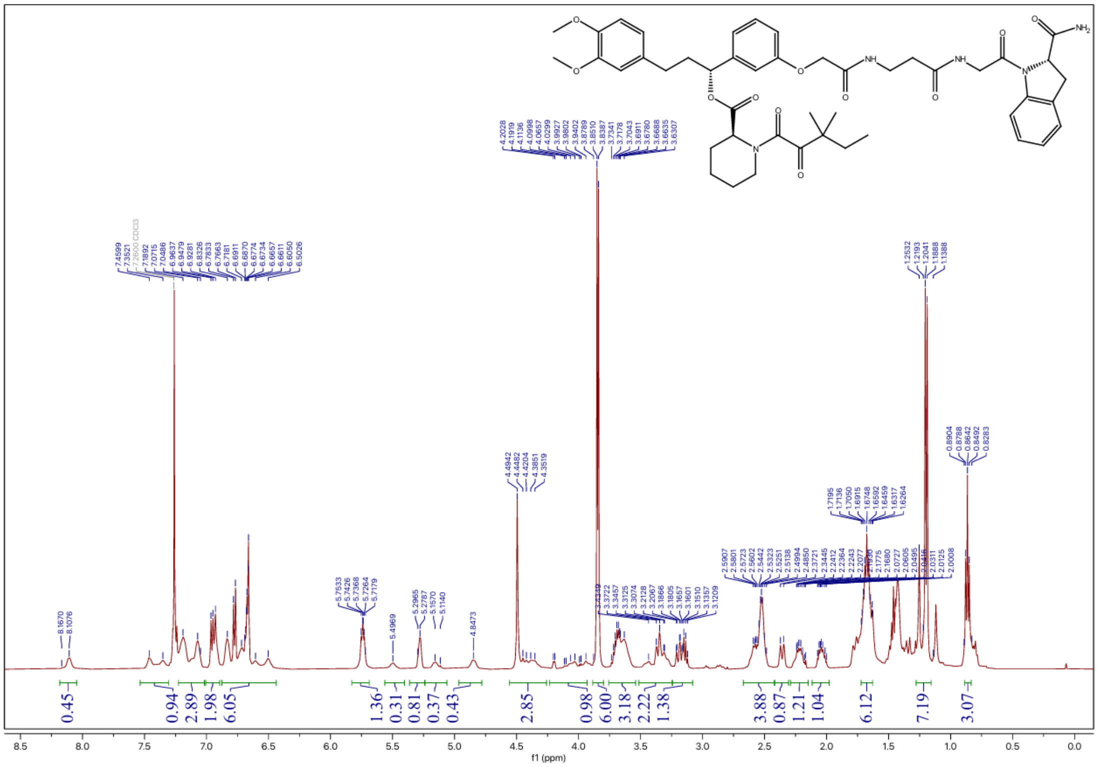

### $^{13}$C NMR spectrum of JZ21

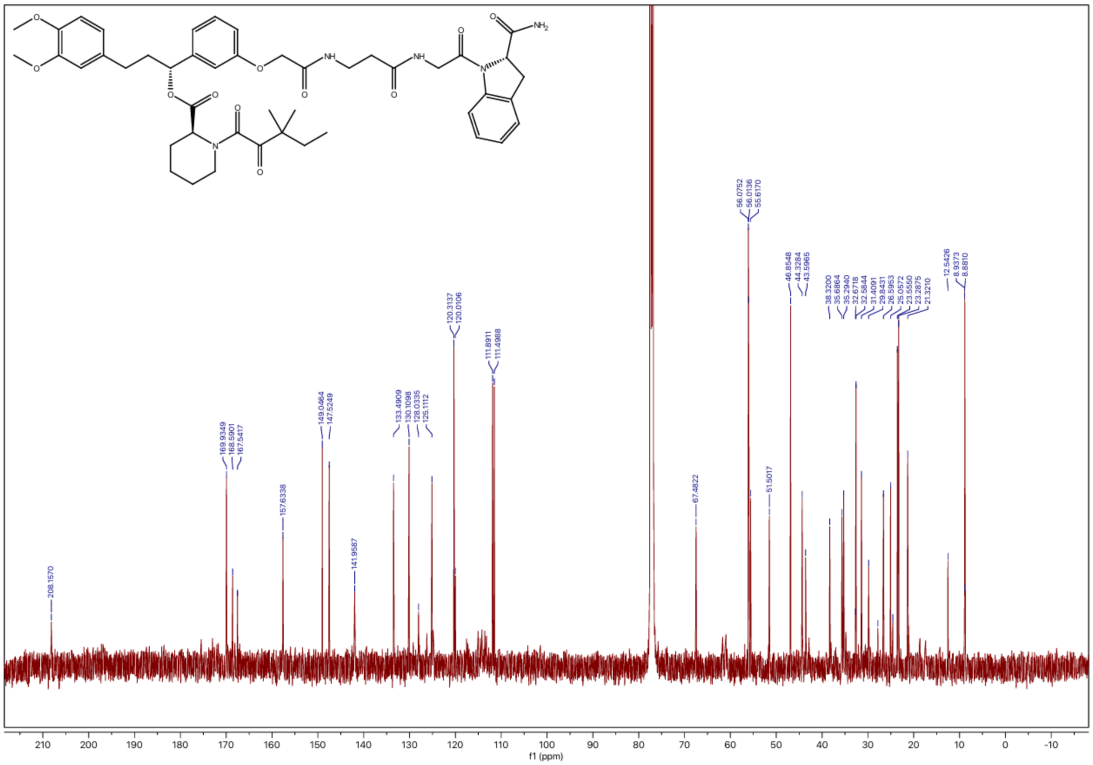

### $^1$H NMR spectrum of JZ22

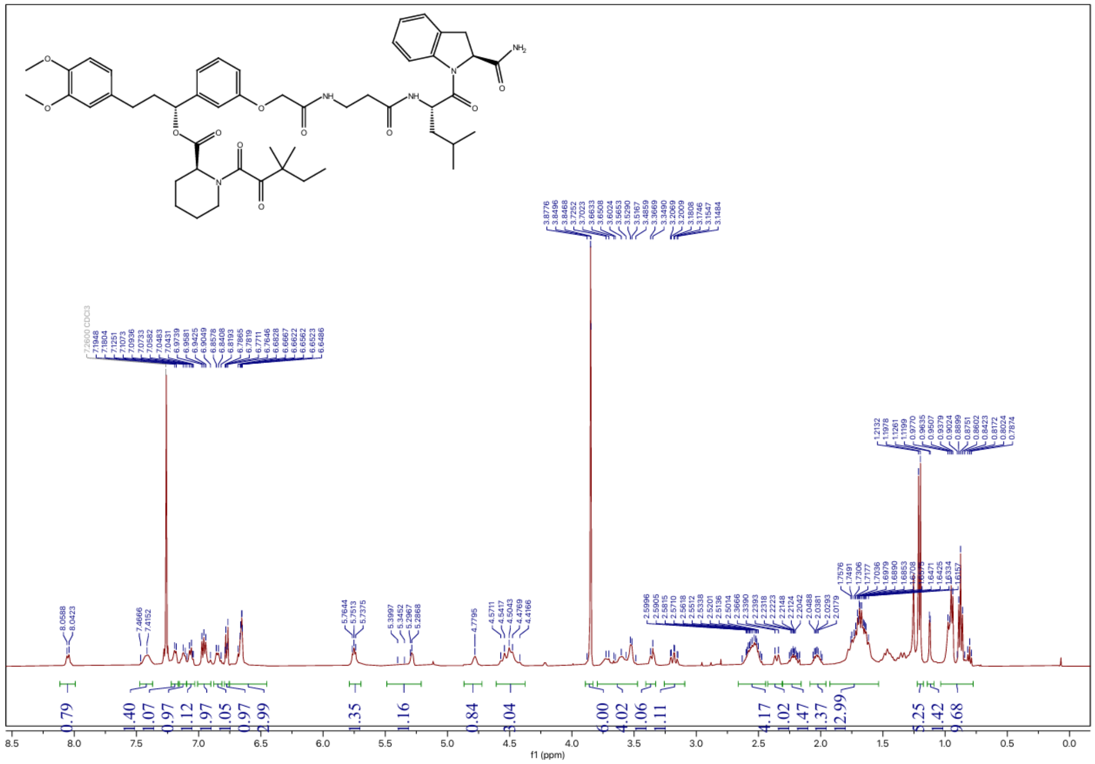

### $^{13}$C NMR spectrum of JZ22

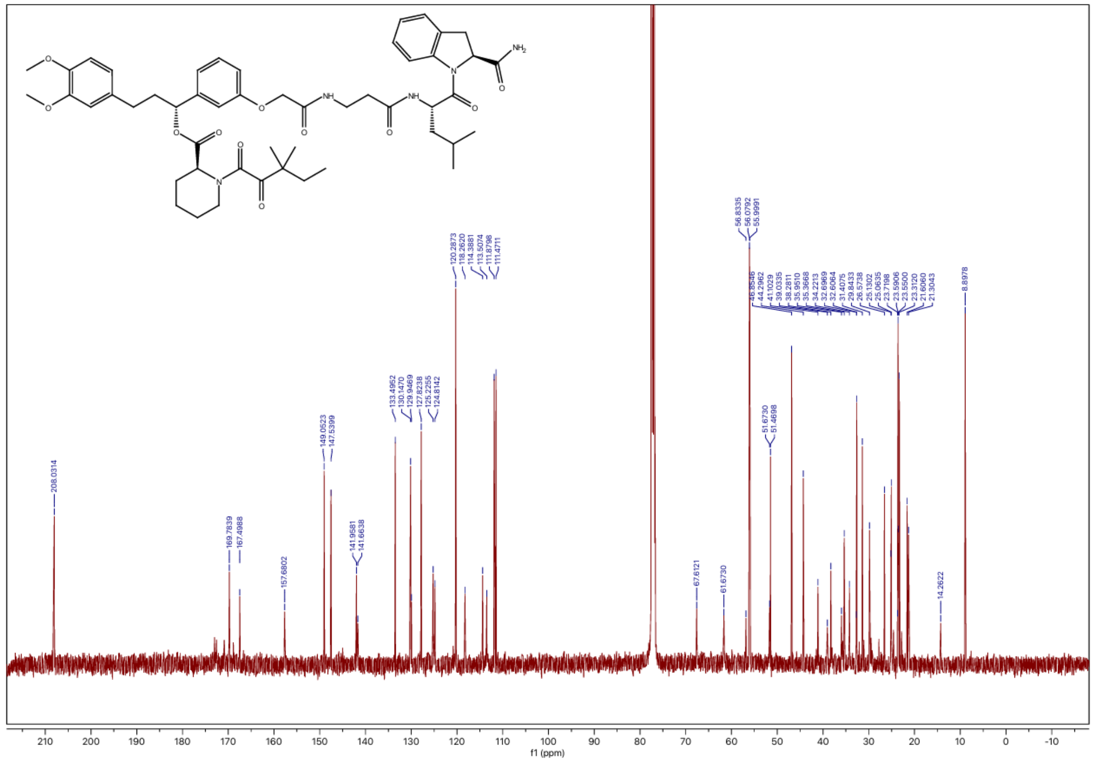

### $^{1}$H NMR spectrum of JZ23

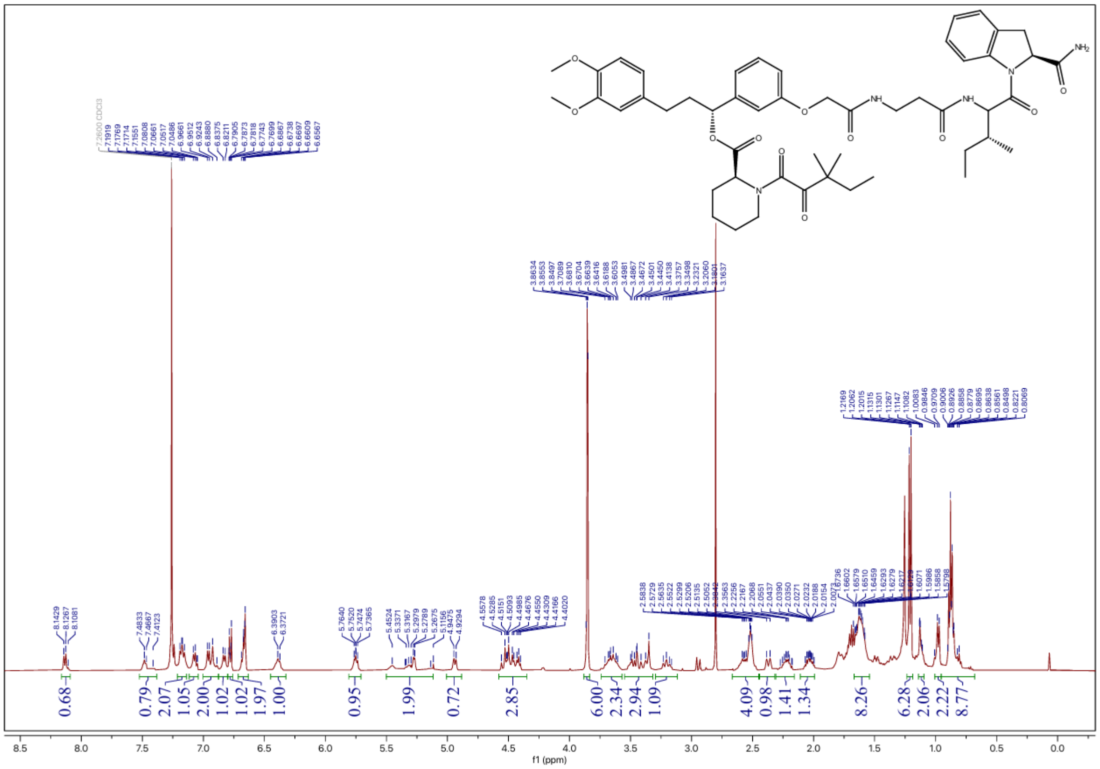

### $^{13}$C NMR spectrum of JZ23

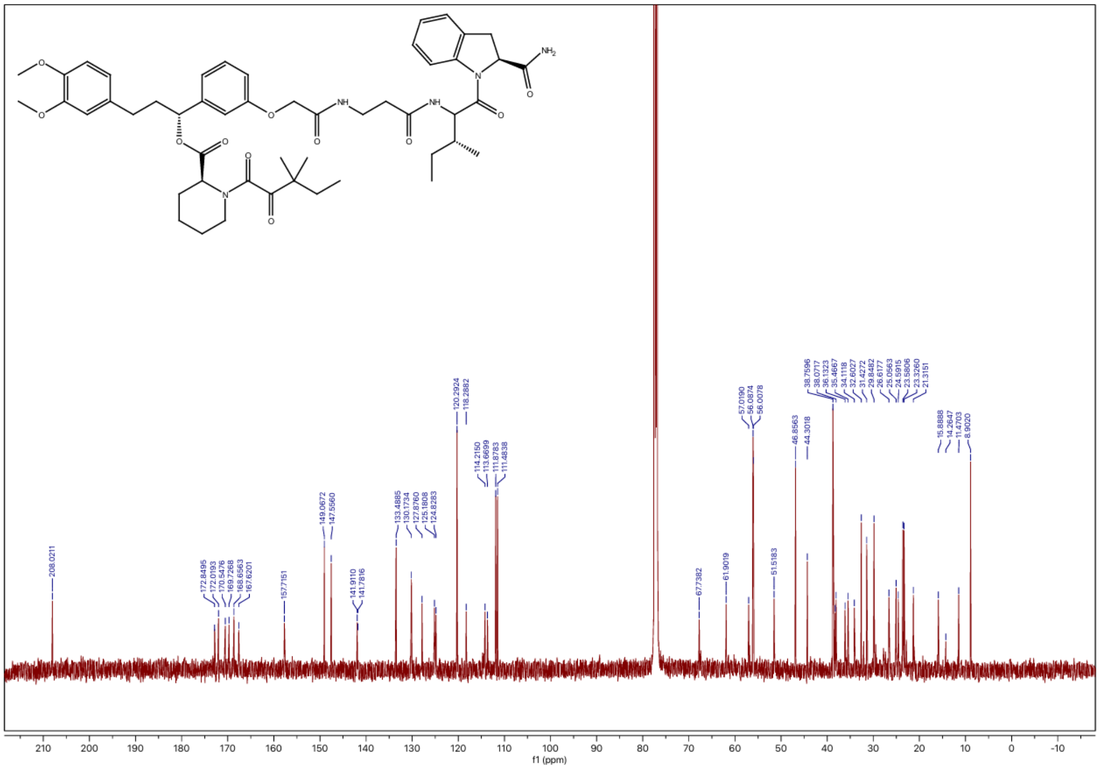

### $^{1}$H NMR spectrum of JZ24

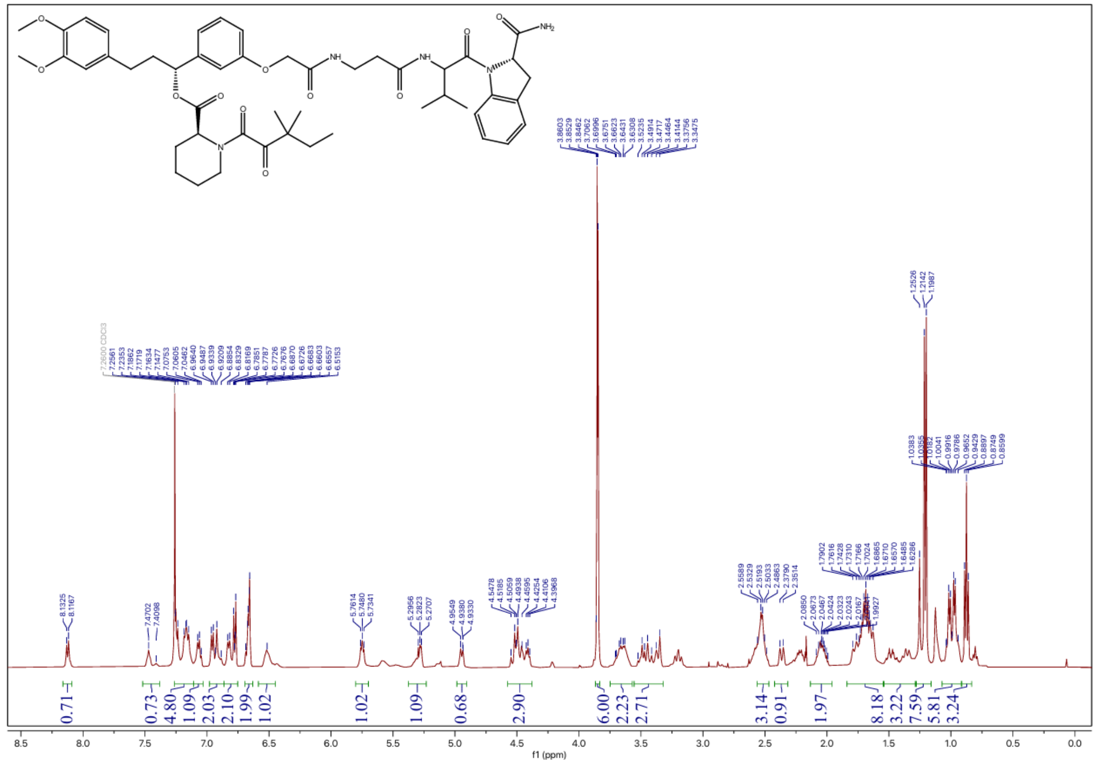

### $^{13}$C NMR spectrum of JZ24

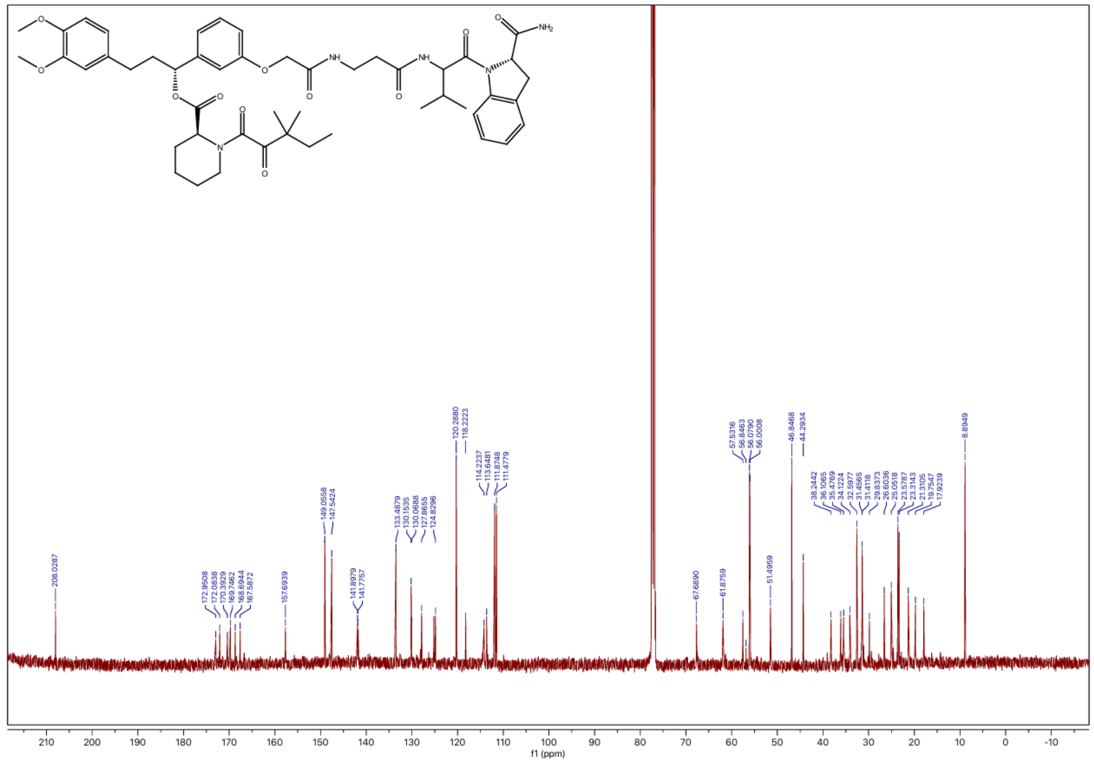

**$^{1}$H NMR spectrum of JZ25**

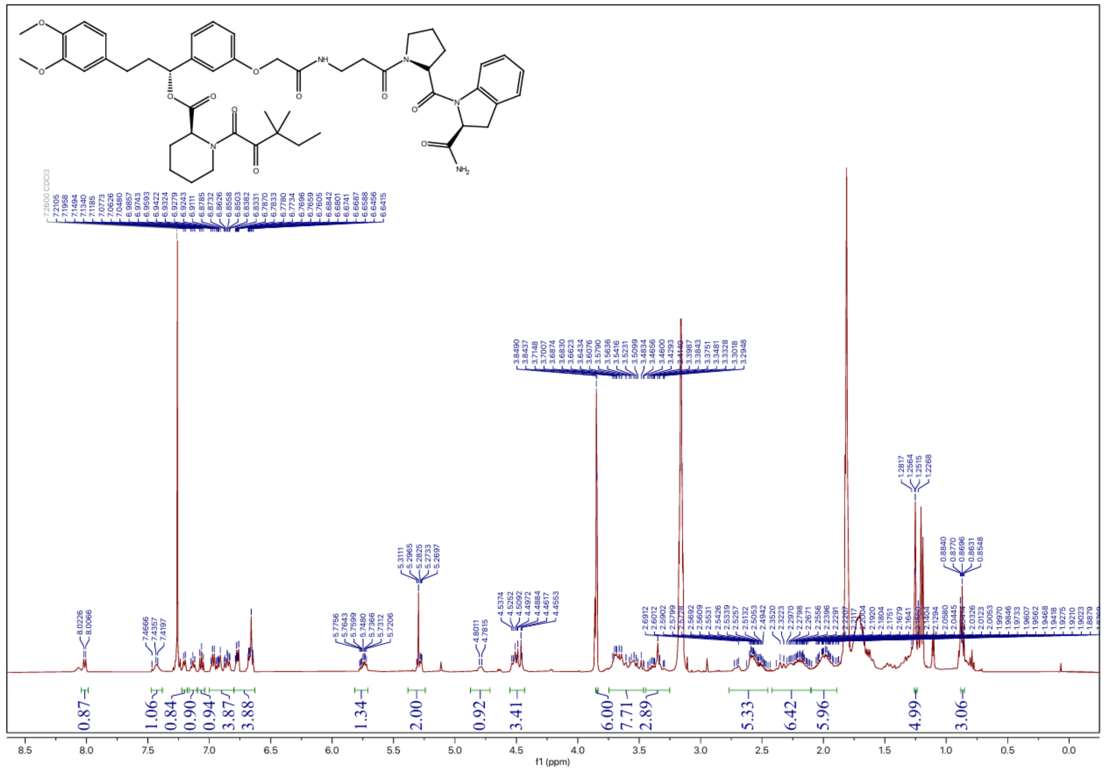

**$^{13}$C NMR spectrum of JZ25**

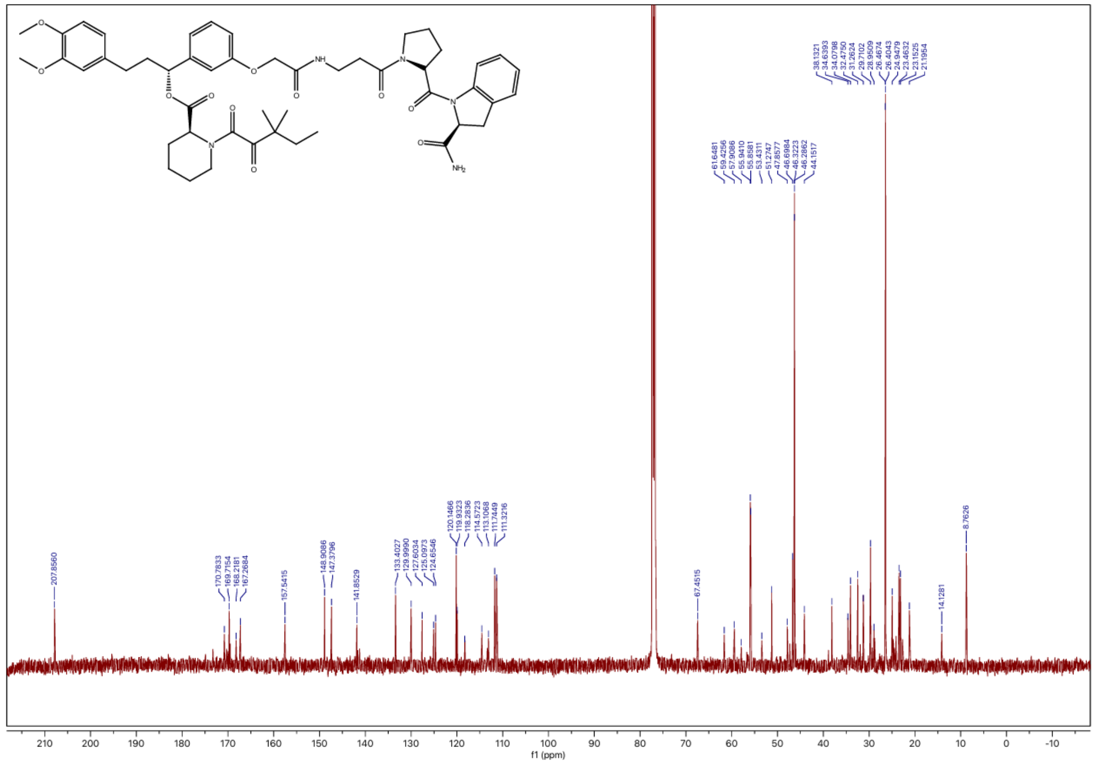

### $^{1}$H NMR spectrum of JZ26

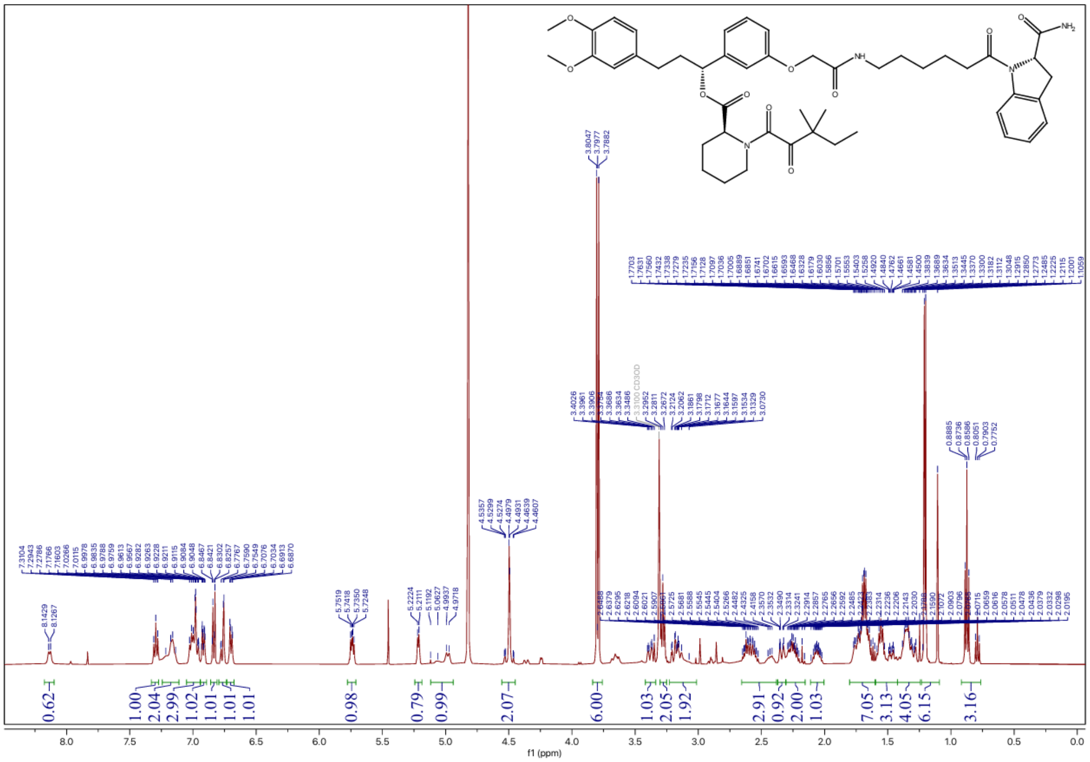


### $^{13}$C NMR spectrum of JZ26

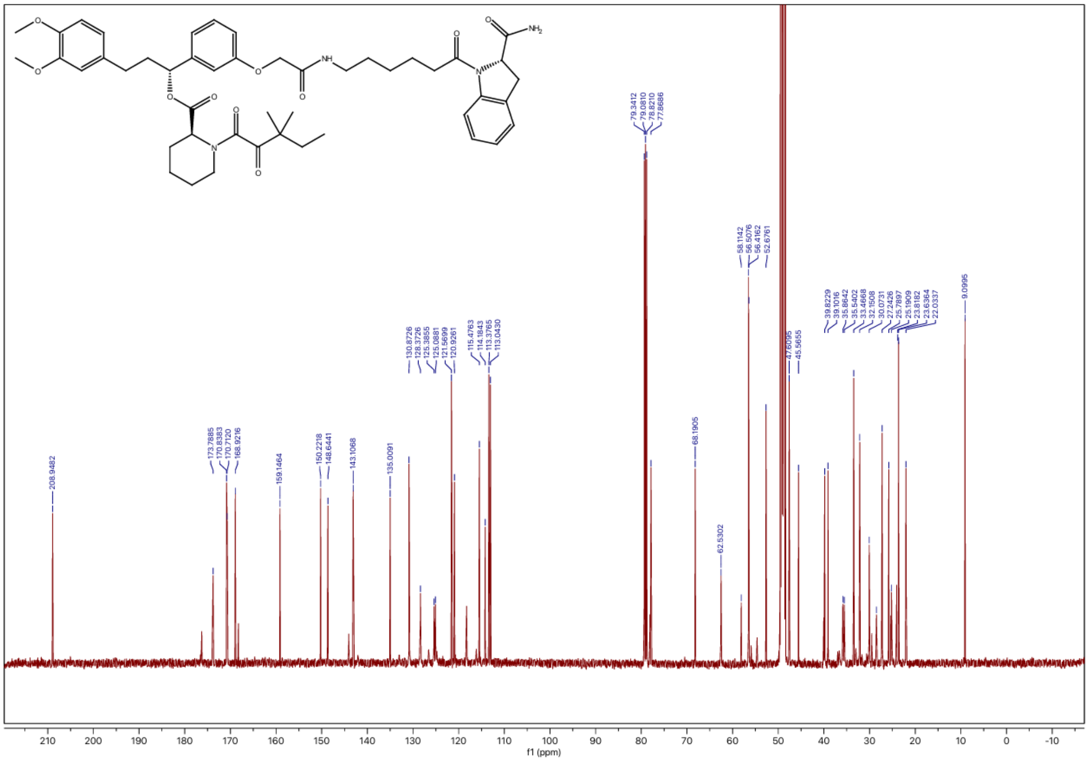

### $^{1}H$ NMR spectrum of JZ27

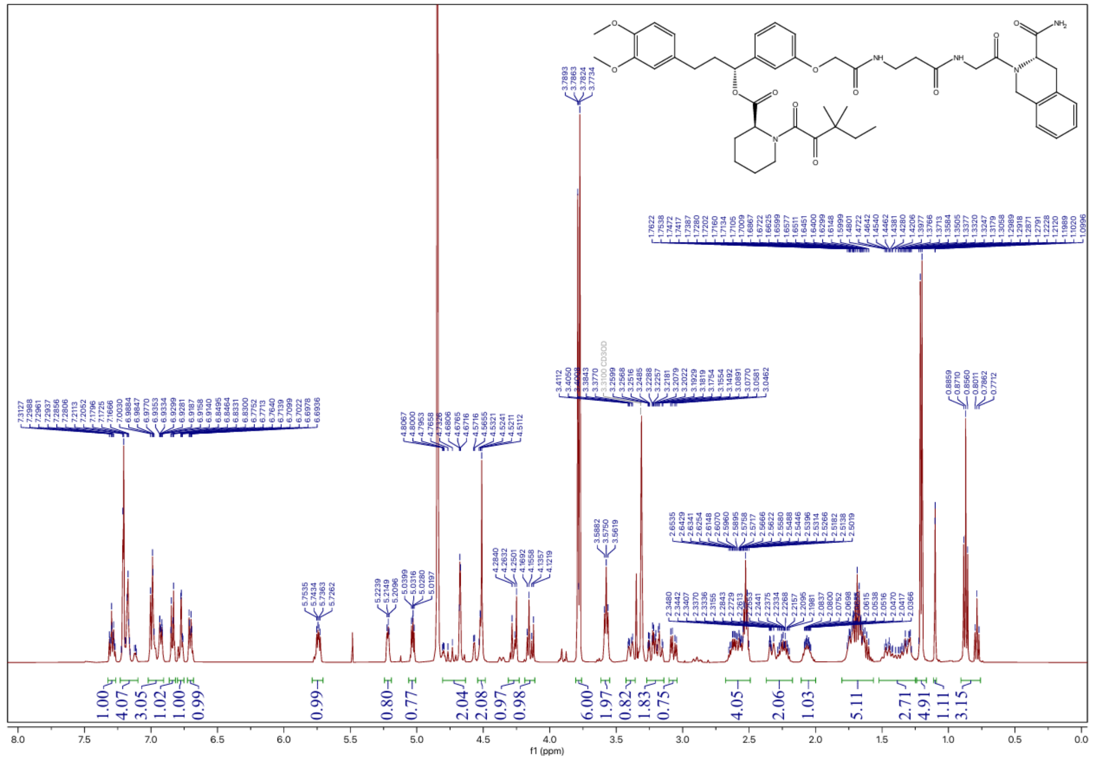

### $^{13}C$ NMR spectrum of JZ27

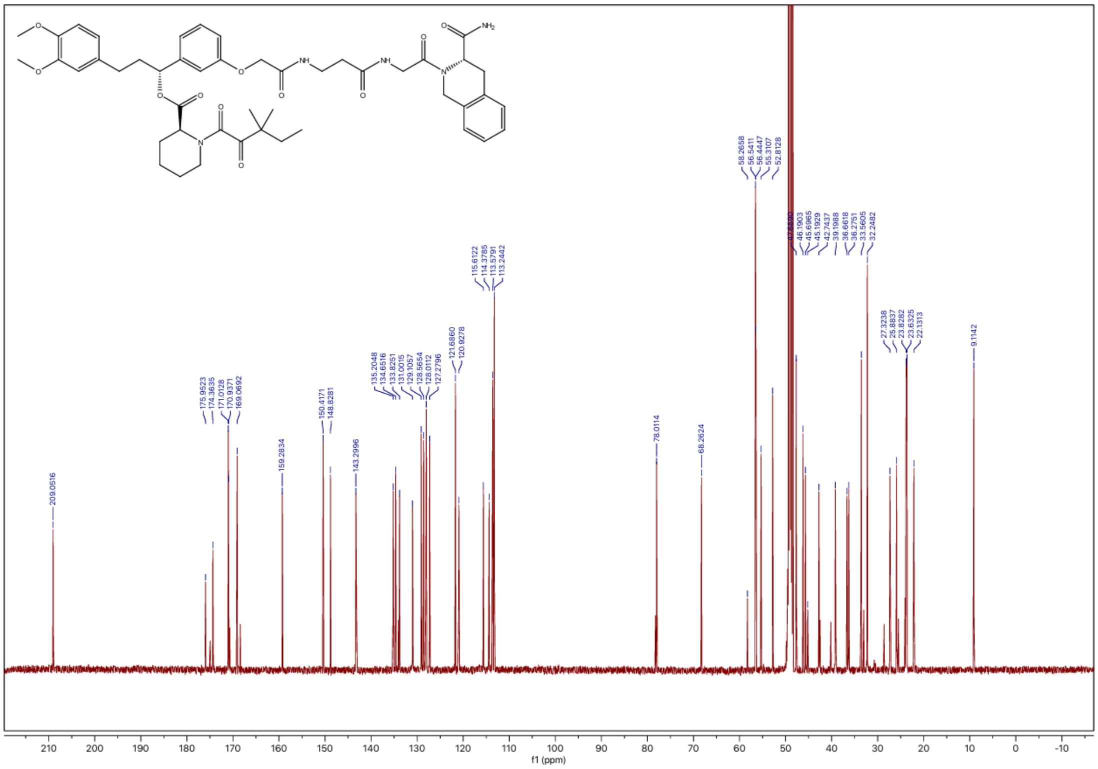

### $^{1}H$ NMR spectrum of JZ28

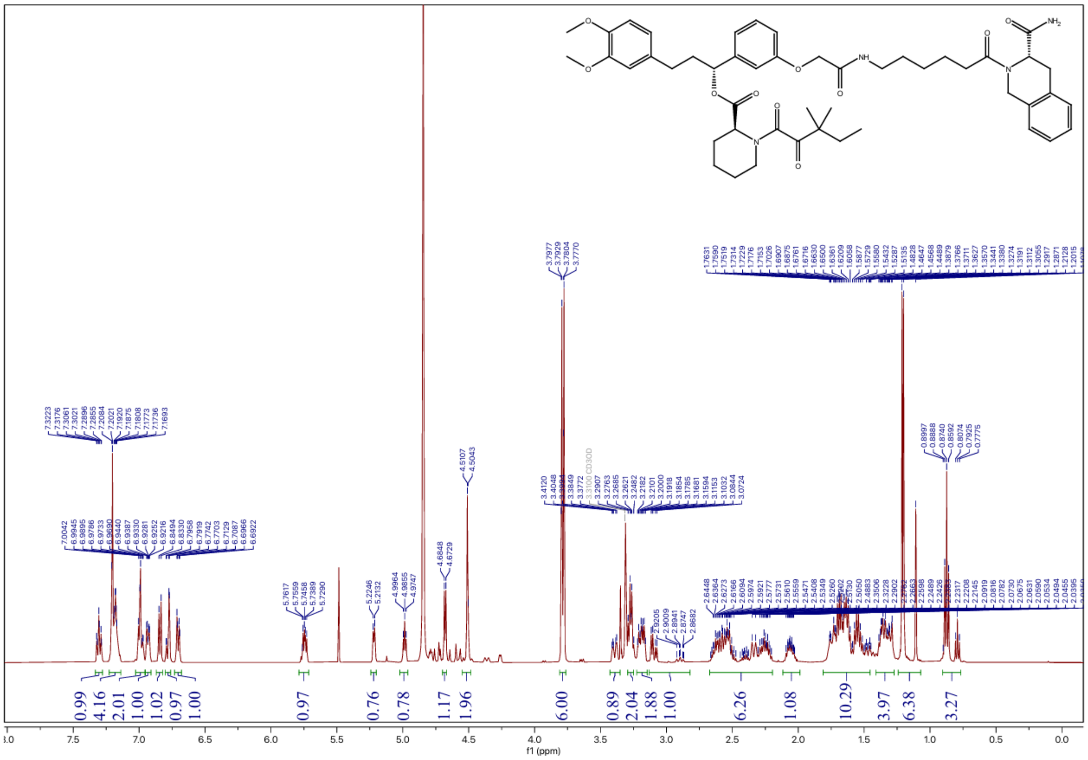

### $^{13}C$ NMR spectrum of JZ28

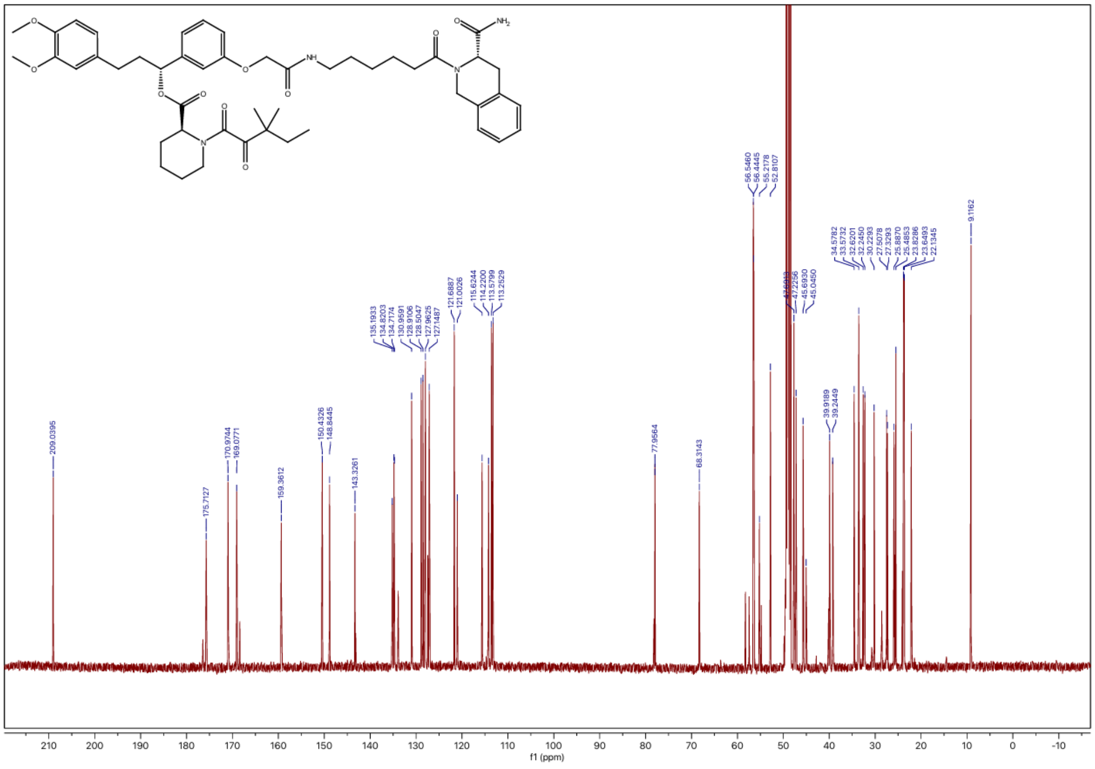

**$^1$H NMR spectrum of JZ29**

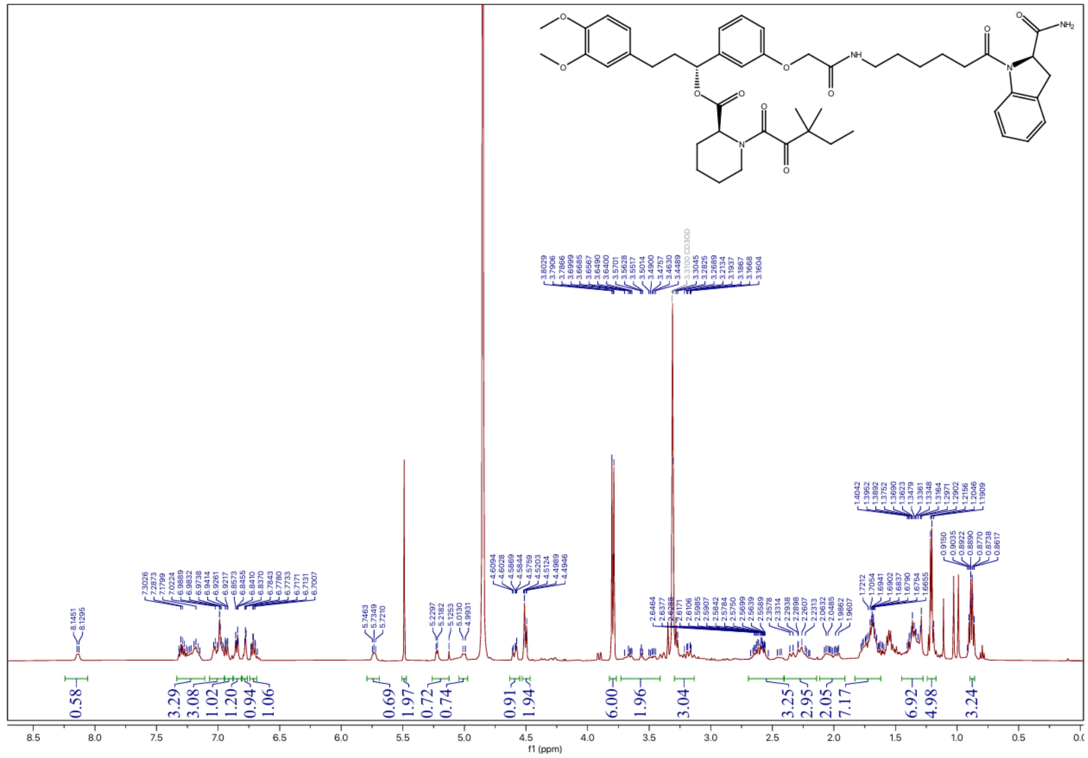

**$^{13}$C NMR spectrum of JZ29**

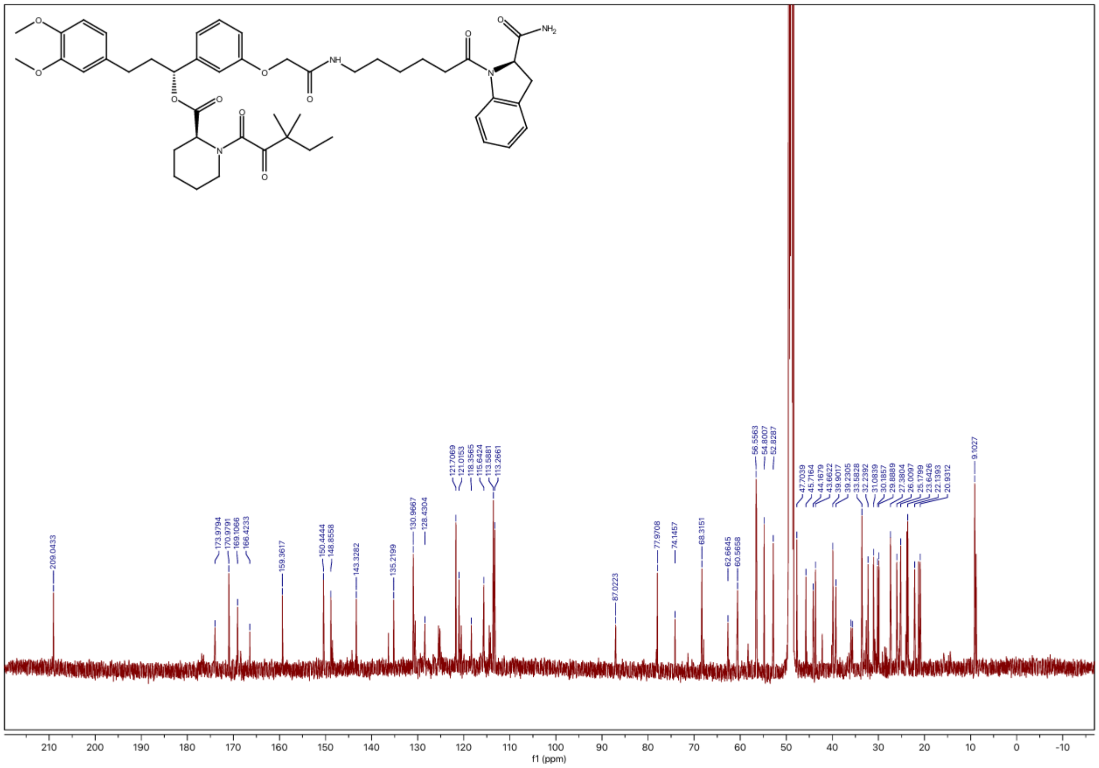

### $^1$H NMR spectrum of JZ30

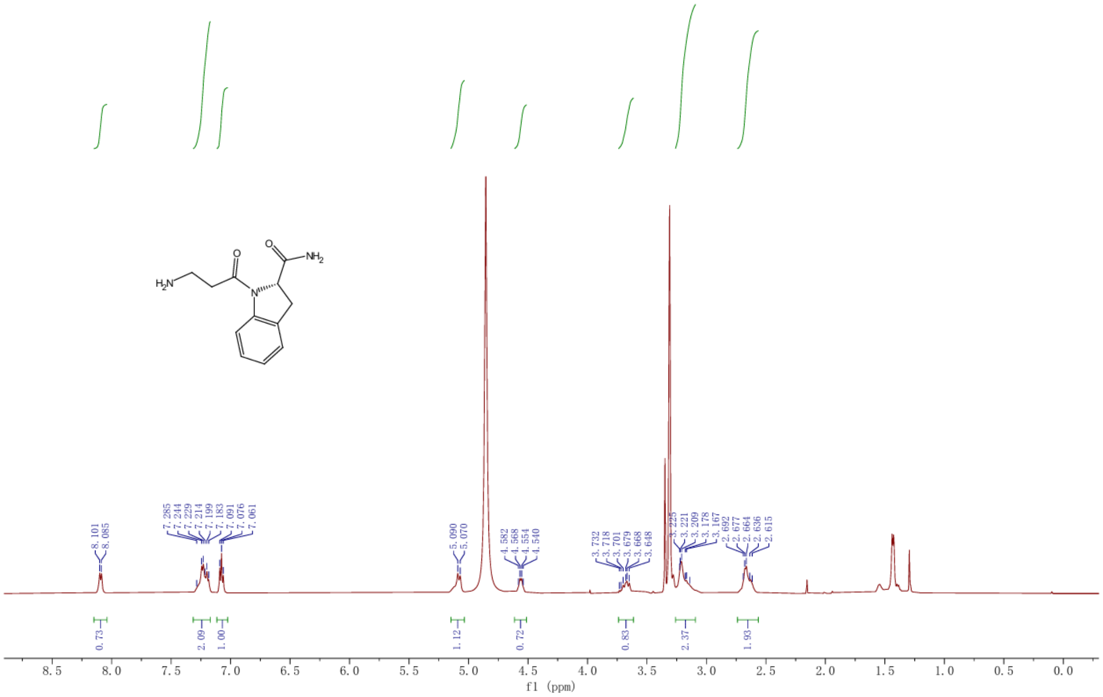

### $^1$H NMR spectrum of JZ31

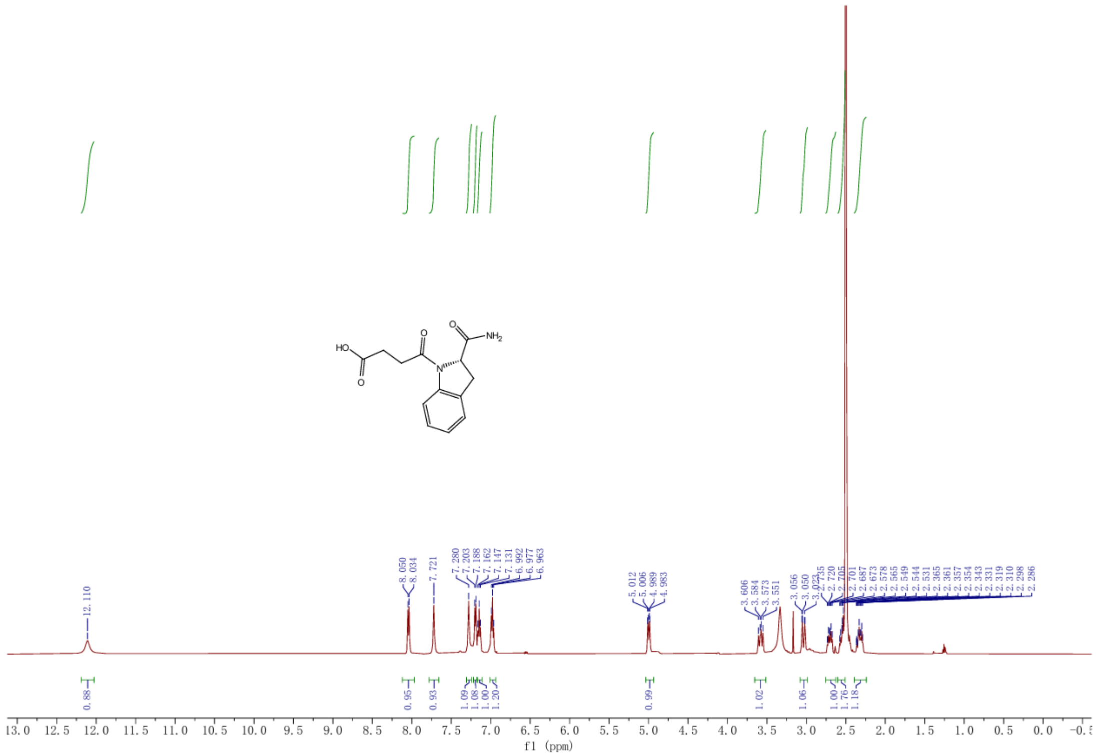

### $^{13}$C NMR spectrum of JZ31

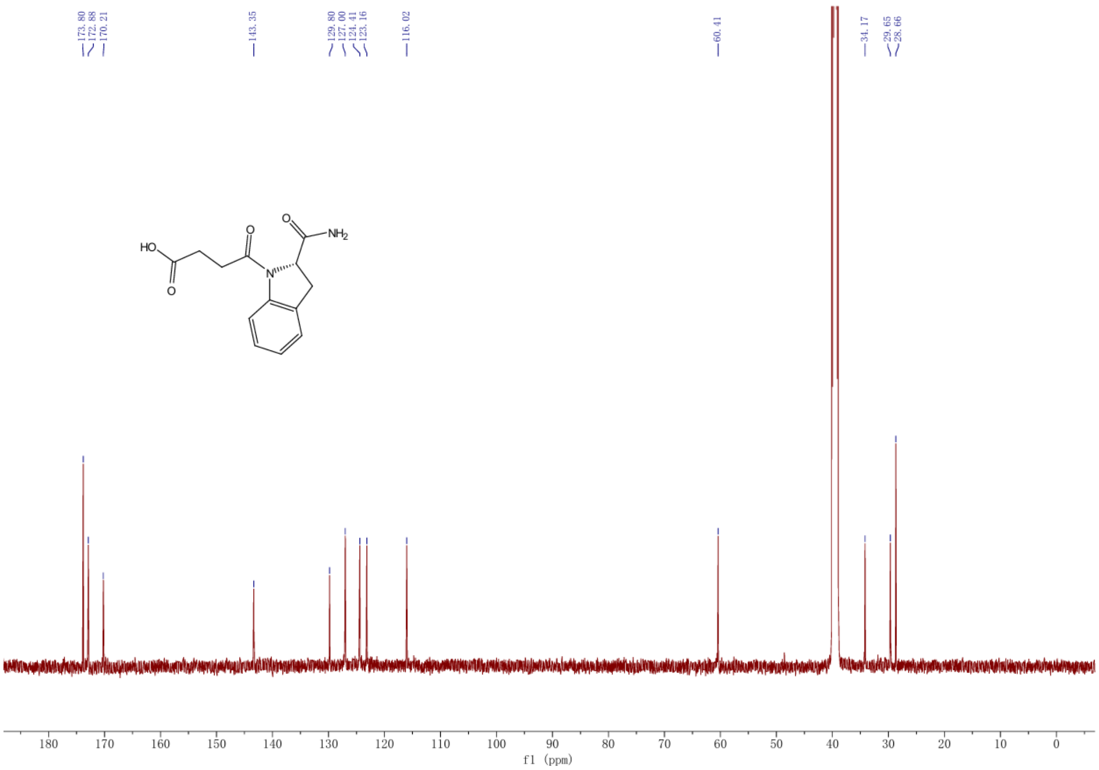

### $^{1}$H NMR spectrum of JZ32

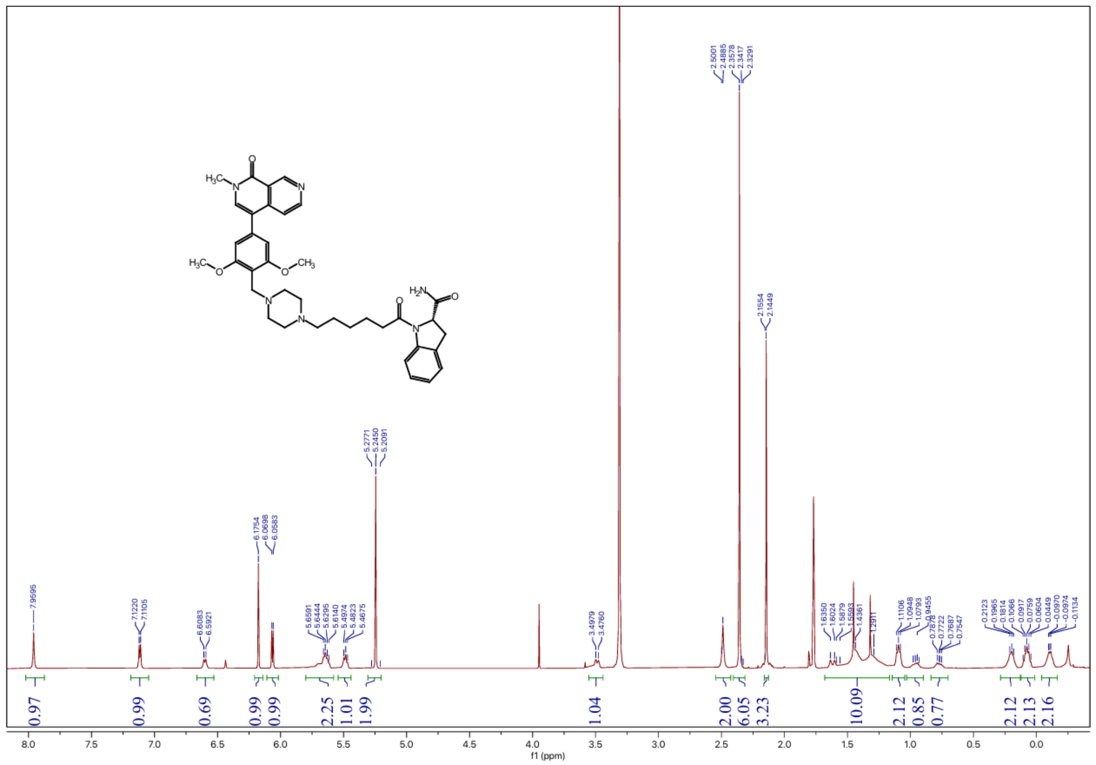

### $^{13}$C NMR spectrum of JZ32

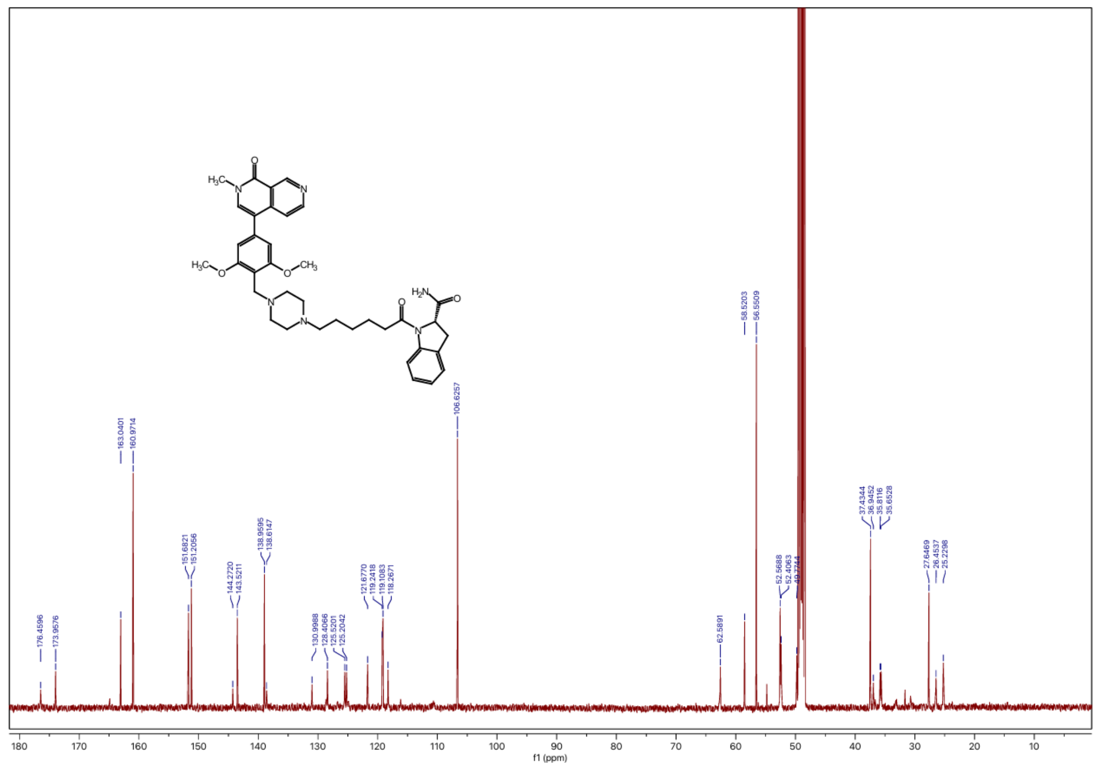

### $^1H$ NMR spectrum of JZ33

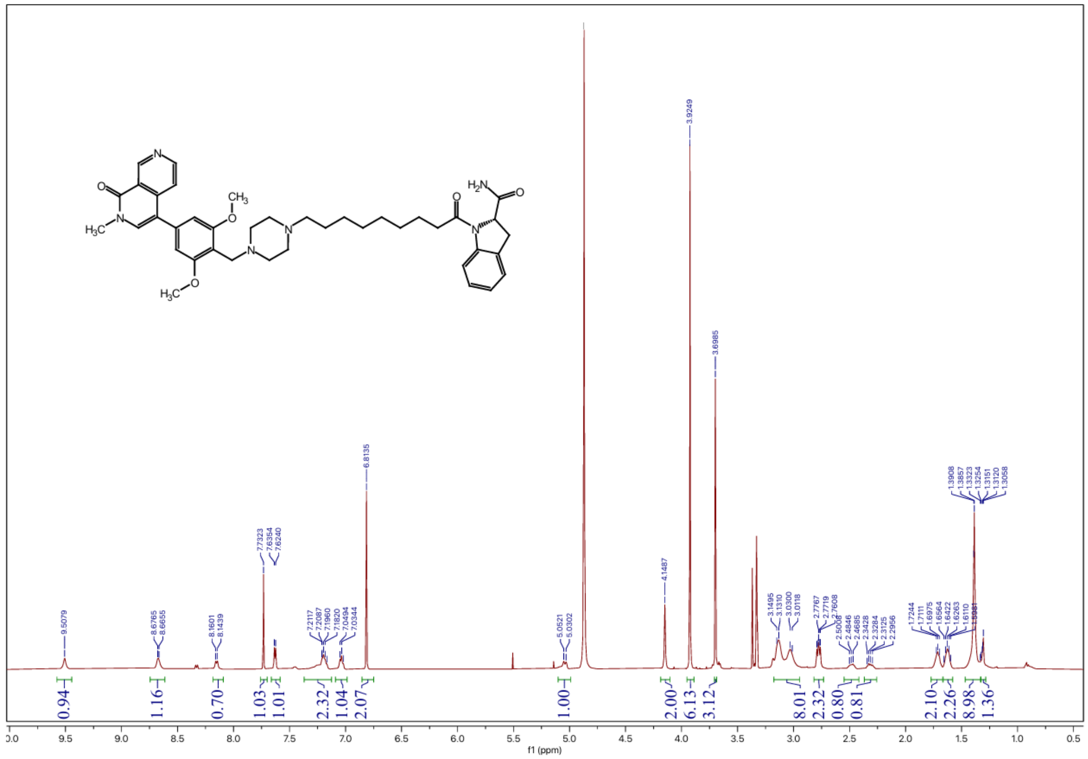


### $^{13}C$ NMR spectrum of JZ33

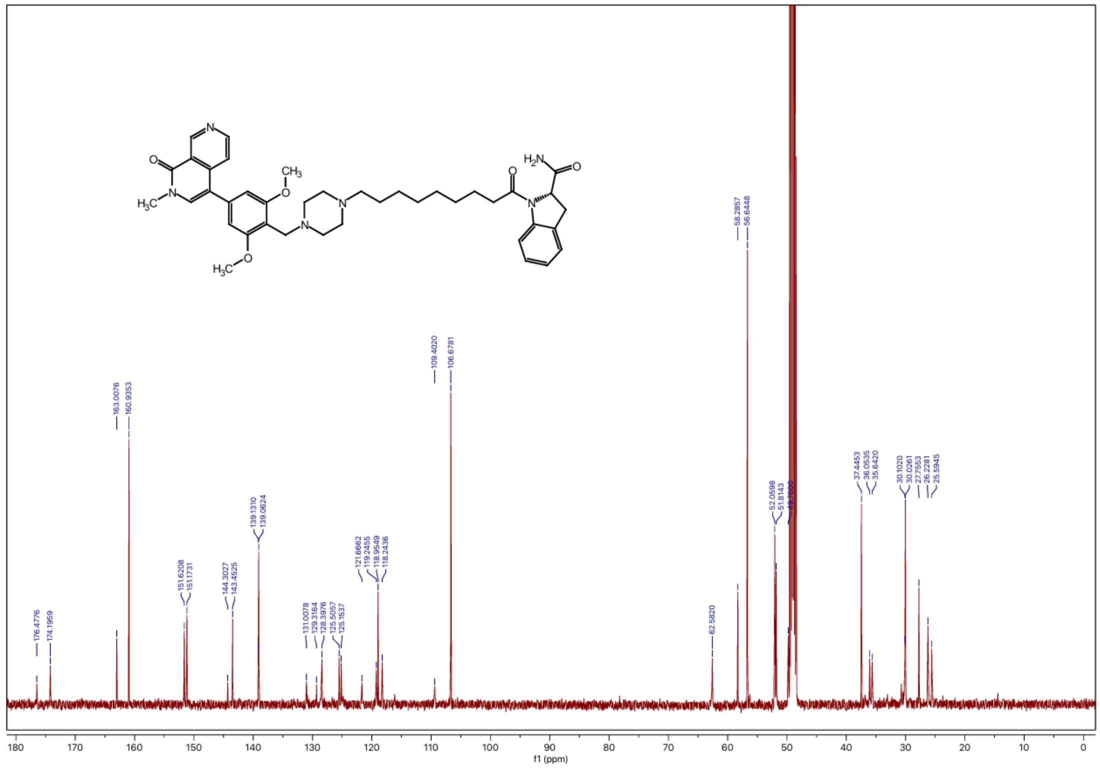

### $^{1}H$ NMR spectrum of JZ34

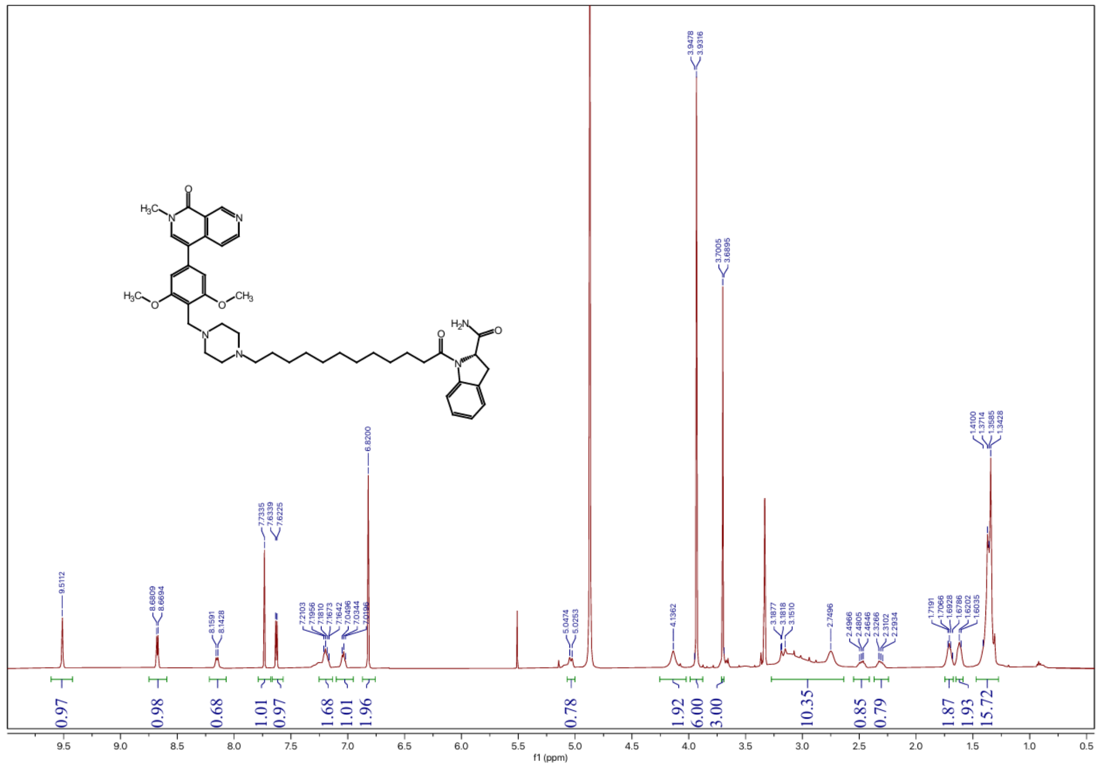

### $^{13}C$ NMR spectrum of JZ34

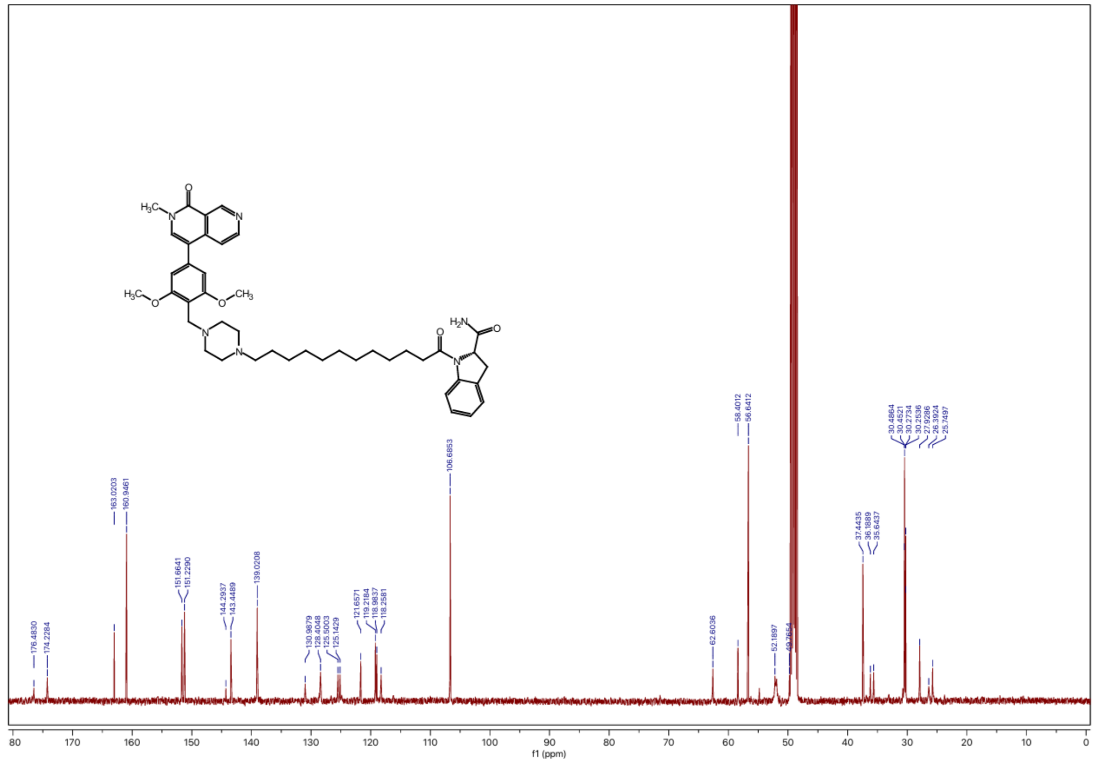

### $^{1}H$ NMR spectrum of JZ35

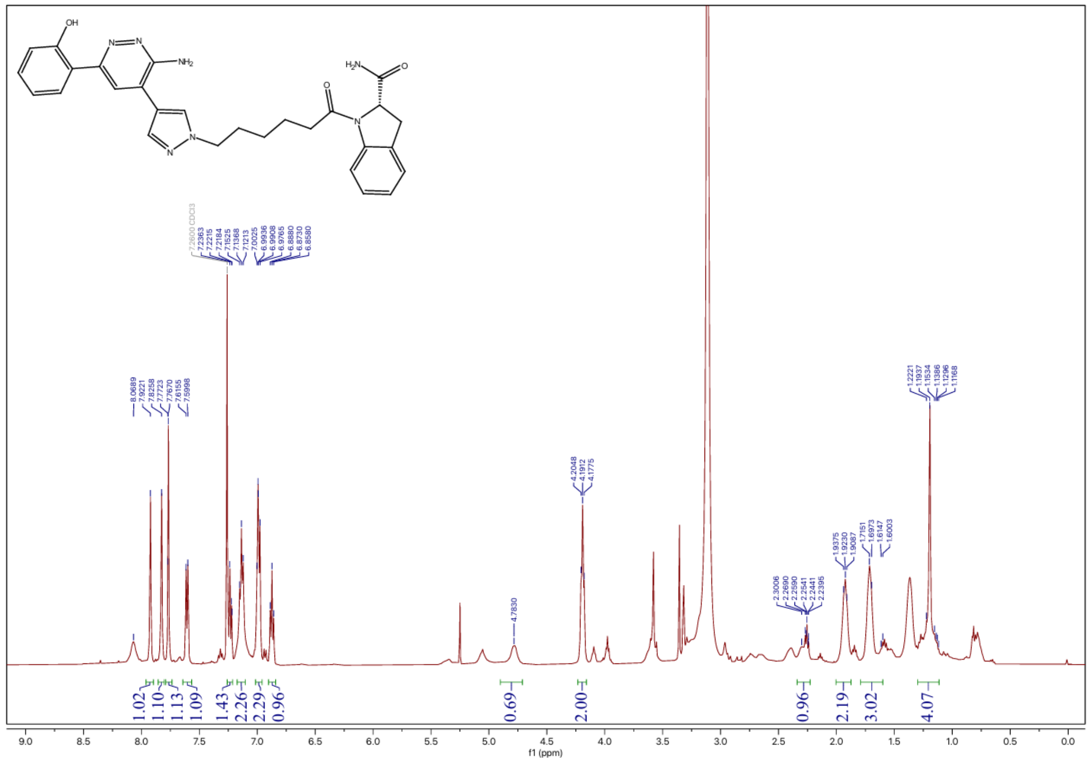


### $^{13}C$ NMR spectrum of JZ35

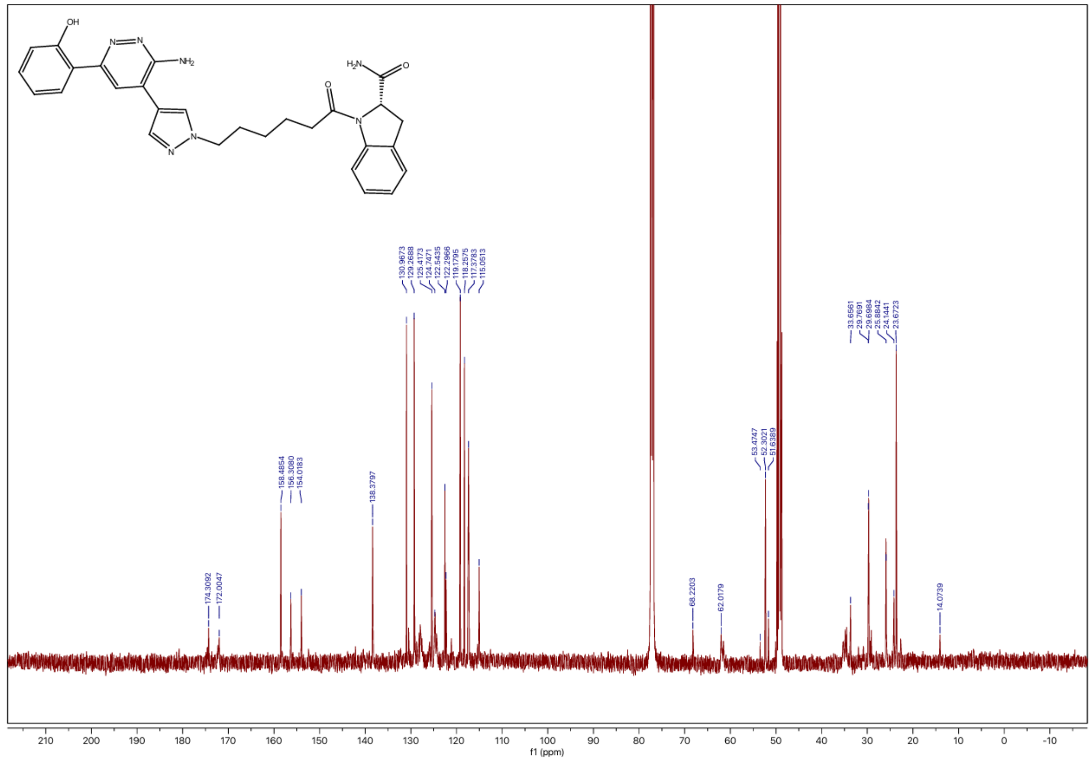

$^{1}$H NMR spectrum of JZ36

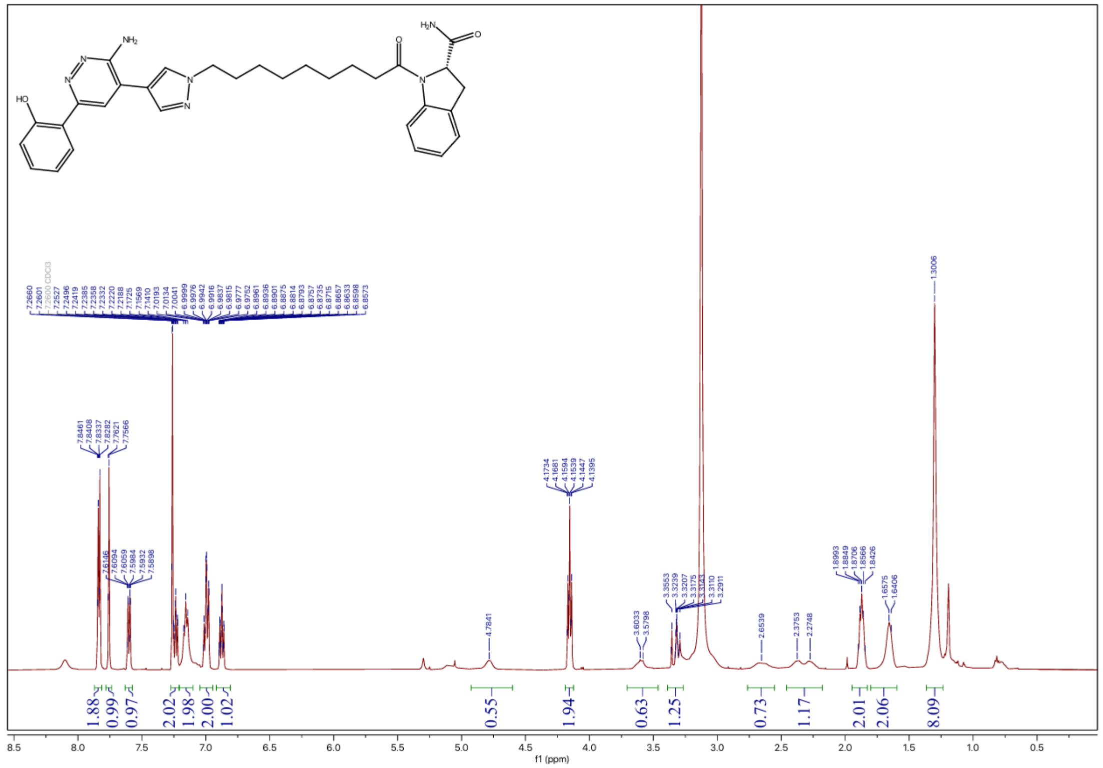

$^{13}$C NMR spectrum of JZ36

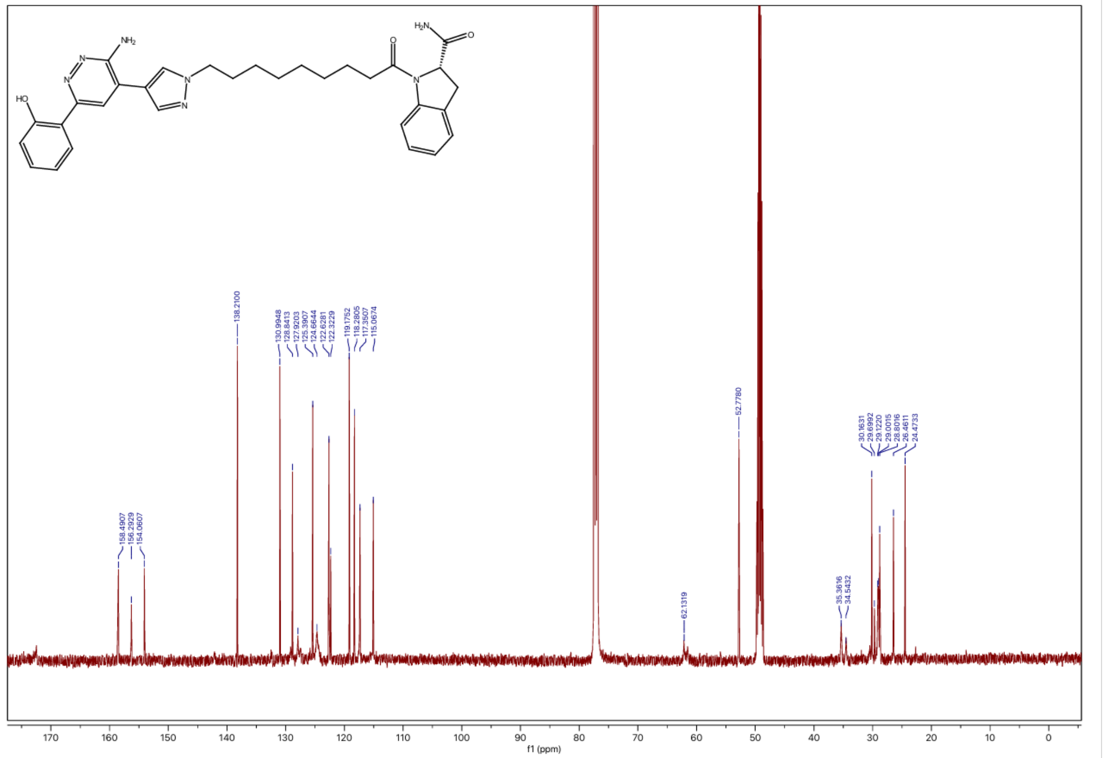

### $^1$H NMR spectrum of JZ37

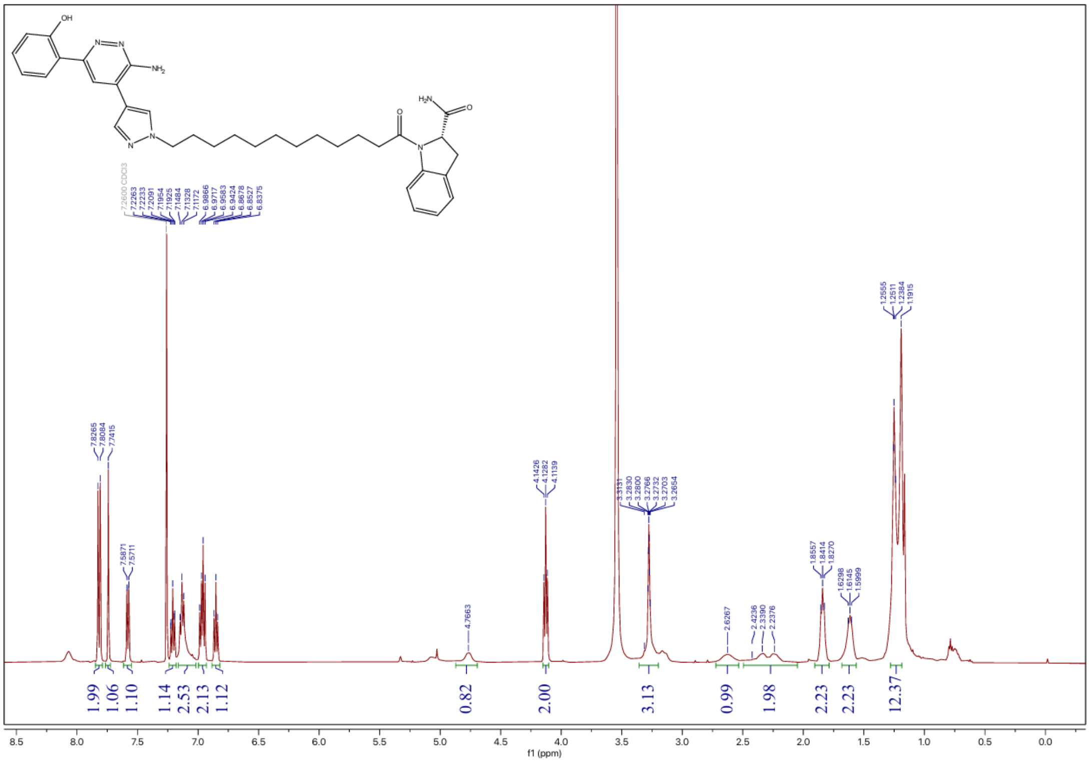


### $^{13}$C NMR spectrum of JZ37

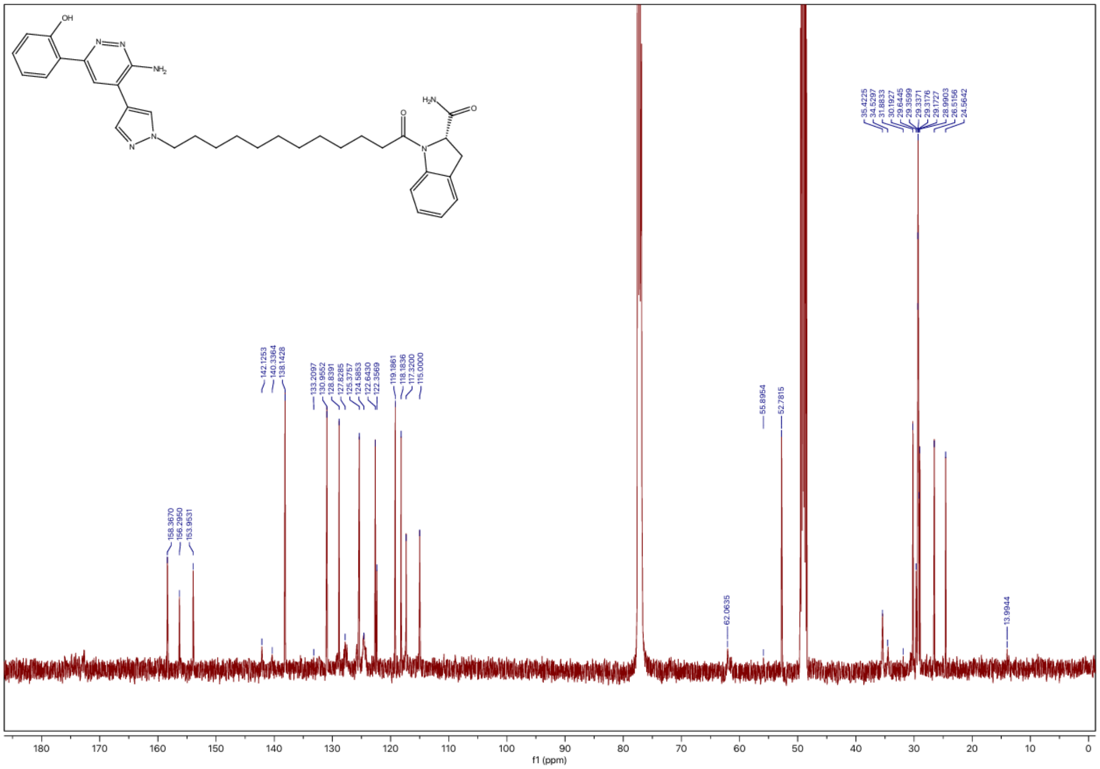
